# Homozygosity destabilizes phenotypic robustness in clonal *Drosophila mercatorum*

**DOI:** 10.64898/2026.04.02.716190

**Authors:** Ayse Kahraman, Marlene Wirth, Hussein Hammoud, Mohammad Reslan, Muhammad A. Haidar, Gabriela Djuhadi, Thomas Mathejzyk, Eric Reifenstein, Jana Balke, Max von Kleist, Gerit A. Linneweber

**Affiliations:** Institut für Biologie Abteilung Neurobiologie Fachbereich Biologie, Chemie und Pharmazie Freie Universität Berlin

## Abstract

Quantitative genetics predicts that reducing genetic variability should reduce phenotypic variability, a common principle that underlies the widespread use of inbred, isogenic, and clonal animals in biomedical research. Yet extreme homozygosity can also expose recessive deleterious alleles and detoriate developmental buffering, potentially destabilizing rather than canalizing phenotypic outcomes. We tested this conflict directly in *Drosophila mercatorum*, a facultatively parthenogenic fly species in which pronuclear duplication results in fully homozygous, clonal offspring already after a single generation. Contrary to the expectation that genetic uniformity should reduce phenotypic variability, clonal parthenogenic flies showed changes in trait means, trait-dependent changes in interindividual variation, increased fluctuating asymmetry, and, most consistently, reduced behavioral and developmental canalization across a broad set of parameters. These effects on phenotypes spanned visually guided locomotion, circadian activity, wing morphology, bristle patterning, eye anatomy, brain volume, and serotonergic neuron numbers and were associated with reduced survival under environmental stress. Inbreeding of sexual *D. mercatorum* reproduced several of these phenotypic changes, whereas a single generation of outcrossing restored phenotypic robustness and the resulting F_1_ hybrids even exceeded the wild-type controls in several traits, consistent with a hybrid vigor effect. Together, these experiments identify loss of heterozygosity, rather than clonality per se, as the primary driver of developmental instability. Our findings show that genetic uniformity achieved through homozygosity can amplify stochastic phenotypic divergence by weakening developmental canalization, and suggest that the genetic context in which experimental standardization is achieved, especially heterozygosity, matters as much as the degree of genetic uniformity itself.

## Introduction

Phenotypic variability arises from the combined effects of genetic differences, environmental influences, and stochastic developmental processes (*1*). A foundational assumption in biological research is that reducing genetic variability should reduce phenotypic variability. This widespread expectation is supported by the greater phenotypic similarity in monozygotic compared to dizygotic twins (*2, 3*) and by quantitative genetic theory (*4*). This assumption underlies the widespread use of inbred, isogenic, and clonal model organisms, which are expected to improve experimental reproducibility by minimizing the contribution of genetic variability to phenotypic outcomes (*5–8*).

Contrary to this assumption, however, the relationship between genetic and phenotypic uniformity is not straightforward (*4, 7, 9, 10*). Increased homozygosity can expose recessive deleterious alleles that were previously masked by heterozygosity, reduce developmental robustness, and promote inbreeding depression, potentially increasing phenotypic variability (*11, 12*). As a result, studies of inbred lines have produced heterogeneous outcomes: some report reduced phenotypic variation (*13–15*), others mixed results (*7, 16–18*), and yet others increased variation or reduced developmental robustness (*19, 20*). The mixed evidence suggests that two opposing forces, 1. the reduction in segregating genetic variability, which should reduce phenotypic variability, and 2. the erosion of developmental buffering capacity, which should increase it, can operate simultaneously in inbred animals, with their relative contribution depending on the organism, the trait, and the degree of homozygosity reached.

Developmental buffering, or canalization, is the tendency of developmental systems to produce consistent phenotypic outcomes despite genetic and environmental perturbations (*21*). Canalization can be assessed at multiple levels: bilateral symmetry of morphological traits, where random deviations from perfect symmetry (fluctuating asymmetry) reflect stochastic developmental noise (*22, 23*); consistency of behavior within individuals over time, where reduced repeatability indicates impaired behavioral canalization (*24, 25*); and the degree to which individuals with the same genotype converge on similar phenotypes (*9*). Heterozygosity has long been proposed to increase canalization, a view supported both theoretically and empirically (*12, 26*). Still, the precise causal relationship has been difficult to test directly, because conventional inbreeding never produces complete homozygosity, and different inbred lines vary in which recessive alleles happen to become fixed. These experimental confounds make it difficult to isolate the pure effect of extreme, complete homozygosity on canalization.

*Drosophila mercatorum* provides an unusually direct way to address this question. This species is facultative parthenogenic: rare parthenogenic reproduction occurs naturally in otherwise sexually reproducing populations, and stable obligately parthenogenic lines have been established experimentally by selecting for elevated parthenogenesis rates (*27*). Critically, parthenogenesis in *D. mercatorum* proceeds by pronuclear duplication following meiosis, which renders offspring completely homozygous after a single generation (*28*). This mechanism produces a degree of genetic uniformity that cannot be achieved even by extreme conventional inbreeding. These flies therefore permit a direct within-species comparison between heterozygous sexual populations and fully homozygous clonal populations, with the further possibility of reintroducing heterozygosity through outcrossing.

Here, we used *D. mercatorum* to test whether complete homozygosity reduces phenotypic variability or instead destabilizes developmental robustness. We compared two sets of sexually reproducing wild-type, obligately parthenogenic, facultatively parthenogenic, inbred, and outcrossed F_1_ hybrid flies across a broad panel of behavioral, physiological, and anatomical assays. Contrary to the expectation that genetic uniformity reduces phenotypic variability (*4, 29*), clonal parthenogenic flies showed across a broad set of parameters changes in trait means, trait-dependent changes in interindividual variation, increased fluctuating asymmetry, and, most consistently across all assays, reduced behavioral and developmental canalization. Inbreeding reproduced substantial parts of this phenotype, while outcrossing largely restored phenotypic robustness, identifying loss of heterozygosity as the primary driver for the phenotypic changes. These findings show that extreme homozygosity can destabilize, rather than stabilize, developmental outcomes, and suggest that the genetic context in which standardization is achieved matters as much as the degree of genetic uniformity itself.

## Materials and Methods

### Fly stocks and rearing

We used five *Drosophila mercatorum* strains. Two strains originated from Brazil and two from Hawaii: Brazil wild-type (NDSSC 15082-1521.25), Brazil obligate parthenogenic (NDSSC 15082-1521.04), Hawaii wild-type (NDSSC 15082-1521.22), and Hawaii obligate parthenogenic (NDSSC 15082-1525.05). In addition, a facultative parthenogenic strain (NDSSC 15082-1527.03) was included to control for the artificial selection procedures used to establish the parthenogenic stocks (*27*).

To restore heterozygosity in the clonal parthenogenic population, Brazil parthenogenic females (NDSSC 15082-1521.04) were crossed to Brazil wild-type males (NDSSC 15082-1521.25). The resulting F_1_ offspring were collected and analyzed for phenotypic means, interindividual variability, and canalization. This cross was designed to test whether phenotypes associated with clonality and extreme homozygosity could be mitigated by reintroducing genetic heterozygosity.

To test whether progressive loss of heterozygosity reproduces the parthenogenic phenotype, inbred lines were established from Brazil (NDSSC 15082-1521.25) and Hawaii (NDSSC 15082-1521.22) wild-type stocks by single-pair sibling mating for 5 or 10 consecutive generations without selection. After 5 and 10 generations of inbreeding, offspring were collected under standardized conditions and analyzed in the same way as the wild-type and parthenogenic flies. All genotypes used in each main and supplementary figure are listed in Supplementary Table 1.

Flies were maintained under standard *Drosophila* husbandry conditions on a cornmeal-agar diet containing, per liter, 7.5 g agar, 64 g cornmeal, 16 g yeast, 85.5 ml sugar cane syrup, 8.5 ml ethanol, 0.51 g Nipagin, and 2.5 ml propionic acid (*30*). All experiments were maintained at 25 °C, 50% relative humidity, and a 12 h:12 h light-dark cycle in a Memmert HPPeco incubator. To minimize larval crowding, experimental stocks were kept at low density with two parental females per vial, and food was changed at intervals of no more than 3 days. To reduce parental-age effects and further standardize developmental conditions, only offspring produced during the first 2 days of egg laying were used for experiments. Experimental cohorts were age-matched and reared under identical conditions.

### Buridan’s paradigm

Buridan’s paradigm experiments were performed as described previously (*31*), with minor modifications. Flies were tested at 25 °C, 6-8 days after eclosion and at least 48 h after CO₂ anesthesia. Wings were clipped before testing, as in the standard protocol; variation in clip length or shape did not measurably affect behavioral outcomes in our experiments.

The assay consisted of a circular platform (117 mm diameter) surrounded by a water-filled moat and enclosed within a uniformly illuminated white cylinder (Fig. 1A). Illumination was provided by four circular fluorescent tubes (Osram L 40W, 640C, cool white) driven by an Osram Quicktronic QT-M 1×26-42 ballast. The tubes were positioned outside a cylindrical diffuser (DeBanier, Belgium; Kalk transparent, 180 g, white) at a distance of 147.5 mm from the arena center. Unless otherwise indicated, visual cues consisted of two opposing vertical black cardboard stripes, each 30 mm wide, attached to the inner surface of the diffuser. Depending on the position of the fly, the retinal width of the stripes ranged from 8.4° to 19.6° (11.7° at the arena center) (*31*).

**Fig. 1:**
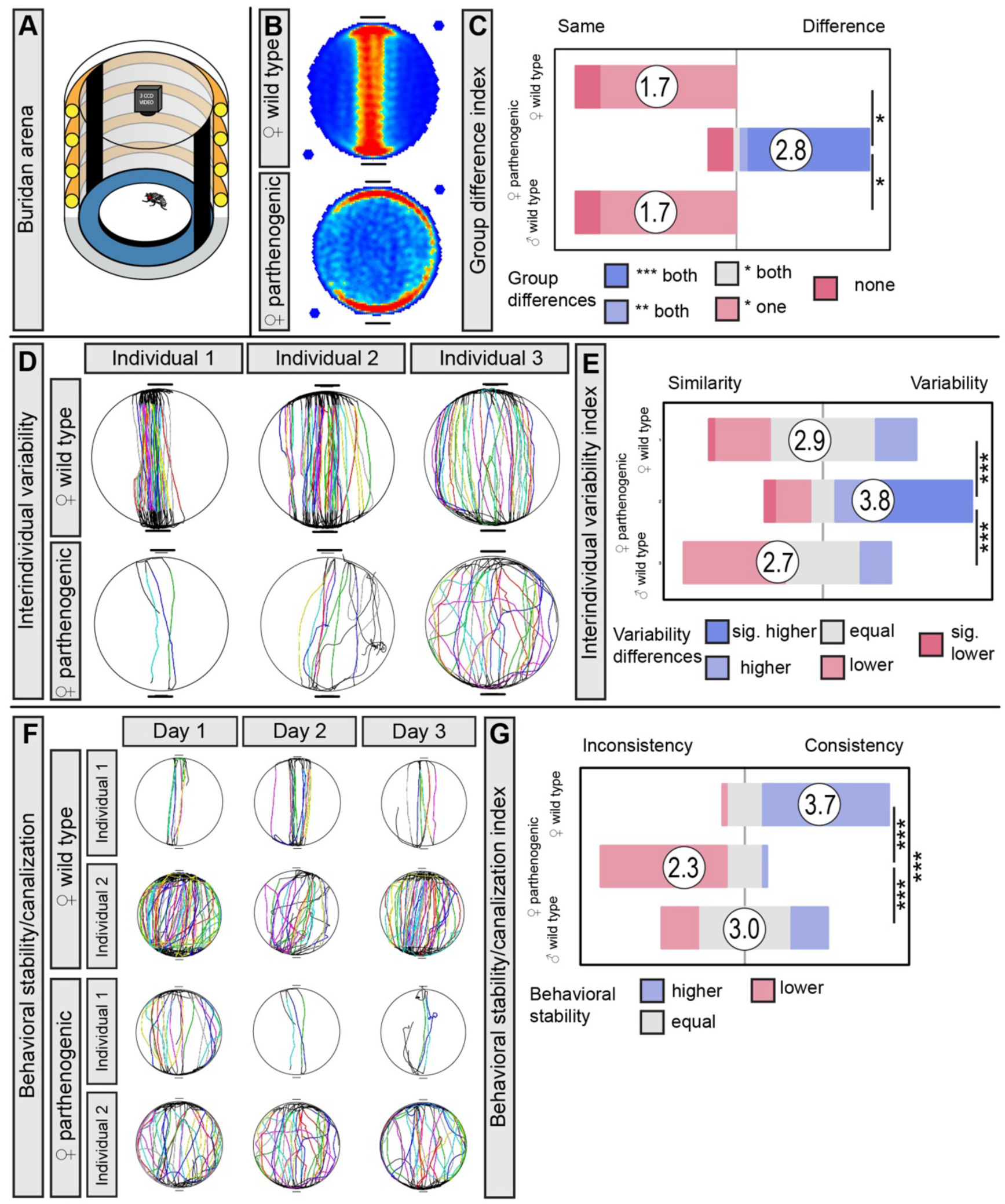
Clonal parthenogenic *Drosophila mercatorum* show altered visual-motor activity, increased interindividual behavioral variation, and reduced behavioral consistency in Buridan’s paradigm. **(A)** Schematic of Buridan’s paradigm, in which flies walk on a circular platform surrounded by water and orient relative to two opposing vertical black stripes. **(B)** Mean occupancy heatmaps show reduced stripe fixation in parthenogenic females relative to wild-type controls. **(C) Mean-difference index** summarizing genotype effects across 41 behavioral parameters shows broad changes in behavioral means in parthenogenic females relative to female and male wild-type (WT) controls. Pairwise Wilcoxon tests with Benjamini-Hochberg correction: female WT versus parthenogenic, P = 0.02; female WT versus male WT, P = 0.9; parthenogenic versus male WT, P = 0.02. **(D)** Representative individual trajectories from the same Buridan assay show greater among-individual variation in parthenogenic females than in wild-type females. **(E) Variability index** based on the median absolute deviation (MAD) across the 41 behavioral parameters shows increased interindividual behavioral variation in parthenogenic females. Pairwise Wilcoxon tests with Benjamini-Hochberg correction: female WT versus parthenogenic, P = 0.0002; female WT versus male WT, P = 0.1; parthenogenic versus male WT, P = 1.8 × 10⁻⁵. **(F)** Representative trajectories from repeated testing over 3 consecutive days show lower day-to-day behavioral consistency in parthenogenic females than in wild-type females. (**G) Behavioral consistency (canalization) index** across the 41 behavioral parameters confirms reduced behavioral stability in parthenogenic females. Pairwise Wilcoxon tests with Benjamini-Hochberg correction: female WT versus parthenogenic, P = 9.4 × 10⁻¹⁴; female WT versus male WT, P = 9.5 × 10⁻⁷; parthenogenic versus male WT, P = 9.5 × 10⁻⁷. Sample sizes: female WT, n = 33; parthenogenic, n = 31; male WT, n = 31. Asterisks denote statistical significance: p < 0.05 (*), p < 0.01 (**), p < 0.001 (***).

Fly trajectories were recorded and analyzed using CeTrAn (*31*) and custom Python scripts (*30*). From each trajectory, we extracted behavioral parameters, including total distance traveled, median speed, meander, median turning angle, absolute angle deviation to the visual stripes, the number of complete stripe-to-stripe walks, centrophobism, and activity-related measures based on speed and time thresholds, including activity time, active bouts, pause duration, and number of pauses. Absolute angle deviation was defined as the smallest angle between the walking direction vector and the vector pointing towards either stripe, such that smaller values indicate stronger stripe fixation. Walks were defined as transitions beginning within 80% of the radius from one stripe and ending within 80% of the radius from the opposite stripe. Centrophobism was quantified as the time spent in the inner versus the outer zones of equal area.

In addition, custom Python-derived metrics were calculated to capture directional and spatial aspects of visual orientation. These included directional angle deviation, stripe deviation relative to the stripe axis, horizontal deviation relative to the arena midline, and center deviation relative to the arena center. For stripe, horizontal, and center deviation, values were analyzed as directional or absolute measures. All Python-derived metrics were computed under four conditions: during locomotion, during stationary periods, across all frames combined, and across all frames with edge correction, excluding 11.78% of the arena radius. Together, all these behavioral measures yielded the 41 behavioral parameters used in the summary analyses. All 41 behavioral parameters were weighted equally in the summary indices and reanalysis frameworks.

For repeated behavioral testing (*30*), individual flies were returned to their original rearing vials after each assay and retested in subsequent sessions, across consecutive days. Each fly was treated as a single biological replicate, and day-to-day behavioral consistency was quantified within individuals before the group-level comparison. Repeated testing under these conditions does not measurably impair performance; if anything, group responses are typically slightly improved by day 3, based on observations obtained over several years.

For the one-stripe Buridan assay, only a single vertical stripe was presented as the visual stimulus. Visual orientation was quantified using horizon deviation and related measures, which report the distribution of time spent in the two halves of the arena. Flies with intact visual orientation preferentially occupied the half containing the stripe, whereas a weaker preference indicated reduced stripe-guided orientation.

### Locomotor activity and sleep measurements

Locomotor activity and sleep were measured using the *Drosophila* Activity Monitor system (DAM; Trikinetics). Flies were collected at 3 days of age and loaded individually into glass tubes (65 mm length, 7 mm inner diameter) containing food at one end and a cotton plug at the other. Tubes were inserted into LAM7 monitors and maintained at 25 °C under a 12 h:12 h light-dark cycle for 10 consecutive days. Activity was recorded in 1 min bins throughout the experiment.

The first 48 h of recording were excluded to allow acclimation to the monitor environment, and the final 24 h were excluded to standardize the analyzed time window. Activity data were analyzed using SleepMat (SleepMat2024.1). Parameters were computed in two ways: first, across 5 consecutive days to quantify overall locomotor and sleep phenotypes; and second, as 24 h repeated-measures summaries over 3 days to assess temporal behavioral consistency. Each fly was considered a biological replicate.

A total of 21 locomotor and sleep parameters were extracted and grouped into three categories: anticipation, activity, and sleep. Anticipation parameters included morning and evening anticipation indices, defined as the ratio of activity during the 3 h preceding the light transition to activity during the preceding 6 h, as well as the phase of morning and evening activity onset. Activity parameters included total activity during the light and dark phases and the full day, as well as waking activity per minute during the light and dark phases. Sleep parameters included total sleep during the light and dark phases and the full day, sleep bout number during the light and dark phases, and average sleep bout length during the light and dark phases. Sleep was defined as at least 5 consecutive minutes without infrared beam crossings (*32*).

For canalization analyses, data collected over 24 h was compared across 3 consecutive recording days for each fly. Day-to-day behavioral consistency was then quantified within individuals before group-level comparison. In the analyses, parameter-level temporal consistency was summarized using Pearson correlation coefficients after Fisher’s z-transformation. Of the 21 extracted parameters, 20 were used in repeated-day canalization analyses.

A second full data analysis was performed using the Mahalanobis distance (*33*), the MAD log-ratio (*34*), and the intraclass correlation coefficient (ICC) (*35*).

### Wing morphometry

Wings were prepared according to standard procedures (*36*). Briefly, left and right wings were dissected, mounted on microscope slides in a 1:1 mixture of glycerol and ethanol, covered with a coverslip, cleared of air bubbles, and fixed with Fixogum adhesive (Marabu). Slides were stored at 4 °C until imaging.

Wing morphology was imaged using a Leica M165 FC stereomicroscope equipped with a Leica K5C camera. Morphometric analysis was performed with the Wings4 software, followed by landmark-based shape analysis in CPReader (*36*). Together, these procedures yielded the 27 wing parameters.

Each fly was treated as one biological replicate, with left and right wings measured separately. Group differences in mean wing morphology were assessed using pairwise Wilcoxon rank-sum tests with Benjamini-Hochberg correction for multiple comparisons. Interindividual variability in wing morphology was assessed across wing traits using dispersion-based measures, including the median absolute deviation and Levene’s test.

Developmental stability and canalization were assessed from bilateral correspondence between left and right wings within individuals. Because deviations between bilateral traits provide a standard measure of fluctuating asymmetry, reduced left-right correspondence was interpreted as increased developmental noise and reduced canalization. In supplementary reanalyses, wing data were additionally evaluated using the Mahalanobis distance, the MAD log-ratio, and the intraclass correlation coefficient (ICC).

### Thorax bristle anatomy

Fly thoraxes were prepared by removing the head and abdomen and mounting the remaining thoracic cuticle on an insect pin, taking care not to damage the dorsal surface. Imaging was performed 1 day after dissection. To ensure consistent centering and orientation, the pin was secured in a micromanipulator before image acquisition. Thoraces were imaged on a Leica M165 FC stereomicroscope using z-stacks, which were subsequently flattened into extended depth-of-field images.

Flattened images were analyzed in ImageJ/Fiji (*37*) to manually extract the spatial coordinates of the bases of microchaete and macrochaete bristles. All annotations were performed by a single experimenter blinded to genotype during image analysis. Each thorax was treated as one biological replicate. The resulting two-dimensional point patterns were used to quantify bristle number, occupied area, bilateral asymmetry, and spatial regularity.

Six parameters were extracted. Total thorax bristle number was defined as the total number of annotated bristles per fly. The thorax bristle area was defined as the area enclosed by a convex hull fitted over the outermost bristles. Bilateral asymmetry was quantified as the root-mean-square point-to-point distance between left and right bristle patterns after mirroring the right side and aligning it to the left side using an iterative closest point (ICP) algorithm. A greater mismatch was interpreted as reduced developmental stability.

Local spatial regularity was assessed using three complementary measures. First, a robust spacing-irregularity metric was calculated from Voronoi tessellation by dividing the median absolute deviation of Voronoi edge lengths by their median; lower values indicate more regular spacing, whereas higher values indicate local clustering. Second, the variance-to-mean ratio (VMR) was calculated by dividing each bristle field into a 5 × 5 grid and dividing the variance of bristle counts across grid cells by the mean count within each grid cell. A VMR of 1 indicates spatial randomness, values below 1 indicate regular spacing, and values above 1 indicate clustering. Third, a regularity index was calculated as 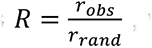, where *r_obs_* is the observed mean nearest-neighbor distance and *r_rand_* is the expected mean nearest-neighbor distance under complete spatial randomness. In two dimensions, 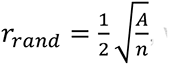, where A is the bristle-field area, and n is the number of bristles. The bristle-field area was estimated using an alpha shape with alpha = 0.01. For comparison, regularity indices were also computed for two null models: random points sampled uniformly within the empirical alpha shape, and random points sampled uniformly within a square of equal area.

### Eye morphometry

Adult compound eyes were imaged using a protocol adapted from Keesey et al. (2019) (*38*). Heads were detached from the body and immobilized by pinning through the cervical cavity with an insect pin. Each head was mounted on the confocal stage with one eye oriented towards the objective and imaged, then rotated by 180° to image the contralateral eye. Imaging was performed on a Leica SP8 white-light laser confocal microscope equipped with a 10× dry objective (NA 0.30).

Ommatidial images were analyzed in Imaris 10 (Bitplane) using semi-automated segmentation and registration. Automated segmentations were manually checked and corrected where necessary before parameter extraction. From these reconstructions, six eye parameters were extracted: number of ommatidia per compound eye (nPoints), number of edges per compound eye (nEdges), number of faces per compound eye (nFaces), mean edge length (edgeLengthMean), mean face area (faceAreaMean), and the total number of ommatidia per animal (TotalN). Together, these parameters captured both eye size and local ommatidial lattice geometry.

Both eyes were measured separately for each fly, and each fly was treated as one biological replicate. Group differences in mean morphology and interindividual variability were assessed across the six eye parameters in the summary and supplementary analyses. In addition, bilateral differences in ommatidial number were used to assess left-right asymmetry.

### Immunohistochemistry and confocal imaging

Adult *Drosophila mercatorum* brains were dissected in fresh ice-cold phosphate-buffered saline (PBS) and fixed immediately in 4% paraformaldehyde for 30 min at room temperature. Samples were washed three times in PBS and three times in PBS containing 0.4% Triton X-100 (PBST) for 10 min each, then blocked in 2% horse serum (Biozol Diagnostica; ENH2000-500) in 0.4% PBST for 1 h at room temperature.

Brains were incubated for 2 nights at 4 °C with primary antibodies against Bruchpilot (nc82; DSHB, 1:50) and serotonin (Abcam, ab66047, 1:1,000). After three 10-minute washes in PBST, samples were incubated overnight at 4 °C with donkey anti-mouse Cy3 (Jackson ImmunoResearch, 715-165-151, 1:200) and donkey anti-goat Alexa Fluor 488 (Jackson ImmunoResearch, 705-545-003, 1:200). Samples were then washed three times in PBST and once in PBS for 10 minutes each.

Brains were mounted on poly-L-lysine-coated grid slides (Electron Microscopy Sciences, 63405-01; coating with Sigma-Aldrich P1524), placed in VECTASHIELD Antifade Mounting Medium (Vector Laboratories, H-1000), sealed with clear nail polish and a coverslip, and stored at 4 °C until imaging.

All immunostained samples were imaged on a Leica TCS SP8 X White Light Laser confocal microscope equipped with a 20× glycerol-immersion objective (NA 0.75).

### Brain neuropile volumetry

Brain neuropile anatomy was quantified in Imaris 10 (Bitplane) from nc82/anti-Brp labeled confocal stacks. Volumes of the left and right optic lobes and of the left and right halves of the central brain were segmented using the *Surface* object-detection algorithm and extracted from the Imaris statistics output. Automated segmentations were manually checked and corrected where necessary before parameter extraction. Left and right measurements were extracted separately and compared within individuals for asymmetry analyses. Each fly was considered a biological replicate. These measurements were used to quantify mean brain anatomy, interindividual variability, and left-right asymmetry across the neuropile parameters.

### Serotonergic neuron quantification

Serotonergic neurons were quantified in Imaris 10 (Bitplane) using the *Spots* object-detection algorithm. Automated detections were manually checked and corrected where necessary before parameter extraction. Left and right measurements were extracted separately and compared within individuals for asymmetry analyses. Each fly was considered a biological replicate. The resulting measurements were used to assess mean serotonergic-neuron number, interindividual variation, and left-right correspondence in downstream analyses.

### Electroretinogram (ERG)

Electroretinogram (ERG) recordings were performed according to standard procedures (*39*). Flies were immobilized by attaching the wings to a microscope slide with craft glue (UHU), allowing stable positioning during recording. A glass microelectrode was inserted into the compound eye, and a reference electrode was placed in the thorax. Each fly was recorded only once to avoid repeated-measure artifacts.

Signals were acquired using Axoscope 10.6 (Molecular Devices). Light stimuli of 1 ms duration were delivered at 1% and 10% of maximum output. For each stimulus intensity, five responses were recorded per fly. From the ERG traces, two response components were quantified: the on-transient amplitude and the sustained depolarization. Responses were averaged per fly and per stimulus condition before statistical analysis.

ERG measurements were used to assess visual responsiveness independently of behavioral orientation in Buridan’s paradigm. In the reported analyses, group comparisons were performed for on-transient amplitude and sustained depolarization at both 1% and 10% stimulus intensity.

### Statistical analysis

All statistical analyses were performed in R. Unless otherwise stated, each fly was treated as one biological replicate. Group differences were assessed using pairwise Wilcoxon rank-sum tests, and homogeneity of variance was assessed using pairwise Levene’s tests. Whenever multiple comparisons were performed, P values were corrected using the Benjamini-Hochberg procedure. Correlations were quantified using the Pearson product-moment correlation coefficient. For repeated-day analyses, Pearson correlation coefficients were averaged after Fisher’s z transformation, followed by back-transformation.

### Categorical summary indices

To summarize large multivariate behavioral and anatomical datasets at the experiment level, parameter-wise results were converted into categorical scores and visualized as Likert summary indices. Full parameter-level tables underlying all categorizations are provided in Supplementary Table 2. Depending on the experiment and the number of comparison groups, categories were assigned slightly differently. In all cases, however, the indices were designed to summarize three properties across a set of parameters: mean phenotype, interindividual variability, and canalization. Pairwise comparisons between index distributions were performed using Wilcoxon rank-sum tests with Benjamini-Hochberg correction.

For mean indices, higher category values corresponded to stronger parameter-wise divergence from the reference or control genotype. For variability indices, category values indicated whether a genotype was more or less variable than comparison groups, based on median absolute deviation (MAD) or related dispersion metrics. For canalization indices, category values indicated whether a genotype showed greater or lower behavioral repeatability or anatomical bilateral correspondence than the comparison groups. Exact category definitions for each assay are given in Supplementary Table 2.

### Multivariate reanalysis of mean differences

As a continuous alternative to the categorical mean index, multivariate phenotypic divergence between genotypes was quantified using a robust Mahalanobis distance relative to a reference/control genotype (*33*). Individual-level data were centered and, where appropriate, projected onto principal component axes to reduce dimensionality and collinearity. A robust covariance matrix was estimated using the Minimum Covariance Determinant method, and Mahalanobis distances between genotype mean vectors and the reference/control genotype were calculated in this multivariate space. Positive values indicate greater divergence from the reference/control genotype, whereas values near zero indicate high similarity. Uncertainty was estimated by nonparametric bootstrap resampling, and a permutation test with random reassignment of genotype labels assessed statistical significance.

### Multivariate reanalysis of interindividual variability

As a continuous alternative to the categorical variability index, phenotypic variability was quantified using a robust median-based log-variability metric (MAD log-ratio) relative to a control/reference genotype (*34*). For each parameter, within-genotype dispersion was estimated using the median absolute deviation. Log-ratios of MAD values were computed for each genotype relative to the reference/control genotype, and the mean across all parameters was used as a summary measure of relative variability. Positive values indicate greater variability than the reference genotype, whereas negative values indicate lower variability. Uncertainty was estimated by bootstrap resampling, and significance was assessed by a permutation test with genotype-label reassignment.

### Quantification of canalization and repeatability

As a continuous alternative to the categorical canalization index, behavioral or anatomical repeatability was quantified using the intraclass correlation coefficient (ICC) (*35*). For repeated behavioral measurements, ICC estimates the proportion of variance attributable to stable between-individual differences across days. For bilateral anatomical measurements, ICC estimates the degree of left-right correspondence within individuals. Higher ICC values indicate greater behavioral consistency or developmental stability, whereas lower ICC values indicate reduced canalization. Group differences in ICC-based canalization were evaluated by permutation testing with Benjamini-Hochberg correction.

## Results

*Drosophila mercatorum* undergoes parthenogenesis by pronuclear duplication following meiosis, producing fully homozygous and clonal offspring after a single generation, a degree of genetic uniformity not ever achievable by conventional inbreeding (*28*). To test whether extreme homozygosity reduces interindividual phenotypic variability or instead destabilizes developmental robustness, we compared eight genotypes that differ in heterozygosity and interindividual genetic variation: sexually reproducing wild-type flies from two geographic origins (Brazil and Hawaii), obligately parthenogenic lines derived from the same geographic origins, a facultatively parthenogenic line, inbred derivatives of wild-type lines, and F_1_ hybrid offspring generated by crossing parthenogenic females with wild-type males. Across all comparisons, we quantified three properties: the mean phenotype, interindividual variation, and canalization, measured as within-individual behavioral repeatability or bilateral anatomical correspondence (fluctuating asymmetry).

### 1. Clonal flies show altered visual behavior, increased interindividual variation, and impaired behavioral canalization

We first quantified visually guided locomotion behavior through Buridan’s paradigm, in which individual flies walk freely on a circular platform and orient themselves relative to two opposing vertical stripes (Fig. 1A) (*30, 31*). Compared with wild-type females and males, parthenogenic females (derived from Brazil) showed markedly reduced stripe fixation and broad changes in mean behavior across all 41 quantified parameters (Fig. 1B,C; Supplementary Fig. S1A,E,F). The same pattern was observed in independently derived Hawaiian lines and confirmed by a Mahalanobis-distance analysis (Supplementary Figs. S1 and S2).

To determine whether the weaker stripe fixation of parthenogenetic flies reflected impaired visual processing or a locomotor deficit, we tested them in a one-stripe version of Buridan’s paradigm. All groups, including parthenogenic flies, maintained normal locomotion behavior and a clear preference for the stripe (Supplementary Fig. S3). Electroretinogram recordings showed normal depolarization amplitudes, whereas the on-transient was moderately reduced in parthenogenic flies (Supplementary Fig. S4). These results argue against blindness and instead suggest a more subtle impairment in visual processing or sensorimotor transformation in the parthenogenic flies.

Contrary to the assumption that genetic uniformity should reduce phenotypic variability (*4, 5, 13, 40*), clonal flies showed greater interindividual variation than controls across the 41 behavioral parameters, as confirmed by both the MAD-based index and MAD log-ratio analysis (Fig. 1D,E; Supplementary Figs. S1 and S2). The same overall pattern was observed in an independent Hawaii-derived dataset (Supplementary Figs. S1 and S2). Thus, in Buridan’s paradigm, genetically clonal populations displayed greater, not lower, interindividual behavioral variation.

To determine whether the elevated variation found in parthenogenic flies reflected stable interindividual differences (*41, 42*) or reduced within-individual consistency (*43*), we retested each fly on three consecutive days. Parthenogenic flies showed reduced day-to-day behavioral consistency (Fig. 1F,G; Supplementary Figs. S2 and S5), indicating that clonal flies are less predictable over time. Critically, reduced temporal consistency did not abolish consistent interindividual differences (*30*): parthenogenic flies remained individually distinct despite weaker behavioral canalization, demonstrating that behavioral individuality persists even when developmental buffering is impaired.

### 2. Circadian and sleep behavior also shows altered trait means and reduced canalization, but not uniformly increased variability

To test whether the changes in trait means, variation, and behavioral consistency observed through Buridan’s paradigm generalized across behavioral contexts, we monitored individual locomotor activity and sleep in *D. mercatorum* over two weeks using the Trikinetics activity monitors. Parthenogenic flies again differed from sexually reproducing controls in mean circadian activity and sleep phenotypes across 21 parameters (Fig. 2A,B; Supplementary Figs. S6 and S7). In contrast to the Buridan assay, however, interindividual variation was not significantly increased in the parthenogenic flies (Fig. 2C,D). Longitudinal analysis nevertheless revealed a marked reduction in within-individual consistency over time (Fig. 2E; Supplementary Fig. S6). Thus, across two independent behavioral paradigms (Buridan’s paradigm and Trikinetics activity monitors), clonality robustly changes mean behavior and reduces behavioral consistency, but does not uniformly increase interindividual variation.

**Fig. 2:**
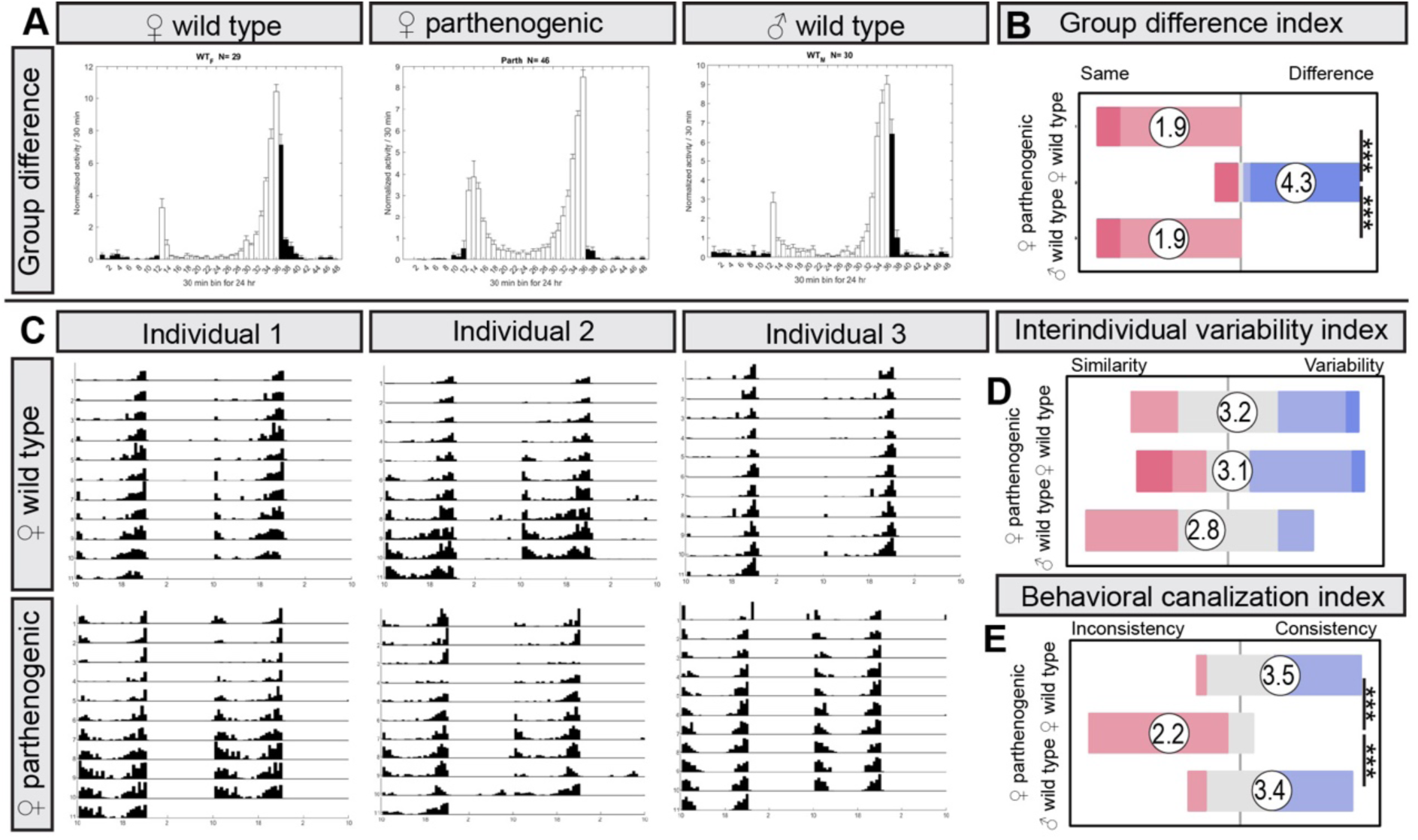
Parthenogenic *Drosophila mercatorum* show altered circadian locomotor output and reduced temporal behavioral canalization without increased overall interindividual variation. **(A)** Mean locomotor activity profiles across the light-dark cycle for female wild-type, parthenogenic female, and male wild-type flies reveal altered daily activity patterns in parthenogenic flies. **(B) Mean-difference index** across 21 locomotor activity and sleep parameters shows broad changes in behavioral means in parthenogenic females relative to both wild-type control groups. Pairwise Wilcoxon tests with Benjamini-Hochberg correction: female WT versus parthenogenic, P = 2.8 × 10⁻⁵; female WT versus male WT, P = 1; parthenogenic versus male WT, P = 2.8 × 10⁻⁵. **(C)** Representative individual actograms from wild-type and parthenogenic females across repeated recording days show reduced temporal stability of activity patterns in parthenogenic flies. **(D) Variability index** based on the median absolute deviation (MAD) across the 21 activity and sleep parameters indicates similar interindividual variation across groups. Pairwise Wilcoxon tests with Benjamini-Hochberg correction: female WT versus parthenogenic, P = 0.9; female WT versus male WT, P = 0.1; parthenogenic versus male WT, P = 0.2. **(E) Behavioral canalization index** across 20 repeatedly measured parameters shows reduced day-to-day behavioral consistency in parthenogenic females. Pairwise Wilcoxon tests with Benjamini-Hochberg correction: female WT versus parthenogenic, P = 8.7 × 10⁻⁷; female WT versus male WT, P = 0.7; parthenogenic versus male WT, P = 4.1 × 10⁻⁶. Sample sizes: female WT, n = 33; parthenogenic, n = 32; male WT, n = 32. Asterisks denote statistical significance: p < 0.05 (*), p < 0.01 (**), p < 0.001 (***).

### 3. Clonality alters multiple anatomical traits and broadly reduces developmental robustness

We next asked whether the phenotypic effects of clonality were specific to behavior or extended to morphology. We first examined the wings, a classic system for studying size variation, fluctuating asymmetry, and developmental canalization (*15, 17, 22, 44*). Contrary to the expectation that inbreeding would reduce size (*45*), parthenogenic female flies had significantly larger wings than both sexually reproducing control groups, with consistent changes across all 27 wing parameters (Fig. 3A; Supplementary Figs. S8 and S9).

**Fig. 3:**
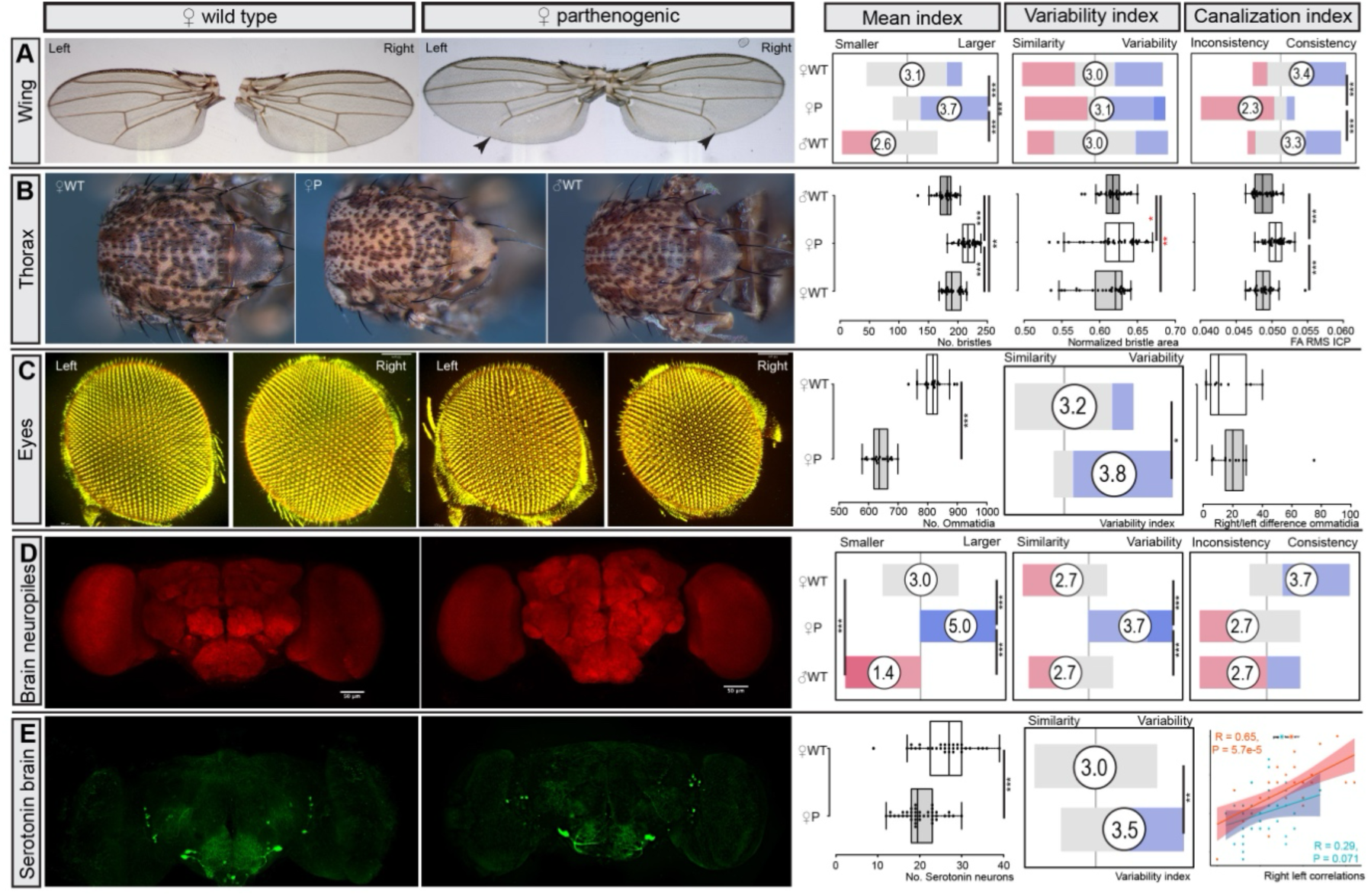
Parthenogenic *Drosophila mercatorum* show altered anatomy, increased developmental instability, and reduced canalization across multiple morphological traits. **(A)** Representative wing images from wild-type and parthenogenic females. A shortened L7 wing vein is consistently observed in parthenogenic flies and not in controls. The **mean-difference index** across 27 wing parameters shows broad changes in wing morphology in parthenogenic females compared with wild-type female and male controls. Pairwise Wilcoxon tests with Benjamini-Hochberg correction: female WT versus parthenogenic, P = 4.5 × 10⁻⁵; female WT versus male WT, P = 1.0 × 10⁻⁴; parthenogenic versus male WT, P = 2.9 × 10⁻⁸. The **variability index** based on the median absolute deviation (MAD indicates no overall increase in interindividual wing variation in parthenogenic females. Pairwise Wilcoxon tests with Benjamini-Hochberg correction: female WT versus parthenogenic, P = 0.8; female WT versus male WT, P = 1; parthenogenic versus male WT, P = 0.7. The **canalization index** based on left-right wing correspondence indicates reduced developmental stability in parthenogenic females. Pairwise Wilcoxon tests with Benjamini-Hochberg correction: female WT versus parthenogenic, P = 2.8 × 10⁻⁶; female WT versus male WT, P = 0.4; parthenogenic versus male WT, P = 1.3 × 10⁻⁶. Sample sizes: female WT, n = 29; parthenogenic, n = 31; male WT, n = 33. **(B)** Representative thorax bristle patterns from wild-type females, parthenogenic females, and wild-type males. Parthenogenic females have more thoracic bristles than both wild-type groups, whereas wild-type males and females show sexual dimorphism in bristle number. Pairwise Wilcoxon tests with Benjamini-Hochberg correction: female WT versus parthenogenic, P = 1.1 × 10⁻⁷; female WT versus male WT, P = 0.003; parthenogenic versus male WT, P = 1.2 × 10⁻¹¹. Variation in bristle number does not differ consistently between parthenogenic and wild-type females. Pairwise Levene’s tests: female WT versus parthenogenic, P = 0.45; female WT versus male WT, P = 0.03; parthenogenic versus male WT, P = 0.001. A fluctuating-asymmetry metric based on the root-mean-square point-to-point mismatch between mirrored left- and right-bristle patterns shows reduced canalization in parthenogenic females. Pairwise Wilcoxon tests with Benjamini-Hochberg correction: female WT versus parthenogenic, P = 7.2 × 10⁻⁶; female WT versus male WT, P = 0.67; parthenogenic versus male WT, P = 2.6 × 10⁻⁵. Sample sizes: female WT, n = 32; parthenogenic, n = 38; male WT, n = 35. **(C)** Representative three-dimensional reconstructions of compound eyes from wild-type and parthenogenic females. Parthenogenic females have smaller eyes with fewer ommatidia than wild-type females. Pairwise Wilcoxon test: P = 6.8 × 10⁻⁸. Across six eye parameters, parthenogenic flies show increased interindividual variation. Pairwise Wilcoxon test: P = 0.03. Left-right differences in ommatidial number are also increased, although not significantly. Pairwise Wilcoxon test: P = 0.3. Sample sizes: female WT, n = 10; parthenogenic, n = 10. **(D)** Representative three-dimensional reconstructions of brain neuropiles labeled with nc82 from wild-type and parthenogenic females. Parthenogenic flies have larger brains than wild-type flies. A **mean-difference index** across seven neuropile parameters shows broad changes in brain anatomy in parthenogenic females relative to both wild-type groups. Pairwise Wilcoxon tests with Benjamini-Hochberg correction: female WT versus parthenogenic, P = 0.0004; female WT versus male WT, P = 0.0008; parthenogenic versus male WT, P = 0.0008. A MAD-based **variability index** indicates increased interindividual variation in parthenogenic females. Pairwise Wilcoxon tests with Benjamini-Hochberg correction: female WT versus parthenogenic, P = 0.001; female WT versus male WT, P = 0.66; parthenogenic versus male WT, P = 0.001. A **canalization index** based on left-right asymmetry across three bilateral parameters shows no significant group difference. Pairwise Wilcoxon tests with Benjamini-Hochberg correction: female WT versus parthenogenic, P = 0.16; female WT versus male WT, P = 0.35; parthenogenic versus male WT, P = 1. Sample sizes: female WT, n = 22; parthenogenic, n = 23; male WT, n = 16. **(E)** Representative three-dimensional reconstructions of serotonergic neurons from wild-type and parthenogenic females. Parthenogenic females show fewer serotonergic neurons and greater left-right asymmetry than wild-type females. Mean neuron number is reduced in parthenogenic females. Pairwise Wilcoxon test: P = 1.9 × 10⁻⁵. Across 16 serotonergic parameters, parthenogenic flies show increased variation. Pairwise Wilcoxon test: P = 0.0014. Left-right correspondence is reduced in parthenogenic flies, indicating reduced canalization. Wild-type female: R = 0.65, P = 5.7 × 10⁻⁵. Parthenogenic: R = 0.29, P = 0.071. Sample sizes: female WT, n = 32; parthenogenic, n = 40. Asterisks denote statistical significance: p < 0.05 (*), p < 0.01 (**), p < 0.001 (***).

Changes in wing variability were not uniform. Parthenogenic flies showed increased variation in some traits and reduced variation in others, including elevated variation in specific wing vein lengths such as L7 and L17. More strikingly, fluctuating asymmetry was increased, most prominently in the length of wing vein L7, resulting in reduced canalization across wing traits (Fig. 3A; Supplementary Figs. S8 and S9). An independently derived Hawaiian parthenogenic stock confirmed the overall pattern, although the effect on variation differed across strains (Supplementary Fig. S8D-H and S9D-F). These results indicate that extreme loss of heterozygosity does not simply increase or decrease morphological variation globally, but it consistently impairs developmental stability.

We next asked whether these patterns of phenotypical changes continue in thoracic bristles, another sensitive readout of developmental precision (*25, 46*). Parthenogenic flies showed reduced thoracic pigmentation and slightly larger thoraces than wild-type controls (Fig. 3B). Bristle counts revealed a modest increase in bristle number in parthenogenic flies relative to wild-type females. Although interindividual variation in bristle number did not change consistently in one direction, canalization was again reduced, as indicated by less regular bristle spacing and increased left-right mismatch (Fig. 3B; Supplementary Fig. S10).

Compound-eye morphology showed a distinct pattern. Despite their generally larger body size, parthenogenic flies had smaller eyes, due to significantly fewer ommatidia (Fig. 3C). In contrast to the wing phenotype, interindividual variation in eye morphology was increased, and asymmetry between the left and right eyes also tended to be more prominent (Fig. 3C, Supplementary Fig. S11). Thus, although changes in variation differed across traits, our measurements of eye morphology indicate reduced developmental robustness in parthenogenetic flies. Altogether, across wing, bristle, and eye morphology, clonality and homozygosity altered mean and variation in trait-specific ways, while most consistently reducing developmental robustness and canalization.

### 4. Instability of neural anatomy mirrors that of external characters

To determine whether the effects of clonality and homozygosity on trait means, variation, and canalization extended to the nervous system, we quantified neuropil anatomy and serotonergic neuron numbers. Parthenogenic flies, as expected from previous results, had larger brains than wild-type controls, with broad changes across neuropile parameters and increased interindividual variation (Fig. 3D; Supplementary Fig. S12). Large-scale left-right asymmetry of major brain compartments was less strongly affected, suggesting that clonality disrupts neural size regulation more than gross bilateral organization.

We next examined serotonergic neurons, whose neuromodulatory function has been implicated in behavioral variability (*47*). Parthenogenic flies had fewer serotonergic neurons than controls and showed increased variation and greater left-right asymmetry (Fig. 3E; Supplementary Fig. S13). Thus, clonality not only affects external morphological characters but also impacts neural cell populations that may directly contribute to altered behavioral individuality (*47, 48*).

Finally, we asked whether reductions in fitness accompanied reduced developmental robustness. Survival assays under environmental challenges showed that parthenogenic flies had lower survival than sexually reproducing controls across multiple environmental conditions (Supplementary Fig. S14). This deficit was especially pronounced at a stress temperature (29°C), where no parthenogenic adults emerged. Thus, the loss of canalization in parthenogenic flies is associated not only with altered anatomy and increased phenotypic variation but also with reduced tolerance to environmental stress.

### 5. Loss of heterozygosity, rather than clonality, drives the major phenotypic effects of parthenogenesis

The behavioral and anatomical effects described so far could, in principle, reflect clonality itself, loss of heterozygosity, or confounding effects specific to the parthenogenic stocks. We performed four experiments to distinguish among these possibilities and to determine whether loss of heterozygosity, rather than clonality itself, was responsible for changes in trait means, interindividual variation, and canalization.

We first tested whether the increased variation in our parthenogenic populations could simply reflect the contribution of a large number of parthenogenic mothers, given the low fertility of the parthenogenic stocks. To address this, we analyzed behavioral and wing-morphological traits in offspring derived from a single parthenogenic mother. Measurements in these flies closely matched those from the pooled parthenogenic populations in both Buridan’s paradigm behavior and wing anatomy (Supplementary Figs. S15 and S16), indicating that high numbers of mothers cannot account for the elevated variation measured in the parthenogenic lines.

We next sought to separate the effects of homozygosity from those of clonality itself by examining a facultative parthenogenic strain with elevated parthenogenic potential (*27, 49*). In this strain, reproduction is typically sexual, so parthenogenic offspring do not form a clonal population and can retain substantial interindividual genetic variation. Nevertheless, these facultatively parthenogenic offspring closely resembled the clonal obligately parthenogenic lines in both behavior and wing morphology (Supplementary Figs. S17 and S18), including similar levels of interindividual variation. These results support the idea that reduced heterozygosity, rather than clonality, is a major driver of the observed changes in trait means, variation, and canalization.

To further test this interpretation and to exclude maternal effects, we crossed parthenogenic females with wild-type males and analyzed the sexually produced F_1_ hybrid offspring. The F_1_ hybrid offspring largely resembled wild-type flies in both behavior and wing morphology (Supplementary Figs. S19-S21), including restoration of traits that were strongly disrupted in the parthenogenic lines, such as the asymmetric L7 wing-vein phenotype. In several canalization measures, the rescue even exceeded the wild-type state (Supplementary Fig. S21C,E,H, and J), consistent with a pronounced hybrid-vigor effect (*4, 29, 50*). This interpretation is further supported by the slightly increased wing and body size observed in one F_1_ hybrid group.

We then directly tested whether reduced heterozygosity is sufficient to reproduce the clonal phenotype by generating inbred lines without selection. Even relatively mild inbreeding over five generations modified mean behavioral phenotypes away from the wild-type state (Fig. 4A; Supplementary Figs. S22-S24). Effects on interindividual behavioral variation were already evident at this stage, although their direction depended on sex and population background. In contrast, reductions in behavioral consistency were more reproducible, with inbred flies often showing intermediate canalization between wild-type and parthenogenic animals. After ten generations of inbreeding, these effects were often amplified, though in some cases partially reversed, indicating that progressive homozygosity has complex, nonlinear effects on behavioral means, variance structure, and repeatability.

**Figure 4:**
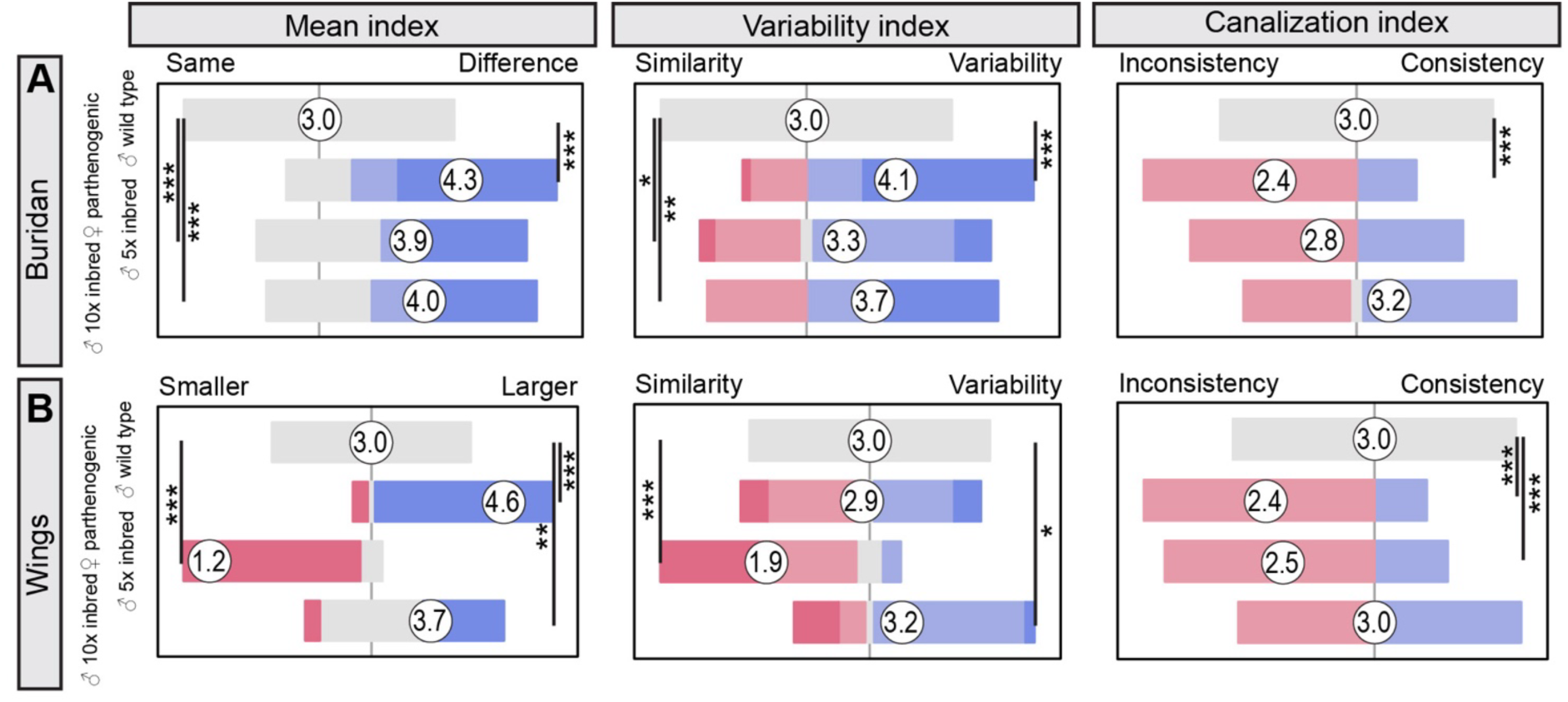
Inbreeding recapitulates the effects of clonality on wing morphology and developmental canalization in Drosophila mercatorum. **(A) Mean-difference index** across 41 Buridan behavioral parameters for Brazil-derived males shows that clonal parthenogenic flies differ strongly from wild-type controls, whereas 5× and 10× inbred wild-type l **(B)** ines show weaker effects. Pairwise Wilcoxon tests with Benjamini-Hochberg correction: male WT versus parthenogenic, P = 0.02; male WT versus 5× inbred, P = 0.61; male WT versus 10× inbred, P = 0.70. The behavioral **variability index** indicates no significant difference among groups. Pairwise Wilcoxon tests with Benjamini-Hochberg correction: male WT versus parthenogenic, P = 0.59; male WT versus 5× inbred, P = 0.44; male WT versus 10× inbred, P = 0.21. The behavioral **canalization index** shows reduced behavioral consistency in clonal parthenogenic flies and, to a lesser extent, in 10× inbred flies. Pairwise Wilcoxon tests with Benjamini-Hochberg correction: male WT versus parthenogenic, P = 2.5 × 10⁻⁶; male WT versus 5× inbred, P = 0.08; male WT versus 10× inbred, P = 0.005. Sample sizes: male WT, n = 33; parthenogenic, n = 31; 5× inbred, n = 29; 10× inbred, n = 29. **(C) Mean-difference index** across 27 wing parameters for Brazil males shows broad changes in wing morphology in parthenogenic flies and in both inbred lines relative to wild-type controls. Mild inbreeding (5×) changes wings towards smaller values, whereas 10× inbreeding changes them back towards the enlarged phenotype seen in parthenogenic flies. Pairwise Wilcoxon tests with Benjamini-Hochberg correction: female WT versus parthenogenic, P = 4.5 × 10⁻⁵; female WT versus 5× inbred, P = 2.6 × 10⁻⁸; female WT versus 10× inbred, P = 8.7 × 10⁻⁴. The **variability index** based on the median absolute deviation (MAD) indicates no overall difference in interindividual wing variation. Pairwise Wilcoxon tests with Benjamini-Hochberg correction: female WT versus parthenogenic, P = 0.44; female WT versus 5× inbred, P = 0.44; female WT versus 10× inbred, P = 0.21. The **canalization index**, based on left-right correspondence across wing traits, shows reduced developmental stability in parthenogenic flies and in 10× inbred flies. Pairwise Wilcoxon tests with Benjamini-Hochberg correction: female WT versus parthenogenic, P = 2.5 × 10⁻⁶; female WT versus 5× inbred, P = 0.08; female WT versus 10× inbred, P = 0.005. Sample sizes: female WT, n = 29; parthenogenic, n = 31; 5× inbred, n = 27; 10× inbred, n = 31. Asterisks denote statistical significance: p < 0.05 (*), p < 0.01 (**), p < 0.001 (***).

Wing anatomy showed a similarly partial recapitulation of parthenogenic phenotypes (Fig. 4B; Supplementary Figs. S25-S27). Inbred lines differed from wild-type in mean wing morphology, with mild inbreeding often modifying wings toward smaller values and more extensive inbreeding changed them toward the enlarged phenotype observed in parthenogenic flies. As in the behavioral assays, effects on interindividual variation were heterogeneous across sexes, populations, and traits. By contrast, changes in developmental stability were more consistent: parthenogenic flies and some inbred lines showed reduced bilateral correspondence across wing traits, indicating impaired canalization. Thus, loss of heterozygosity is sufficient to reproduce substantial parts of the clonal phenotype, particularly changes in mean trait values and reduced developmental stability, even though effects on variation remain trait- and background-dependent.

Taken together, these experiments identify reduced heterozygosity as a major driver of the phenotypic effects observed in parthenogenic flies. They further show that the consequences of extreme homozygosity are not limited to changes in trait means, but extend to interindividual variability and, most consistently, to reduced behavioral and developmental canalization.

## Discussion

A central assumption in experimental biological research is that reducing genetic variability should reduce phenotypic variability (*4, 5, 7, 13*). Our results challenge this assumption when the reduction in genetic variability is achieved through extreme homozygosity.

In *Drosophila mercatorum*, clonal parthenogenic populations did not become phenotypically uniform. Instead, they showed broad changes in trait means, trait-dependent changes in interindividual variation, increased fluctuating asymmetry, and, most consistently, reduced behavioral and developmental canalization across multiple independent assays compared with wild-type controls. These effects were partially reproduced by inbreeding and largely reversed by outcrossing, identifying loss of heterozygosity as the principal driver of the phenotype changes.

The most reproducible consequence of extreme homozygosity in our experiments was not a change in variation per se, but a loss of phenotypic robustness. Across multiple behaviors and morphological characters, parthenogenic flies repeatedly showed reduced within-individual repeatability, increased bilateral asymmetry, and weaker left-right correspondence, all indicators of impaired canalization. By contrast, changes in interindividual variation were more trait-dependent: in most assays, variation increased, in others it remained unchanged, and in a few cases it even decreased. This distinction matters. Quantitative genetics predicts that reducing genetic variability should reduce phenotypic variability (*4*), and our data does not violate this principle. What our results show, however, is that when homozygosity becomes extreme, destabilization of developmental buffering can offset or outweigh the variability-reducing effects of genetic uniformity. This tension likely explains why studies of inbred lines have historically produced mixed outcomes across traits and systems (*7, 13–20*). In *D. mercatorum*, complete homozygosity resulting from pronuclear duplication makes this destabilizing component of the phenotype especially visible.

The genotypes compared in this study point to heterozygosity, rather than clonality, as a phenotypic stabilizing factor. Parthenogenic offspring derived from a single mother closely resembled the pooled parthenogenic lines, ruling out maternal diversity as an explanation for elevated variation. Facultative parthenogenic offspring, homozygous but not forming a clonal population, reproduced all major parthenogenic phenotypes, showing that clonality is not required. Most compellingly, outcrossing parthenogenic females to wild-type males largely restored wild-type behavior and morphology, and in these hybrids several canalization measures exceeded the wild-type state, consistent with a hybrid vigor effect (*50*). Together, these experiments indicate that heterozygosity is a stabilizing force, likely because it masks recessive deleterious alleles and preserves the genetic buffering capacity at the level of developmental or regulatory pathways (*11, 12*).

More broadly, our findings support the view that developmental robustness is itself a biologically variable property that depends on genetic context. Canalization is understood as a buffering process that stabilizes phenotypes against perturbation (*21*), but our data show that this buffering can break down under extreme homozygosity. Importantly, this breakdown was evident across multiple levels of biological organization, including locomotor behavior, repeated behavioral consistency, bilateral wing symmetry, bristle patterning, eye morphology, and brain anatomy. The recurrent appearance of reduced repeatability and increased asymmetry across such diverse traits argues against a purely assay-specific explanation and instead suggests that homozygosity compromises general mechanisms of developmental stabilization. In this sense, the most consistent consequence of clonality in *D. mercatorum* was not simply a change in trait variation, but a loss of the robustness that normally constrains phenotypic divergence among genetically similar individuals.

These findings have direct implications for the design and interpretation of experiments that rely on genetically uniform animals. Inbred and isogenic strains are widely used because they are expected to reduce phenotypic variability and thereby improve reproducibility and statistical power (*4, 5, 7, 8*). Our results suggest this expectation has practically important biological limits. Beyond a threshold of homozygosity, developmental buffering may erode, producing phenotypic outcomes that are harder to predict and less reproducible across experiments, even within a genetically identical cohort. This may help explain why highly standardized animal models can still show substantial unexplained variability despite strong control of genotype and environment (*6, 51*). Controlled heterozygosity (*52*), rather than maximal genetic uniformity, may in some contexts provide a more robust experimental substrate by preserving developmental buffering while still limiting uncontrolled genetic variation.

Several limitations of our study should be considered. First, the specific route to clonality in *D. mercatorum*, pronuclear duplication following meiosis, is biologically unusual (*28, 53*), and the degree to which the same relationships between homozygosity, variability, and canalization hold in other inbred or clonal systems remains to be tested, despite strong supporting evidence in line with our study (*7, 15, 17, 19, 20, 54, 55*). Second, the effects we observed were not uniform across traits, sexes, or population backgrounds. Some traits showed increased variation, others showed no change, and a few even showed reduced variation; the magnitude of canalization effects also differed between strains. This heterogeneity is informative in itself, but it argues against overly simple generalizations. Our data do not imply that homozygosity universally increases phenotypic variability; rather, they show that extreme homozygosity reproducibly erodes developmental robustness, with consequences for trait means, variability structure, and repeatability that are context-dependent.

Taken together, our results show that genetic uniformity does not necessarily produce phenotypic uniformity. In *D. mercatorum*, extreme homozygosity broadly altered mean phenotypes, changed interindividual variability in trait-dependent ways, and most consistently reduced behavioral and developmental canalization. Outcrossing restored robustness and, in some cases, amplified it, identifying heterozygosity as an active stabilizing force in development and, by extension, in homeostasis. These findings challenge a foundational assumption of experimental biology, that more genetic uniformity means more phenotypic reproducibility, and suggest instead that developmental robustness depends on genetic context. Extreme homozygosity can itself become a source of instability, not just a means of reducing variability.

## Acknowledgements

The authors thank Alexis Sperling and David Glover for fly stocks and reagents. This work was supported by the Deutsche Forschungsgemeinschaft (DFG) through the DFG research unit 5289 RobustCircuit (G.A.L.), through grants LI 2640/1-1, FOR5289 LI 2640/2-1 (G.A.L.), and with support from the Fachbereich Biologie, Chemie & Pharmazie of the Freie Universität Berlin, as well as the Division of Neurobiology at Freie Universität Berlin. We thank members of the Linneweber lab, Mathias Wernet, Randolf Menzel, and Robin Hiesinger for helpful and intense discussions.

## Author Contributions

Conceptualization: G.A.L.; Methodology: M.W., A.K., H.H., M.R., M.A.H., G.D., T.M., E.R., J.B., M.v.K. and G.A.L.; Investigation: M.W., A.K., H.H., M.R., M.A.H., G.D., T.M., E.R., J.B, M.v.K. and G.A.L.; Resources: G.A.L.; Supervision: G.A.L.; Funding Acquisition: G.A.L.

---

**Supplementary Fig.S1:**
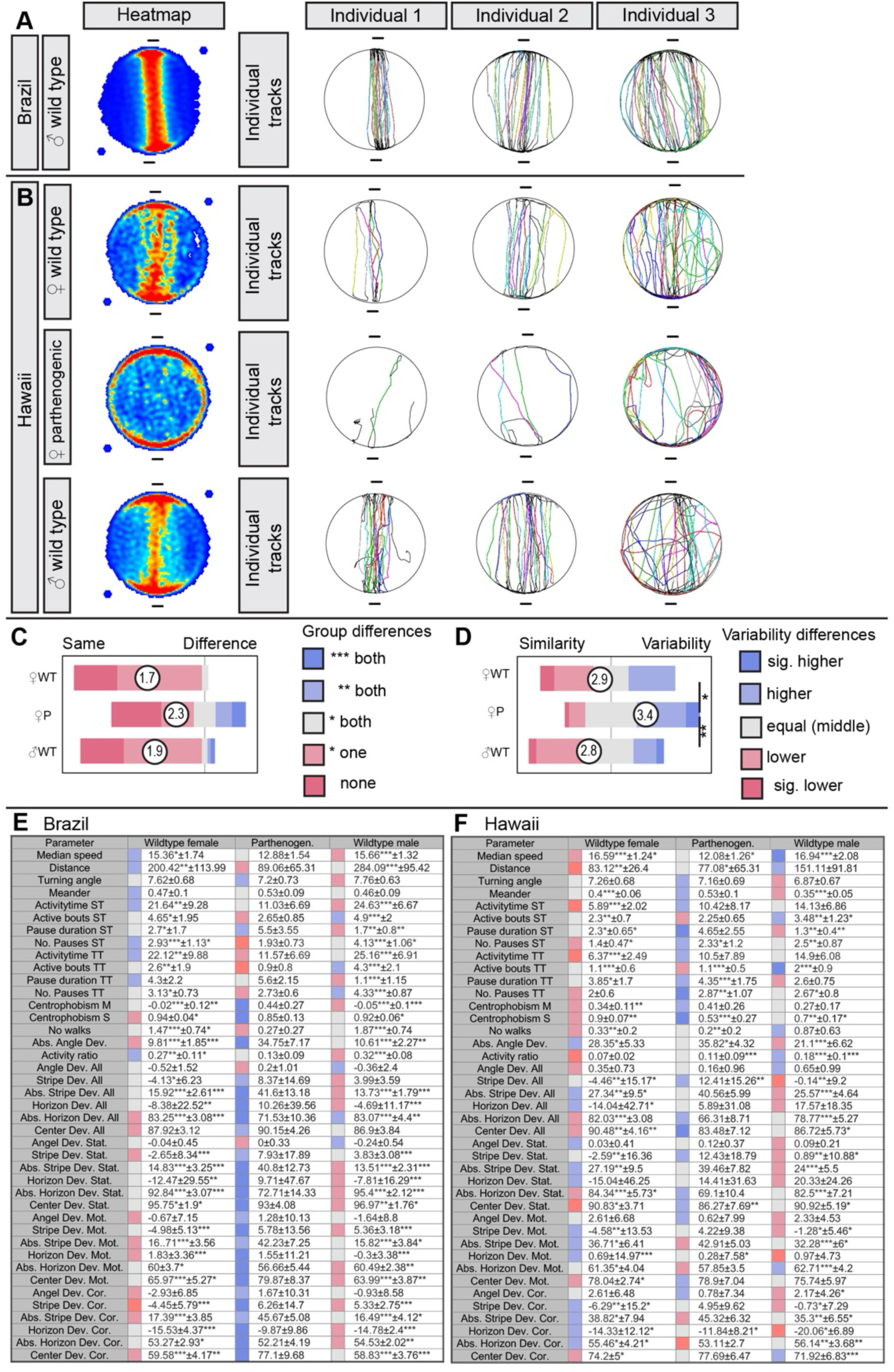
Buridan analysis of Hawaiian and Brazilian Drosophila mercatorum confirms reduced stripe fixation and increased interindividual behavioral variation in parthenogenic flies. **(A)** Mean occupancy heatmaps for Brazilian wild-type males show robust fixation of the visual stripes. Representative individual trajectories illustrate diverse patterns in Buridan’s paradigm **(B)** Mean occupancy heatmaps for Hawaiian wild-type and parthenogenic females reveal reduced stripe fixation in parthenogenic flies. Representative individual trajectories show greater phenotypic variation in parthenogenic females than in wild-type controls. **(C) Mean-difference index** across 41 behavioral parameters for the Hawaiian comparison indicates changes in behavioral means between parthenogenic females and control groups. Pairwise Wilcoxon tests with Benjamini-Hochberg correction: female WT versus parthenogenic, P = 0.1; female WT versus male WT, P = 0.8; parthenogenic versus male WT, P = 0.2. **(D) Variability index** based on the median absolute deviation (MAD) across the 41 behavioral parameters confirms increased interindividual behavioral variation in parthenogenic females. Pairwise Wilcoxon tests with Benjamini-Hochberg correction: female WT versus parthenogenic, P = 0.02; female WT versus male WT, P = 0.6; parthenogenic versus male WT, P = 0.003. **(E-F)** Full parameter-level data for the mean-difference and variation analyses for the Brazilian **(E)** and Hawaiian **(F)** wild-type and parthenogenic comparisons. Sample sizes: Brazil female WT, n = 33; parthenogenic, n = 31; male WT, n = 31; Hawaii female WT, n = 31; parthenogenic, n = 31; male WT, n = 30. Asterisks denote statistical significance: p < 0.05 (*), p < 0.01 (**), p < 0.001 (***).

**Supplementary Fig. S2:**
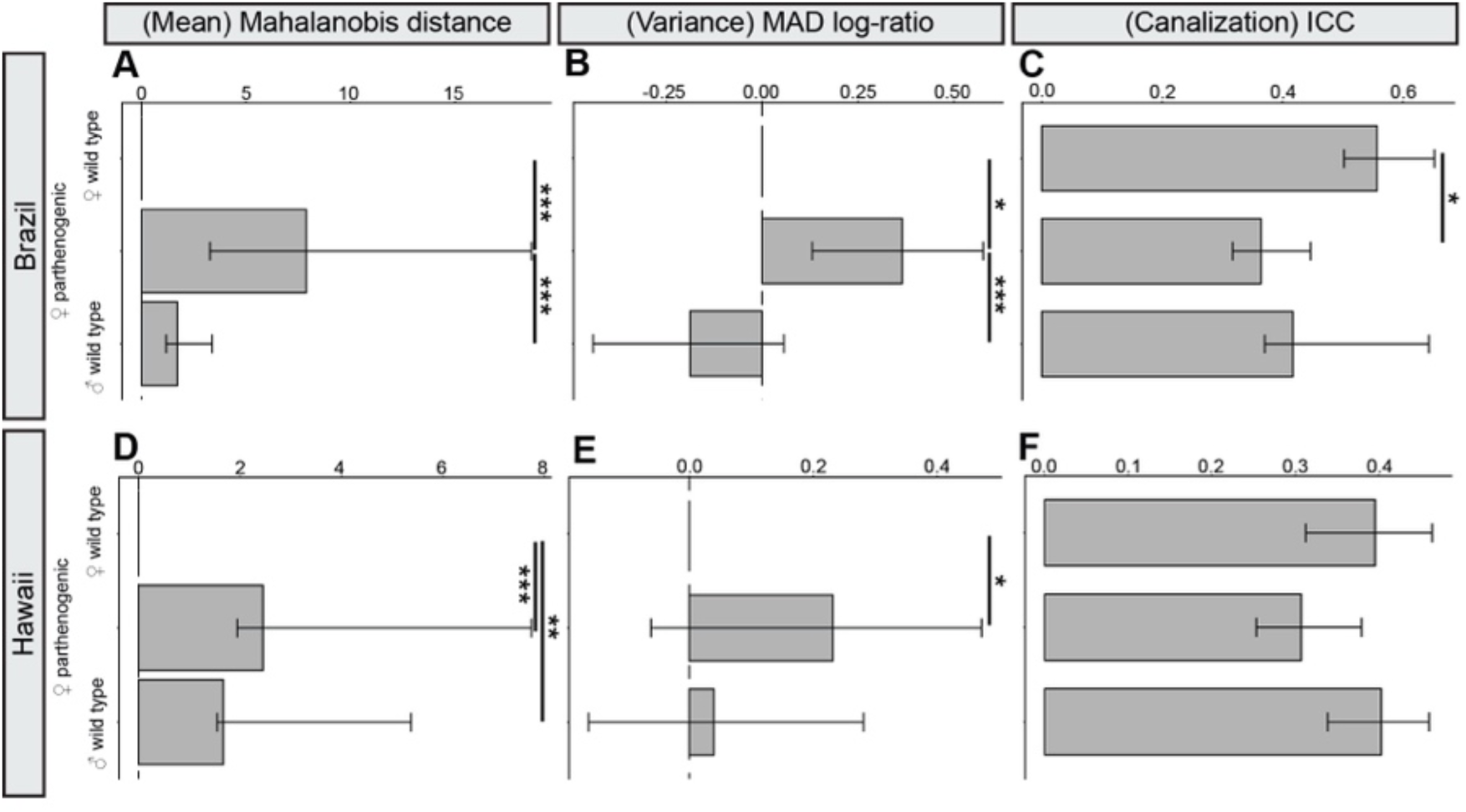
Reanalysis of Buridan data with multivariate and repeatability-based metrics confirms altered behavior, increased interindividual variation, and reduced behavioral consistency in parthenogenic flies. Buridan data from Fig. 1 and Supplementary Fig. S1 were reanalyzed using Mahalanobis distance, MAD log-ratio, and the intraclass correlation coefficient (ICC). **(A-C)** Brazil-derived flies. **(A)** Mahalanobis distance indicates a strong multivariate change in behavioral means in parthenogenic females relative to both control groups. Pairwise permutation tests with Benjamini-Hochberg correction: female WT versus parthenogenic, P < 0.001; female WT versus male WT, P = 0.7; parthenogenic versus male WT, P < 0.001. **(B)** MAD log-ratio confirms increased interindividual behavioral variation in parthenogenic females. Pairwise permutation tests with Benjamini-Hochberg correction: female WT versus parthenogenic, P = 0.011; female WT versus male WT, P = 0.14; parthenogenic versus male WT, P < 0.001. **(C)** ICC analysis indicates reduced behavioral consistency in parthenogenic females. Pairwise permutation tests with Benjamini-Hochberg correction: female WT versus parthenogenic, P = 0.012; female WT versus male WT, P = 0.062; parthenogenic versus male WT, P = 0.45. **(D-F)** Hawaii-derived flies. **(D)** Mahalanobis distance indicates a multivariate change in behavioral means between groups. Pairwise permutation tests with Benjamini-Hochberg correction: female WT versus parthenogenic, P < 0.001; female WT versus male WT, P = 0.001; parthenogenic versus male WT, P = 0.23. **(E)** MAD log-ratio indicates greater interindividual behavioral variation in parthenogenic females than in wild-type female controls. Pairwise permutation tests with Benjamini-Hochberg correction: female WT versus parthenogenic, P = 0.03; female WT versus male WT, P = 0.7; parthenogenic versus male WT, P = 0.077. **(F)** ICC analysis shows a trend towards reduced behavioral consistency in parthenogenic females, but no significant pairwise differences after correction. Pairwise permutation tests with Benjamini-Hochberg correction: female WT versus parthenogenic, P = 0.09; female WT versus male WT, P = 0.85; parthenogenic versus male WT, P = 0.09. Sample sizes: Brazil female WT, n = 33; parthenogenic, n = 31; male WT, n = 31; Hawaii female WT, n = 31; parthenogenic, n = 31; male WT, n = 30. Asterisks denote statistical significance: p < 0.05 (*), p < 0.01 (**), p < 0.001 (***).

**Supplementary Fig. S3:**
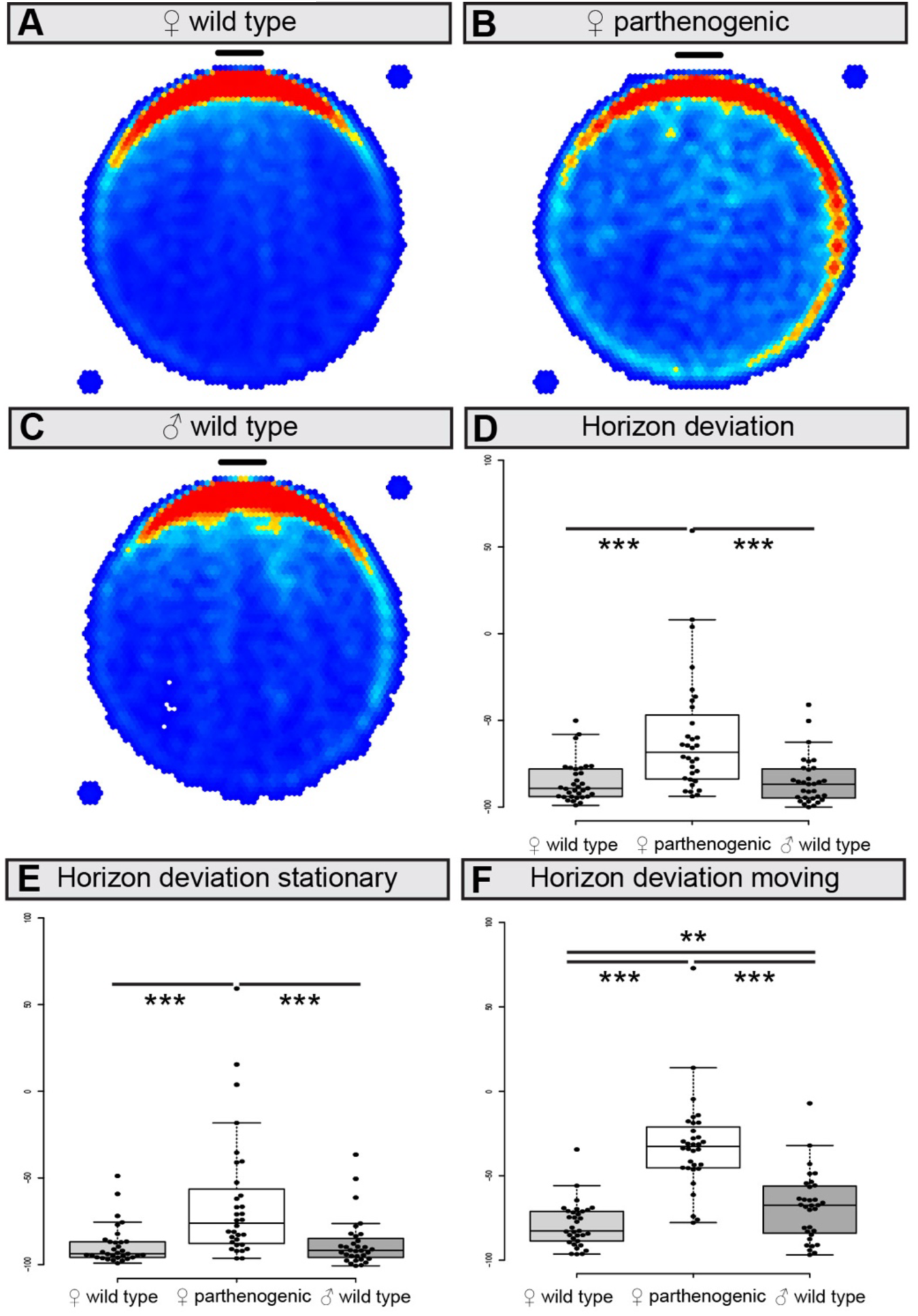
One-stripe Buridan assay reveals reduced, but preserved, visual stripe fixation in parthenogenic *Drosophila mercatorum*. **(A-C)** Representative occupancy heatmaps of female wild-type (WT) **(A)**, parthenogenic female **(B),** and male wild-type **(C)** flies in the one-stripe Buridan assay. All groups preferentially occupy the side containing the stripe, but parthenogenic flies show a broader spatial distribution and weaker stripe fixation than controls. **(D)** Quantification of stripe fixation using horizon deviation (−100% to +100%) shows a significant reduction of stripe fixation in parthenogenic flies, although all groups retain a clear preference for the stripe. Medians: −89.79% (female WT), −68.47% (parthenogenic), −86.79% (male WT). Pairwise Wilcoxon tests with Benjamini-Hochberg correction: female WT versus parthenogenic, P = 2.8 × 10⁻⁵; female WT versus male WT, P = 0.85; parthenogenic versus male WT, P = 2.8 × 10⁻⁵. **(E-F)** Similar effects are observed when horizon deviation is analyzed separately during stationary (e) and locomotor (f) periods. During stationary periods, medians were −93.79% (female WT), −76.10% (parthenogenic), and −91.80% (male WT), with pairwise Wilcoxon tests and Benjamini-Hochberg correction: P = 1.8 × 10⁻⁵, 0.86, and 4.0 × 10⁻⁵, respectively. During locomotion, medians were −82.65% (female WT), −32.59% (parthenogenic) and −67.65% (male WT), with corresponding P-values of 2.5 × 10⁻¹², 0.008 and 1.5 × 10⁻⁸.Sample sizes: female WT (N = 33), parthenogenic (N = 32), male WT (N = 32). Asterisks denote statistical significance: p < 0.05 (*), p < 0.01 (**), p < 0.001 (***).

**Supplementary Fig. S4:**
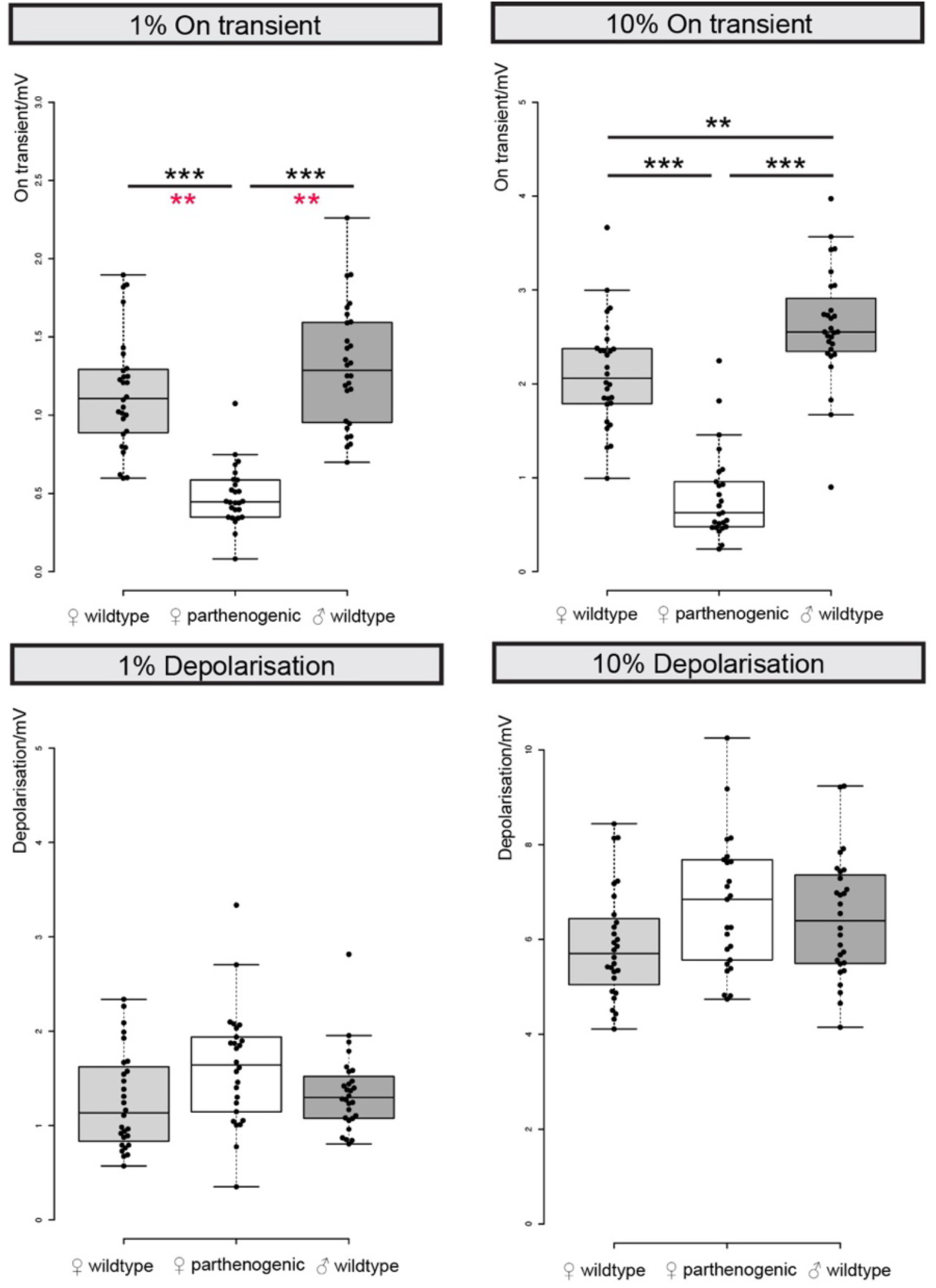
Electroretinogram recordings reveal reduced visual response amplitudes in parthenogenic *Drosophila mercatorum* without evidence of blindness. Boxplots compare electroretinogram (ERG) responses in female wild-type (WT), parthenogenic female, and male wild-type flies at 1% and 10% stimulus intensity. For each stimulus intensity, two response components were quantified: on-transient amplitude and sustained depolarization. **(A)** On-transient amplitude at 1% stimulus intensity is significantly reduced in parthenogenic flies relative to both wild-type control groups. Pairwise Wilcoxon tests with Benjamini-Hochberg correction: female WT versus parthenogenic, P = 6.9 × 10⁻⁹; female WT versus male WT, P = 0.33; parthenogenic versus male WT, P = 2.8 × 10⁻⁹. **(B)** On-transient amplitude at 10% stimulus intensity is also significantly reduced in parthenogenic flies. Pairwise Wilcoxon tests with Benjamini-Hochberg correction: female WT versus parthenogenic, P = 1.9 × 10⁻⁸; female WT versus male WT, P = 0.001; parthenogenic versus male WT, P = 6.1 × 10⁻⁹. **(C)** Sustained depolarization at 1% stimulus intensity does not differ significantly between groups. Pairwise Wilcoxon tests with Benjamini-Hochberg correction: female WT versus parthenogenic, P = 0.07; female WT versus male WT, P = 0.32; parthenogenic versus male WT, P = 0.16. **(D)** Sustained depolarization at 10% stimulus intensity likewise shows no significant group differences. Pairwise Wilcoxon tests with Benjamini-Hochberg correction: female WT versus parthenogenic, P = 0.09; female WT versus male WT, P = 0.13; parthenogenic versus male WT, P = 0.41. Sample sizes: female WT, n = 28; parthenogenic, n = 26; male WT, n = 28. Asterisks denote statistical significance: p < 0.05 (*), p < 0.01 (**), p < 0.001 (***).

**Supplementary Fig. S5:**
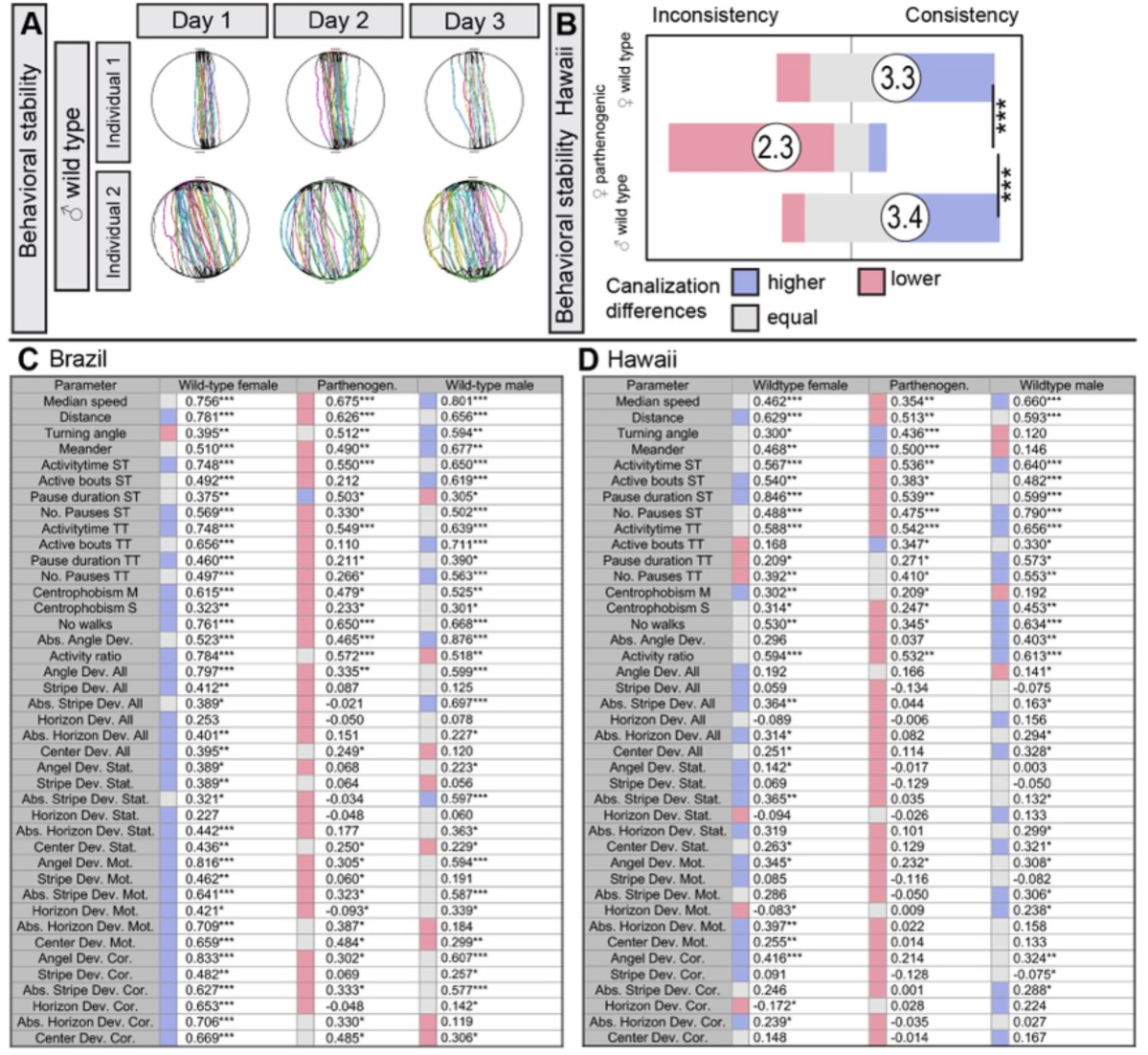
Repeated Buridan testing confirms reduced behavioral consistency in parthenogenic *Drosophila mercatorum* from both Brazilian and Hawaiian strains. **(A)** Representative trajectories from 3 consecutive days of testing show high day-to-day behavioral consistency in Brazilian wild-type (WT) males. **(B)** Behavioral consistency (canalization) index across 41 Buridan parameters for Hawaiian flies confirms reduced day-to-day behavioral stability in parthenogenic females relative to both wild-type control groups. Pairwise Wilcoxon tests with Benjamini-Hochberg correction: female WT versus parthenogenic, P = 3.0 × 10⁻⁸; female WT versus male WT, P = 0.8; parthenogenic versus male WT, P = 4.2 × 10⁻⁹. **(C-D)** Full parameter-level behavioral canalization analysis based on Pearson correlation coefficients for the Brazilian (C) and Hawaiian (C) comparisons. Sample sizes: Brazil: female WT (N = 33), parthenogenic (N = 31), male WT (N = 31), Hawaii: female WT (N = 31), parthenogenic (N = 31), male WT (N = 30). Asterisks denote significant correlations across one-day (*), two-day (**), and three-day (***) interval comparisons.

**Supplementary Fig. S6:**
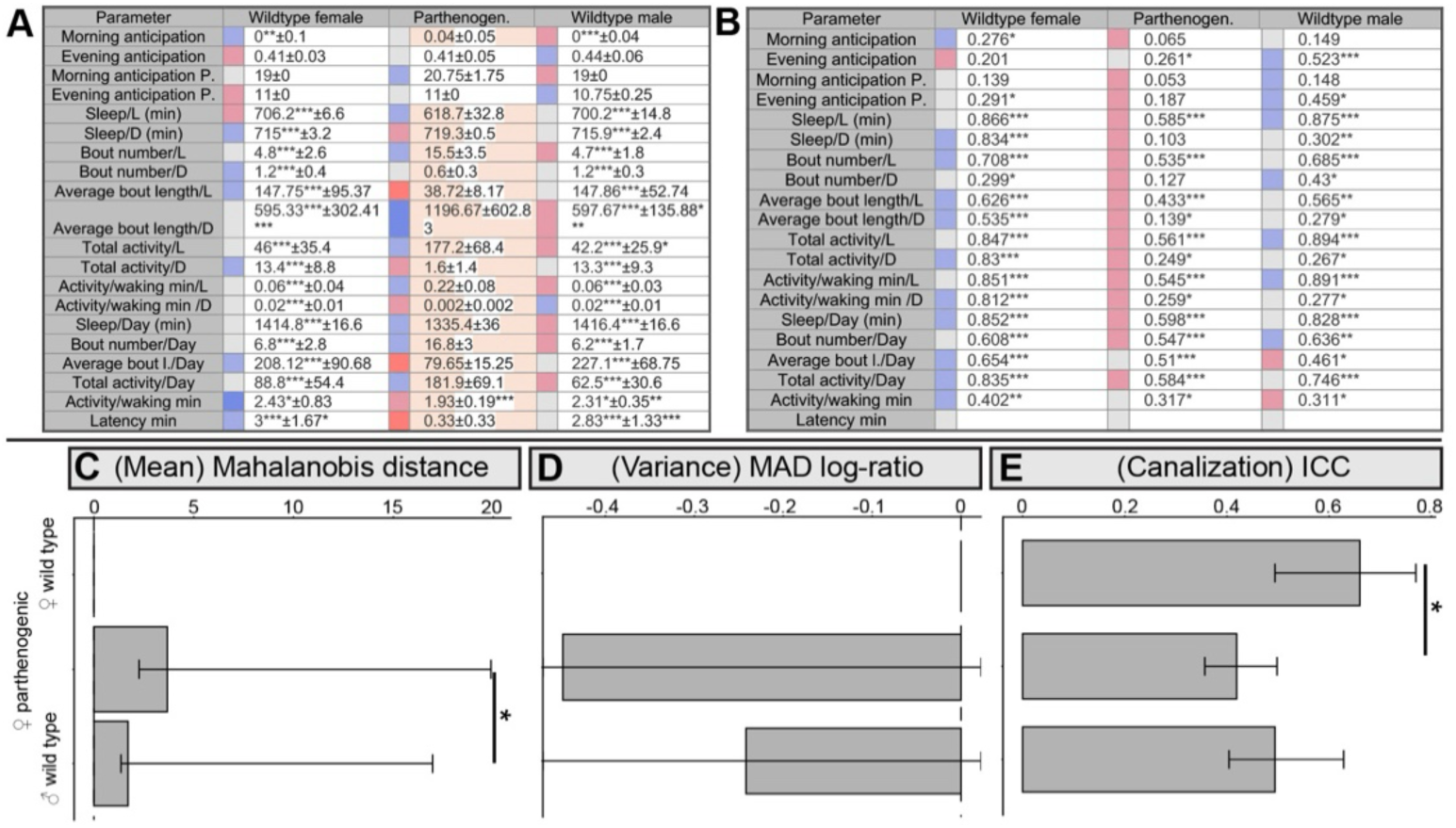
Full statistical analysis and alternative reanalysis of locomotor activity and sleep in Brazilian parthenogenic *Drosophila mercatorum*. **(A)** Parameter-level analysis of mean and dispersion across 21 locomotor activity and sleep traits for female wild-type, parthenogenic female, and male wild-type flies. The table shows group medians, with significance assessed by pairwise Wilcoxon tests with Benjamini-Hochberg correction, and median absolute deviation (MAD), with significance assessed by Levene’s test. Color scale denotes relative MAD values, from low (blue) to high (red), and darker colors indicate parameters that differ significantly from both comparison groups. **(B)** Parameter-level analysis of day-to-day behavioral consistency across 3 recording days for 20 of the 21 activity and sleep parameters. The table shows Pearson correlation coefficients for all pairwise day comparisons after Fisher’s z-transformation. Asterisks indicate parameters with significant correlations across the three-day pair comparisons. Color scale denotes relative correlation strength, from low (red) to high (blue), with intermediate values in grey. **(C-E)** Alternative reanalysis of the same locomotor dataset shown in (A-B). (C) Mahalanobis distance indicates no significant difference between female wild-type and parthenogenic flies or between the two wild-type groups, but a significant multivariate change between parthenogenic females and wild-type males. Pairwise permutation tests with Benjamini-Hochberg correction: female WT versus parthenogenic, P = 0.28; female WT versus male WT, P = 0.31; parthenogenic versus male WT, P = 0.01. (D) MAD log-ratio indicates no significant difference in overall interindividual variation between groups. Pairwise permutation tests with Benjamini-Hochberg correction: female WT versus parthenogenic, P = 0.46; female WT versus male WT, P = 0.46; parthenogenic versus male WT, P = 0.46. (E) Intraclass correlation coefficient (ICC) analysis confirms reduced temporal behavioral consistency in parthenogenic females compared with wild-type female controls. Pairwise permutation tests with Benjamini-Hochberg correction: female WT versus parthenogenic, P = 0.012; female WT versus male WT, P = 0.063; parthenogenic versus male WT, P = 0.39. Sample sizes: female WT, n = 33; parthenogenic, n = 32; male WT, n = 32. Asterisks denote statistical significance: p < 0.05 (*), p < 0.01 (**), p < 0.001 (***). In (B), asterisks denote significant correlations across one-day (*), two-day (**), and three-day (***) interval comparisons.

**Supplementary Fig. S7:**
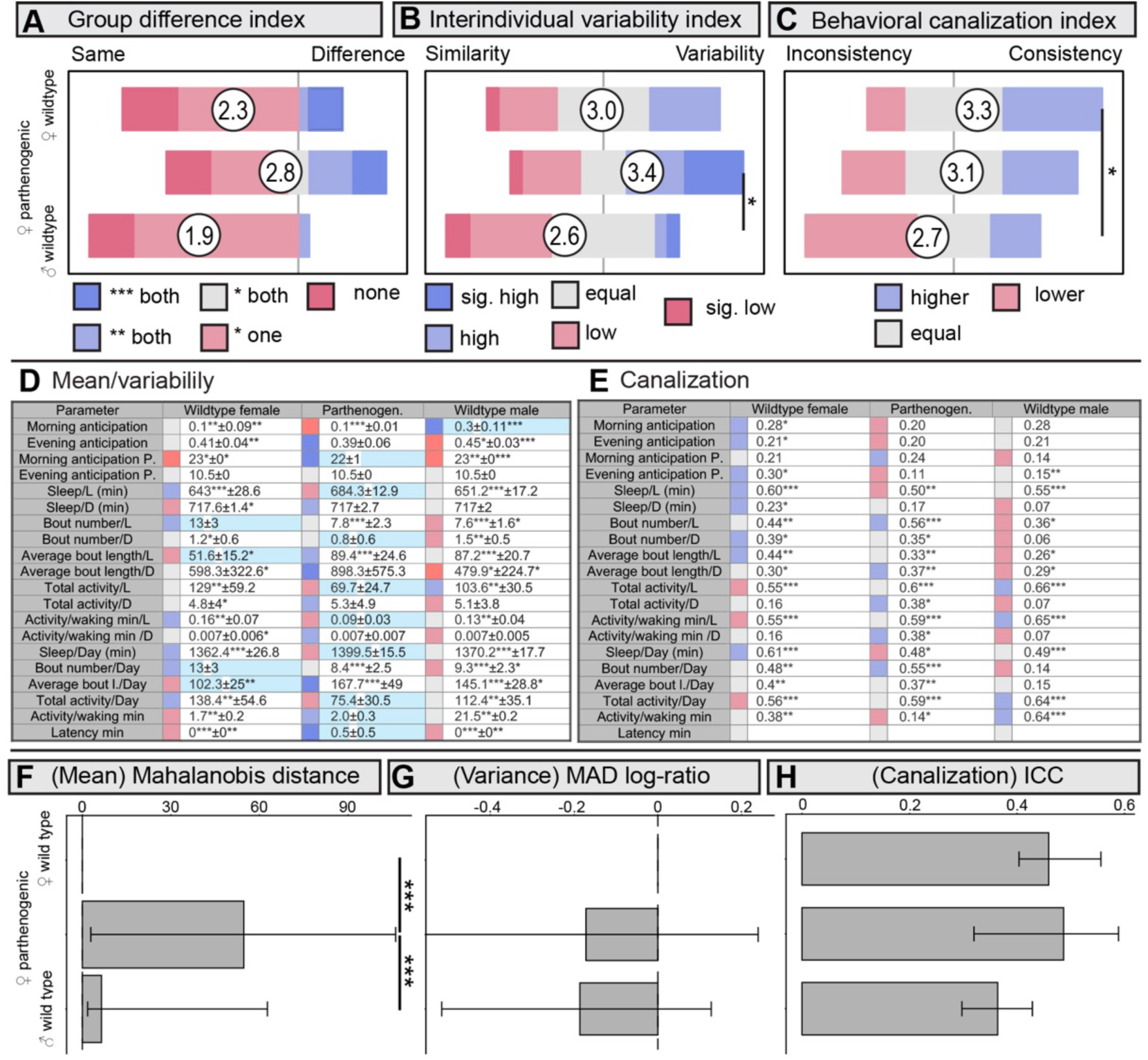
Locomotor activity and sleep analysis in Hawaiian *Drosophila mercatorum* reveal modest genotype-dependent differences without a strong reduction in temporal behavioral consistency in parthenogenic flies. Activity and sleep measurements in Hawaiian wild-type and parthenogenic *D. mercatorum* show genotype-dependent differences in behavioral means and dispersion that are weaker than those observed in the Brazilian comparison. **(A) Mean-difference index** across 21 locomotor activity and sleep parameters indicates modest changes in behavioral means in parthenogenic females relative to the two wild-type (WT) control groups. Full parameter-level data are shown in (D). Pairwise Wilcoxon tests with Benjamini-Hochberg correction: female WT versus parthenogenic, P = 0.29; female WT versus male WT, P = 0.62; parthenogenic versus male WT, P = 0.06. **(B) Variability index** based on the median absolute deviation (MAD) across the 21 parameters shows a trend towards increased interindividual variation in parthenogenic females. Pairwise Wilcoxon tests with Benjamini-Hochberg correction: female WT versus parthenogenic, P = 0.23; female WT versus male WT, P = 0.18; parthenogenic versus male WT, P = 0.04. **(C) Behavioral canalization index** across 20 repeatedly measured activity and sleep parameters shows sexual dimorphism in temporal behavioral consistency, but no significant reduction in canalization in parthenogenic females relative to female wild-type controls. Pairwise Wilcoxon tests with Benjamini-Hochberg correction: female WT versus parthenogenic, P = 0.41; female WT versus male WT, P = 0.04; parthenogenic versus male WT, P = 0.22. **(D)** Parameter-level analysis of mean and dispersion across all 21 locomotor activity and sleep traits. The table shows group medians, with significance assessed by pairwise Wilcoxon tests with Benjamini-Hochberg correction, and MAD values, with significance assessed by Levene’s test. Colors denote relative MAD values, from low (blue) to high (red), and darker colors indicate parameters that differ significantly between the two comparison groups. **(E)** Parameter-level analysis of day-to-day behavioral consistency across 3 recording days for 20 of the 21 parameters. The table shows Pearson correlation coefficients for all pairwise day comparisons after Fisher’s z-transformation. Colors denote relative correlation strength, from low (red) to high (blue), with intermediate values in grey. **(F-H)** Reanalysis of the same behavioral data shown in (D-E). (**F)** Mahalanobis distance indicates a significant multivariate change in behavioral means between parthenogenic females and both wild-type control groups. Pairwise permutation tests with Benjamini-Hochberg correction: female WT versus parthenogenic, P < 0.001; female WT versus male WT, P = 0.31; parthenogenic versus male WT, P < 0.001. **(G)** MAD log-ratio indicates no significant difference in overall interindividual variation between groups. Pairwise permutation tests with Benjamini-Hochberg correction: female WT versus parthenogenic, P = 0.38; female WT versus male WT, P = 0.94; parthenogenic versus male WT, P = 0.38. **(H)** Intraclass correlation coefficient (ICC) analysis shows no significant difference in temporal behavioral consistency between groups. Pairwise permutation tests with Benjamini-Hochberg correction: female WT versus parthenogenic, P = 0.64; female WT versus male WT, P = 0.23; parthenogenic versus male WT, P = 0.12. Sample sizes: Hawaii female WT, n = 31; parthenogenic, n = 31; male WT, n = 30. Asterisks denote statistical significance: p < 0.05 (*), p < 0.01 (**), p < 0.001 (***). Only in (E), asterisks indicate the number of significant comparisons across days. Sample sizes: female WT (N = 27), parthenogenic (N = 27), male WT (N = 29).

**Supplementary Fig. S8:**
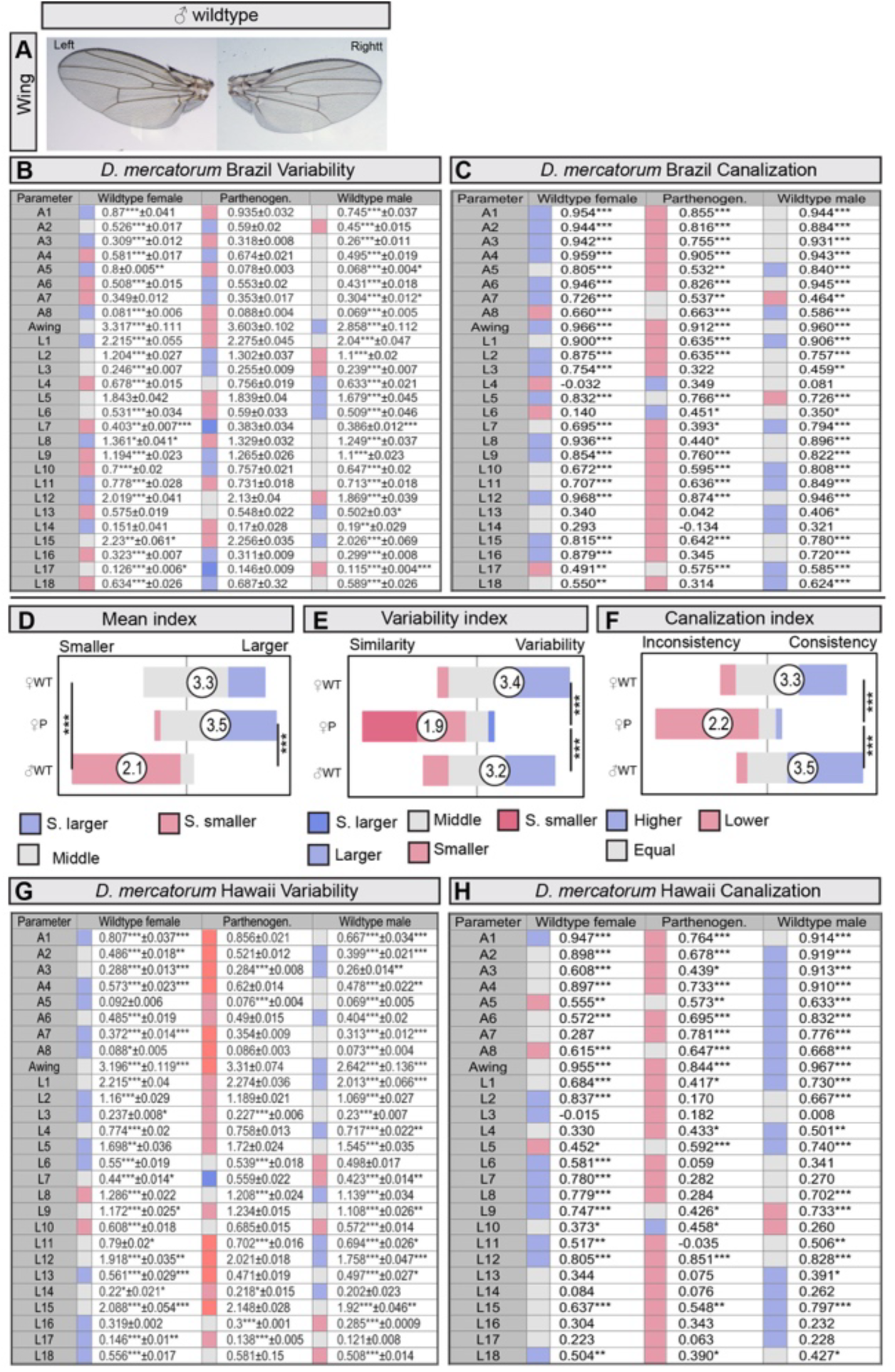
Wing morphometry analysis in wild-type and parthenogenic *Drosophila mercatorum* reveals altered morphology, reduced interindividual variation, and reduced developmental canalization in parthenogenic flies. **(A)** Representative wing image from a Brazilian wild-type (WT) male *D. mercatorum*. **(B)** Parameter-level analysis of mean and dispersion across all 27 wing traits in Brazilian flies. The table shows group medians, with significance assessed by pairwise Wilcoxon tests with Benjamini-Hochberg correction, and median absolute deviation (MAD), with significance assessed by Levene’s test. Colors denote relative MAD values, from low (blue) to high (red), and darker colors indicate parameters that differ significantly between the two comparison groups. **C)** Parameter-level analysis of left-right correspondence across the 27 wing traits in Brazilian flies. The table shows Pearson correlation coefficients between right and left wings, with asterisks denoting statistical significance. Colors denote correlation strength, from low (red) to high (blue), with intermediate values in grey. **(D) Mean-difference index** across 27 wing parameters for Hawaiian flies indicates broad changes in wing morphology in wild-type males relative to both female groups. In contrast, female wild-type and parthenogenic flies do not differ significantly in overall mean trait values. Full parameter-level data are shown in (G). Pairwise Wilcoxon tests with Benjamini-Hochberg correction: female WT versus parthenogenic, P = 0.2; female WT versus male WT, P = 2.5 × 10⁻¹⁰; parthenogenic versus male WT, P = 7.0 × 10⁻¹⁰. **(E) Variability index** based on the median absolute deviation (MAD) across the 27 wing parameters indicates reduced interindividual variation in Hawaiian parthenogenic females. Full parameter-level data are shown in (G). Pairwise Wilcoxon tests with Benjamini-Hochberg correction: female WT versus parthenogenic, P = 2.9 × 10⁻⁷; female WT versus male WT, P = 0.3; parthenogenic versus male WT, P = 4.2 × 10⁻⁶. **(F) Canalization index** based on left-right correspondence across the 27 wing parameters confirms reduced developmental stability in Hawaiian parthenogenic flies. Full parameter-level data are shown in (H). Pairwise Wilcoxon tests with Benjamini-Hochberg correction: female WT versus parthenogenic, P = 4.9 × 10⁻⁷; female WT versus male WT, P = 0.1; parthenogenic versus male WT, P = 3.4 × 10⁻⁸. **(G)** Parameter-level analysis of mean and dispersion across all 27 wing traits in Hawaiian flies. The table shows group medians, with significance assessed by pairwise Wilcoxon tests with Benjamini-Hochberg correction, and MAD values, with significance assessed by Levene’s test. Colors denote relative MAD values, from low (blue) to high (red), and darker colors indicate parameters that differ significantly between the two comparison groups. **(H)** Parameter-level analysis of left-right correspondence across the 27 wing traits in Hawaiian flies. The table shows Pearson correlation coefficients between right and left wings, with asterisks denoting statistical significance. Sample sizes: Brazil: female WT (N = 29), parthenogenic (N = 31), male WT (N = 33). Hawaii: female WT (N = 30), parthenogenic (N = 30), male WT (N = 30). Asterisks denote statistical significance: p < 0.05 (*), p < 0.01 (**), p < 0.001 (***). Only in (C) and (H) do asterisks refer to the number of significant comparisons between different days.

**Supplementary Fig. S9:**
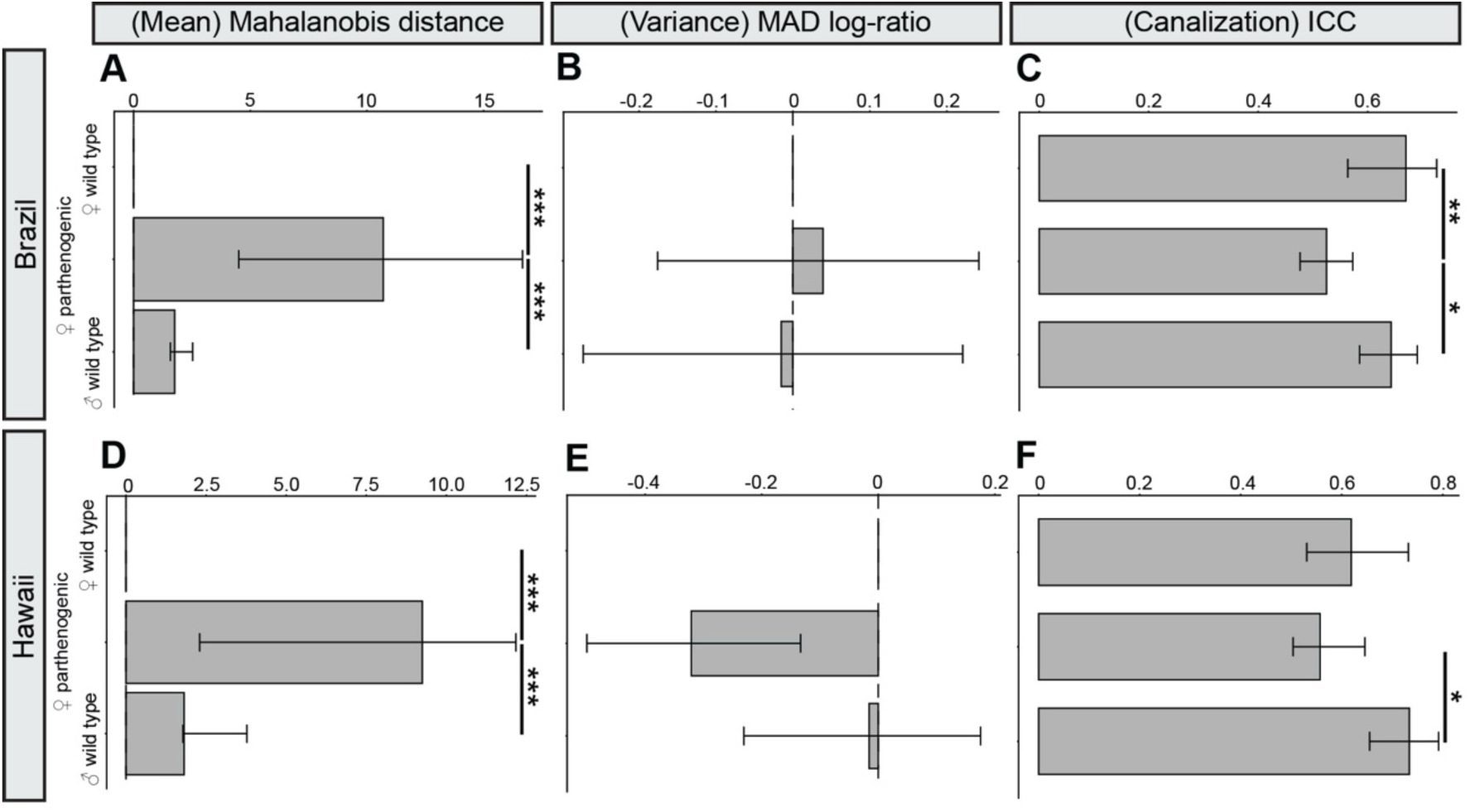
Reanalysis of wing morphometry confirms altered morphology and reduced developmental canalization in parthenogenic *Drosophila mercatorum*. Wing data from Fig. 3 and Supplementary Fig. S8 were reanalyzed using Mahalanobis distance, MAD log-ratio, and the intraclass correlation coefficient (ICC). **(A-C)** Brazil-derived flies. (A) Mahalanobis distance indicates a significant multivariate change in wing morphology in parthenogenic females relative to both wild-type control groups. Pairwise permutation tests with Benjamini-Hochberg correction: female WT versus parthenogenic, P < 0.001; female WT versus male WT, P = 0.37; parthenogenic versus male WT, P < 0.001. (B) MAD log-ratio indicates no significant difference in overall interindividual wing variation between groups. Pairwise permutation tests with Benjamini-Hochberg correction: female WT versus parthenogenic, P = 0.88; female WT versus male WT, P = 0.88; parthenogenic versus male WT, P = 0.88. (C) ICC analysis indicates reduced left-right correspondence in parthenogenic females, consistent with reduced developmental canalization. Pairwise permutation tests with Benjamini-Hochberg correction: female WT versus parthenogenic, P = 0.003; female WT versus male WT, P = 0.43; parthenogenic versus male WT, P = 0.02. **(D-F)** Hawaii-derived flies. (D) Mahalanobis distance likewise indicates a significant multivariate change in wing morphology in parthenogenic females relative to both wild-type control groups. Pairwise permutation tests with Benjamini-Hochberg correction: female WT versus parthenogenic, P < 0.001; female WT versus male WT, P = 0.16; parthenogenic versus male WT, P < 0.001. (E) MAD log-ratio indicates reduced interindividual wing variation in Hawaiian parthenogenic females. Pairwise permutation tests with Benjamini-Hochberg correction: female WT versus parthenogenic, P = 0.02; female WT versus male WT, P = 0.9; parthenogenic versus male WT, P = 0.02. (F) ICC analysis shows no significant difference between Hawaiian parthenogenic and female wild-type flies, although parthenogenic females differ from wild-type males. Pairwise permutation tests with Benjamini-Hochberg correction: female WT versus parthenogenic, P = 0.42; female WT versus male WT, P = 0.15; parthenogenic versus male WT, P = 0.024. Sample sizes: Brazil: female WT (N = 29), parthenogenic (N = 31), male WT (N = 33). Hawaii: female WT (N = 30), parthenogenic (N = 30), male WT (N = 30). Asterisks denote statistical significance: p < 0.05 (*), p < 0.01 (**), p < 0.001 (***).

**Supplementary Fig. S10:**
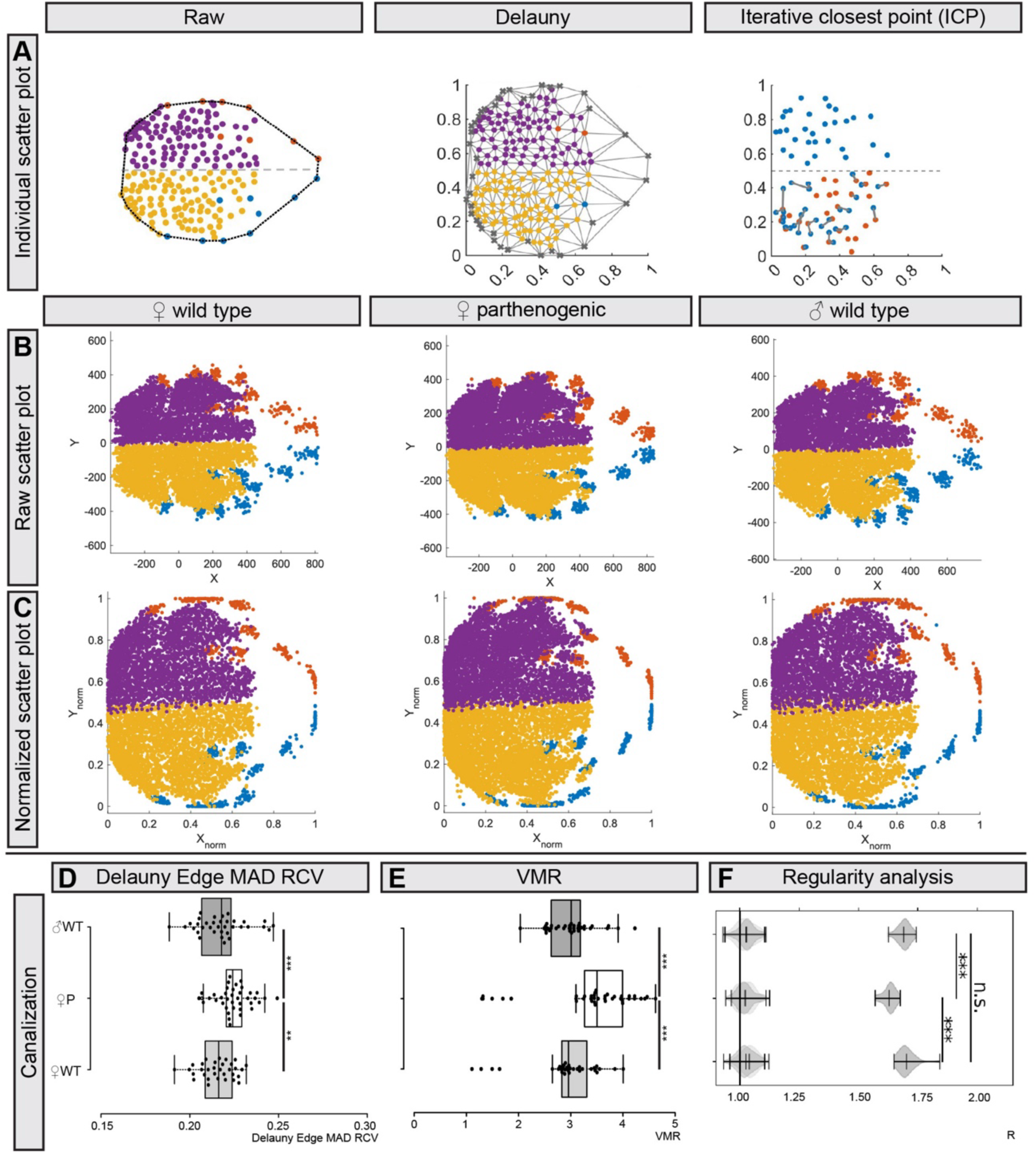
Spatial analysis of thoracic bristle patterns reveals reduced regularity in parthenogenic Drosophila mercatorum. **(A)** Overview of the thorax-bristle analysis pipeline. Left, raw manually annotated bristle coordinates from an individual fly (yellow, small left bristles; magenta, small right bristles; blue, large left bristles; red, large right bristles). Middle, bristle coordinates normalized to unit scale (0-1) to enable size-independent comparison between individuals; grey Delaunay edges represent the shortest connections between neighboring bristles. Right, point-to-point mismatches (grey lines) after mirroring the right-side bristles, randomly sampling subsets from each side, aligning them by iterative closest point (ICP), and computing distances to the nearest bristles on the left side. For each individual, the mismatch score was averaged over 1,000 bootstrap iterations. **(B)** Centroid-aligned raw bristle coordinates for all individuals in the tested population: wild-type females (left), parthenogenic females (middle), and wild-type males (right). **(C)** Same as (B), but with bristle coordinates normalized to unit scale (0-1) to remove overall size differences and permit unbiased comparison of spatial patterning between genotypes. **(D-F)** Additional group-level metrics quantifying canalization and spatial regularity of thoracic bristle patterns. **(D)** Robust coefficient of intra-individual variation (RCV), calculated as the median absolute deviation (MAD) of Delaunay edge lengths divided by the individual median edge length. Pairwise Wilcoxon tests with Benjamini-Hochberg correction: female WT versus parthenogenic, P = 0.0006; female WT versus male WT, P = 0.75; parthenogenic versus male WT, P = 0.0042. **(E)** Variance-to-mean ratio (VMR) of bristle density across 5 × 5 quadrats, quantifying spatial dispersion. Pairwise Wilcoxon tests with Benjamini-Hochberg correction: female WT versus parthenogenic, P = 7.1 × 10⁻^5^; female WT versus male WT, P = 0.93; parthenogenic versus male WT, P = 7.1 × 10⁻^5^. **(F)** Regularity analysis indicates reduced spatial regularity in parthenogenic flies. Pairwise Wilcoxon tests with Benjamini-Hochberg correction: female WT versus parthenogenic, P = 2.0 × 10⁻^10^; female WT versus male WT, P = 0.52; parthenogenic versus male WT, P = 9.0 × 10⁻^13^ Sample sizes: female WT (N = 29), parthenogenic (N = 31), male WT (N = 33). Asterisks denote statistical significance: p < 0.05 (*), p < 0.01 (**), p < 0.001 (***).

**Supplementary Fig. S11:**
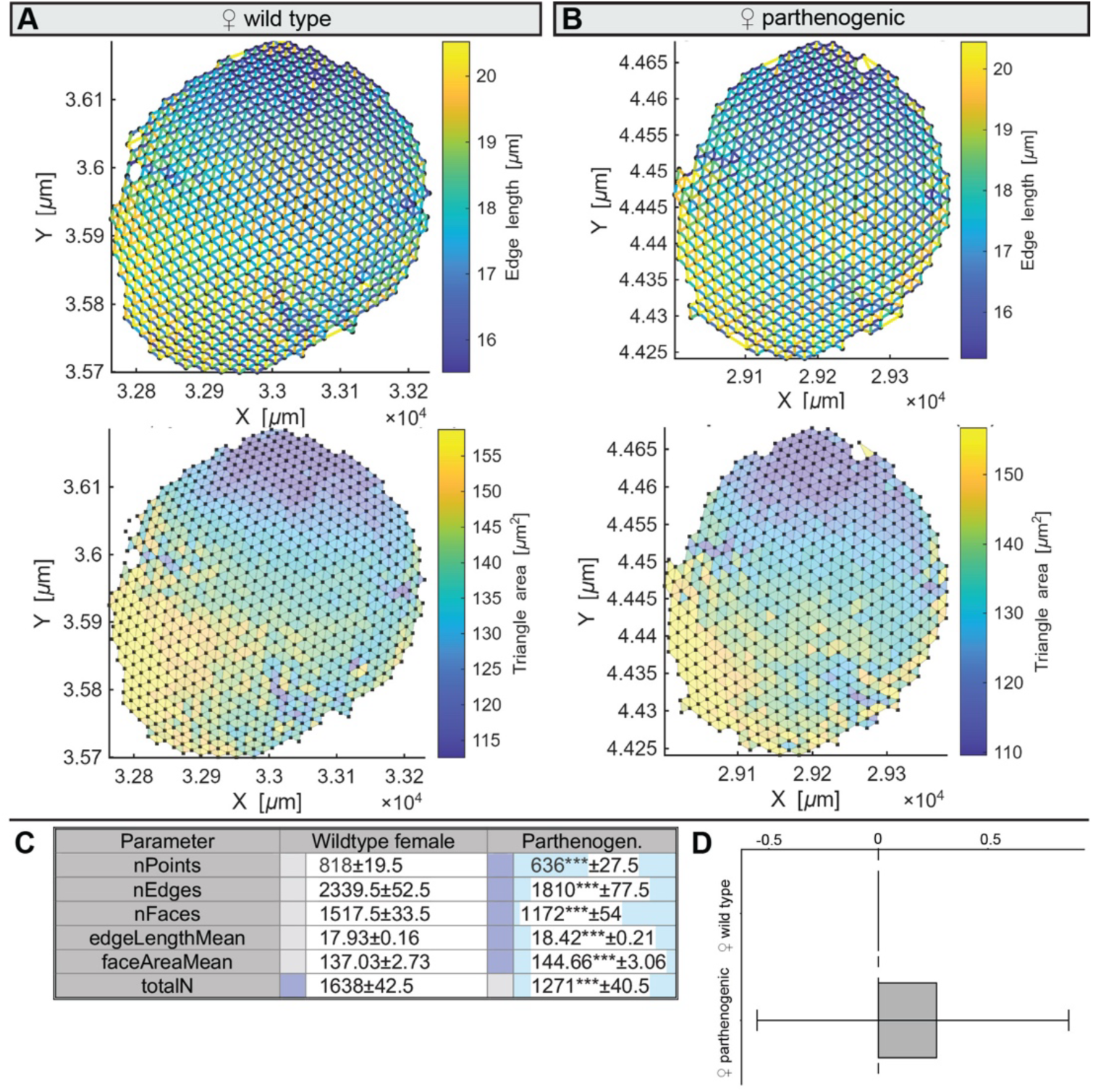
Eye lattice analysis reveals altered ommatidial geometry and a trend towards increased interindividual variation in parthenogenic *Drosophila mercatorum*. **(A-B)** Representative local lattice geometry from one eye of a wild-type (WT) **(A)** and a parthenogenic **(B)** Brazilian female, shown as edge lengths and triangle areas derived from neighboring ommatidia. **(C)** Parameter-level analysis of mean and dispersion across all six eye parameters in Brazilian wild-type and parthenogenic flies. The table shows group medians, with significance assessed by pairwise Wilcoxon tests with Benjamini-Hochberg correction, and median absolute deviation (MAD), with significance assessed by Levene’s test. Colors denote relative MAD values, with blue indicating the highest MAD values and darker colors indicating significant differences. **(D)** Reanalysis of the same dataset using MAD log-ratio shows no significant difference in overall interindividual variation between wild-type and parthenogenic flies. Pairwise permutation test with Benjamini-Hochberg correction: female WT versus parthenogenic, P = 0.08. Sample sizes: female WT (N = 10), parthenogenic (N = 10). Asterisks denote statistical significance: p < 0.05 (*), p < 0.01 (**), p < 0.001 (***).

**Supplementary Fig. S12:**
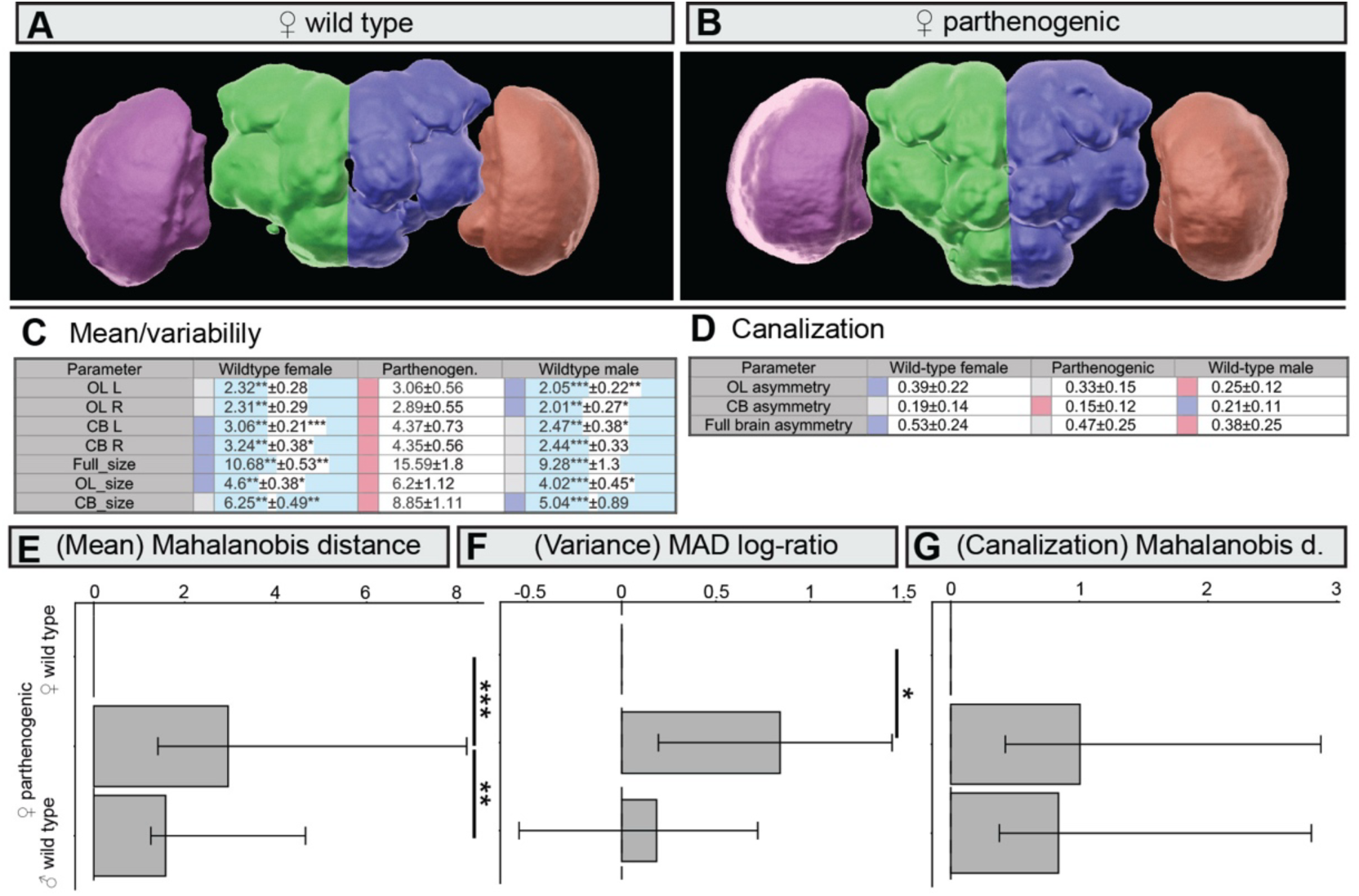
Nc82-based analysis of brain neuropiles reveals enlarged and more variable brains in parthenogenic *Drosophila mercatorum*. **(A-B)** Representative three-dimensional surface reconstructions of brains from Brazilian wild-type (WT) **(A)** and parthenogenic **(B)** females. Four neuropile regions are shown: left optic lobe (OL_L), left central brain (CB_L), right central brain (CB_R), and right optic lobe (OL_R). **(C)** Parameter-level analysis of mean and dispersion across all seven brain-neuropile parameters in Brazil-derived flies. The table shows group medians, with significance assessed by pairwise Wilcoxon tests with Benjamini-Hochberg correction, and median absolute deviation (MAD), with significance assessed by Levene’s test. colors denote relative MAD values, with red indicating the highest MAD values and blue the lowest; darker colors denote statistically significant differences. **(D)** Parameter-level analysis of left-right asymmetry across the three bilateral brain-neuropile parameters. Relative asymmetry is color-coded, with blue indicating the highest asymmetry and red the lowest. **(E-G)** Reanalysis of the same data shown in (C-D). **(E)** Mahalanobis distance indicates a significant multivariate change in brain neuropile anatomy in parthenogenic females relative to both wild-type control groups. Pairwise permutation tests with Benjamini-Hochberg correction: female WT versus parthenogenic, P < 0.001; female WT versus male WT, P = 0.06; parthenogenic versus male WT, P = 0.007. **(F)** MAD log-ratio indicates increased interindividual variation in parthenogenic females relative to female wild-type controls. Pairwise permutation tests with Benjamini-Hochberg correction: female WT versus parthenogenic, P = 0.045; female WT versus male WT, P = 0.68; parthenogenic versus male WT, P = 0.11. **(G)** Mahalanobis distance of symmetry values indicates no significant difference in overall left-right asymmetry between groups after correction. Pairwise permutation tests with Benjamini-Hochberg correction: female WT versus parthenogenic, P = 0.089; female WT versus male WT, P = 0.33; parthenogenic versus male WT, P = 0.67. Sample sizes: female WT (N = 22), parthenogenic (N = 23), male WT (N = 16). Asterisks denote statistical significance: p < 0.05 (*), p < 0.01 (**), p < 0.001 (***).

**Supplementary Fig. S13:**
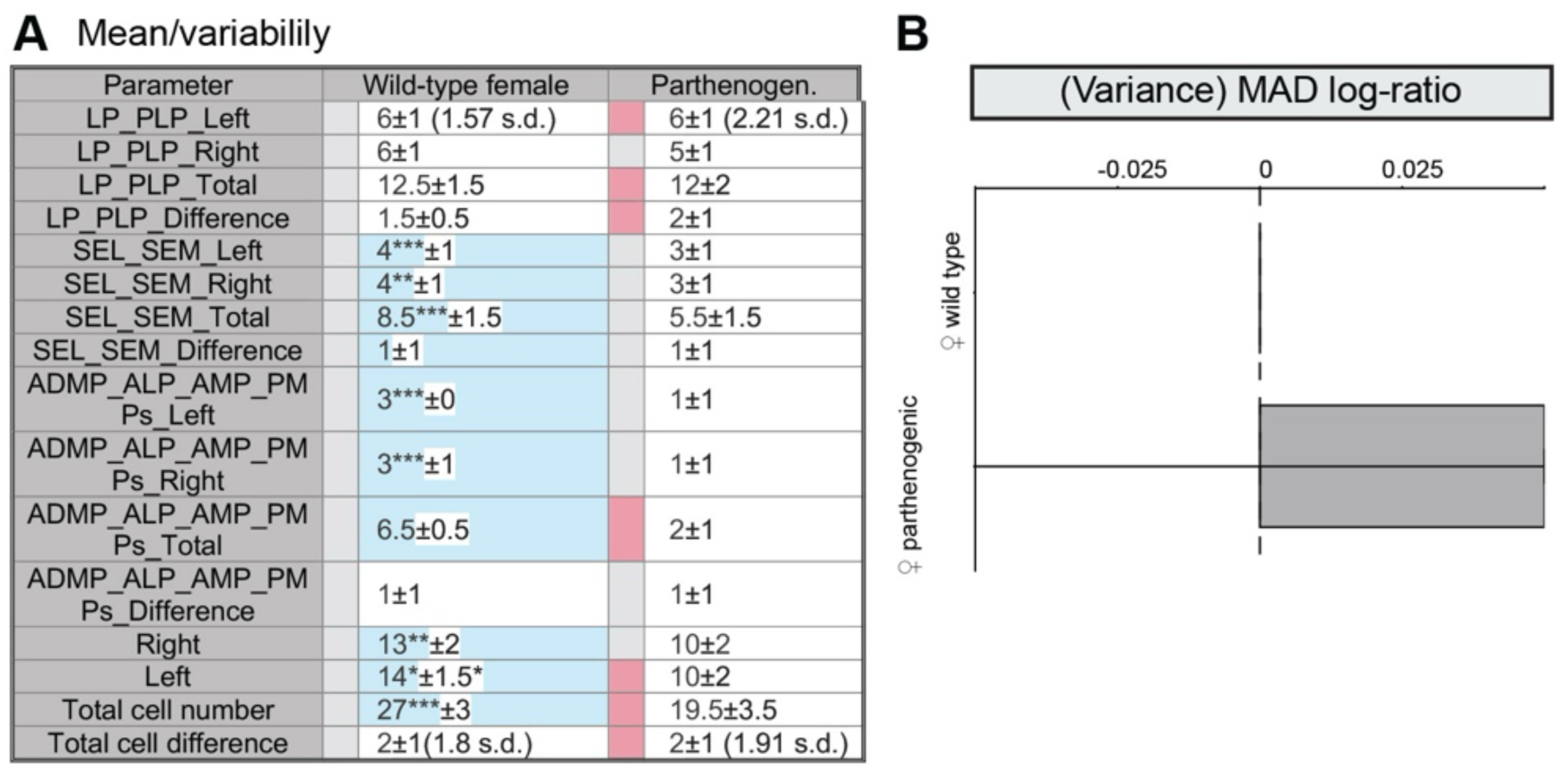
Quantitative analysis of serotonergic neurons reveals reduced neuron number, and MAD log-ratio analysis reveals a trend towards altered variation in parthenogenic *Drosophila mercatorum*. **(A)** Parameter-level analysis of mean and dispersion across all serotonergic-neuron parameters in Brazilian *D. mercatorum* flies. The table shows group medians, with significance assessed by pairwise Wilcoxon tests with Benjamini-Hochberg correction, and median absolute deviation (MAD), with significance assessed by Levene’s test. Colors denote relative MAD values, with red indicating the highest MAD values and blue the lowest; darker colors denote statistically significant differences. **(B)** Reanalysis of the same dataset using MAD log-ratio shows no significant difference in overall interindividual variation between groups after correction. Pairwise permutation tests: female WT versus parthenogenic, P = 0.18; Sample sizes: female WT (N =32), parthenogenic (N = 40). Asterisks denote statistical significance: p < 0.05 (*), p < 0.01 (**), p < 0.001 (***).

**Supplementary Fig. S14.**
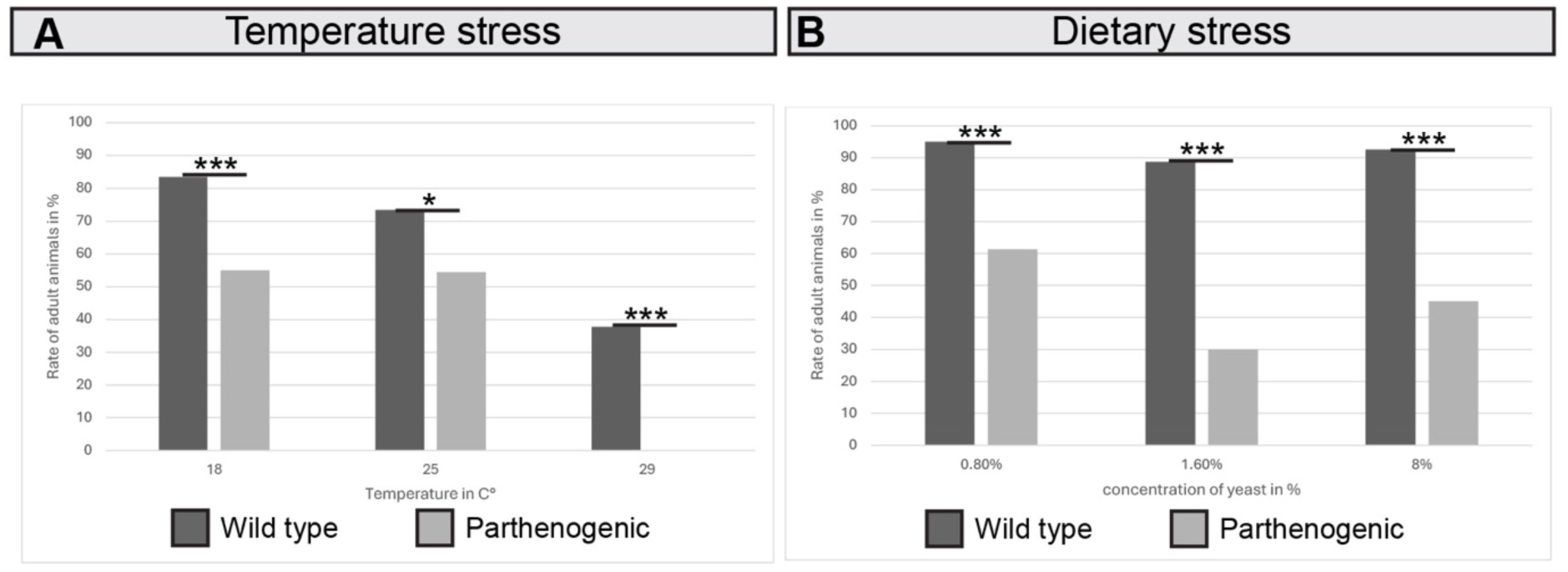
Parthenogenic *Drosophila mercatorum* show reduced developmental survival across environmental conditions, with complete lethality at high temperature. Wild-type and parthenogenic Brazilian flies were compared for developmental survival, quantified as the proportion of larvae that emerged as adults under different environmental conditions. **(A)** Developmental survival across three temperatures. Wild-type flies showed significantly higher survival than parthenogenic flies at all tested temperatures (18 °C, P = 0.0001; 25 °C, P = 0.019; 29 °C, P = 2.6 × 10⁻⁹). At 29 °C, no parthenogenic adults emerged. Sample sizes: wild type, N = 90, 90, 90; parthenogenic, N = 80, 70, 80. **(B)** Developmental survival across diets differing in yeast concentration. Wild-type flies again showed significantly higher survival than parthenogenic flies at all tested yeast concentrations (0.8%, P = 0.0001; 1.6% and 8%, P < 0.05 for both comparisons). Sample sizes: wild type, N = 80, 80, 80; parthenogenic, N = 80, 80, 80. The data were analyzed using a two-sample test for equality of proportions with continuity correction. Significance levels are denoted as follows: p < 0.05 (*), p < 0.01 (**), p < 0.001 (***).

**Supplementary Fig. S15:**
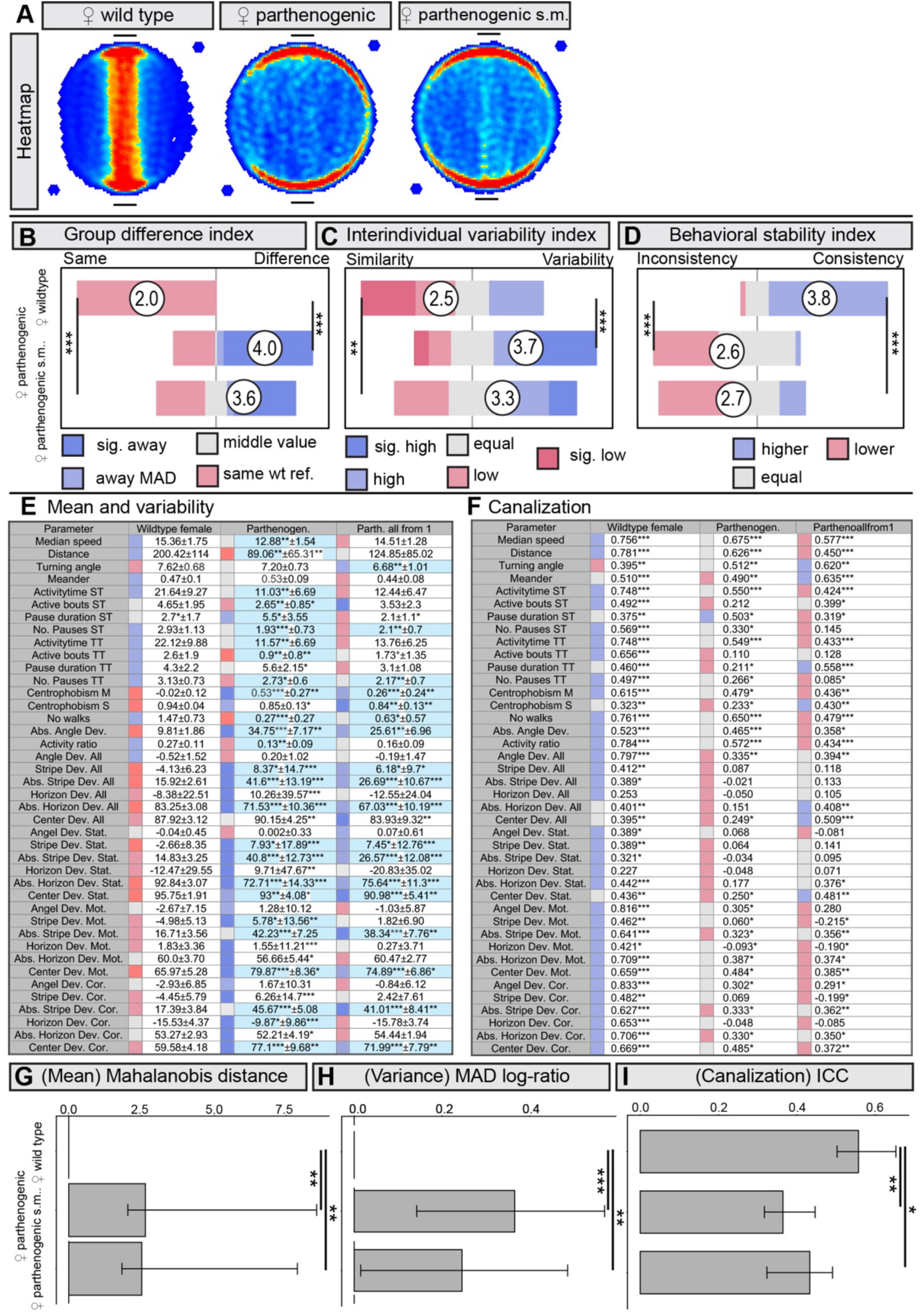
Parthenogenic offspring derived from a single mother recapitulate the Buridan phenotype of the parthenogenic stock. Buridan analysis shows that parthenogenic flies from the stock population and those derived from a single mother are highly similar to one another and differ markedly from wild-type controls. **(A)** Mean occupancy heatmaps show reduced stripe fixation in both parthenogenic groups relative to wild-type females **(B) Mean-difference index** across 41 behavioral parameters shows broad changes in behavioral means in both parthenogenic groups relative to wild-type controls. Full parameter-level data are shown in e. Pairwise Wilcoxon tests with Benjamini-Hochberg correction: female WT versus parthenogenic, P = 4.3 × 10⁻¹¹; female WT versus parthenogenic single-mother, P = 6.1 × 10⁻¹⁰; parthenogenic versus parthenogenic single-mother, P = 0.2. **(C) Variability index** based on the median absolute deviation (MAD) across the 41 behavioral parameters indicates increased interindividual behavioral variation in both parthenogenic groups relative to wild-type controls. Full parameter-level data are shown in e. Pairwise Wilcoxon tests with Benjamini-Hochberg correction: female WT versus parthenogenic, P = 6.3 × 10⁻⁵; female WT versus parthenogenic single-mother, P = 0.004; parthenogenic versus parthenogenic single-mother, P = 0.06. **(D) Behavioral consistency (canalization) index** across the 41 Buridan parameters shows reduced day-to-day behavioral consistency in both parthenogenic groups. Full parameter-level data are shown in f. Pairwise Wilcoxon tests with Benjamini-Hochberg correction: female WT versus parthenogenic, P = 2.1 × 10⁻¹²; female WT versus parthenogenic single-mother, P = 1.6 × 10⁻⁹; parthenogenic versus parthenogenic single-mother, P = 0.9. **(E)** Parameter-level analysis of mean and dispersion across all 41 Buridan parameters. The table shows group medians, with significance assessed by pairwise Wilcoxon tests with Benjamini-Hochberg correction, and MAD values, with significance assessed by Levene’s test. Colors denote relative MAD values, from low (blue) to high (red), and darker colors indicate parameters that differ significantly from both comparison groups. **(F)** Parameter-level analysis of day-to-day behavioral consistency across 3 testing days for all 41 Buridan parameters. The table shows Pearson correlation coefficients for all pairwise day comparisons after Fisher’s z-transformation. Asterisks denote significant correlations across the three-day pair comparisons. **(G-I)** Reanalysis of the same behavioral data shown in (E-F). **(G)** Mahalanobis distance indicates significant multivariate changes in behavioral means between wild-type females and both parthenogenic groups, whereas the two parthenogenic groups do not differ from one another. Pairwise permutation tests with Benjamini-Hochberg correction: female WT versus parthenogenic, P = 0.002; female WT versus parthenogenic single-mother, P = 0.007; parthenogenic versus parthenogenic single-mother, P = 0.77. **(H)** MAD log-ratio confirms increased overall interindividual behavioral variation in both parthenogenic groups relative to wild-type controls, with no significant difference between the two parthenogenic groups. Pairwise permutation tests with Benjamini-Hochberg correction: female WT versus parthenogenic, P < 0.001; female WT versus parthenogenic single-mother, P = 0.003; parthenogenic versus parthenogenic single-mother, P = 0.14. **(I)** Intraclass correlation coefficient (ICC) analysis confirms reduced temporal behavioral consistency in both parthenogenic groups relative to wild-type controls, with no significant difference between the two parthenogenic groups. Pairwise permutation tests with Benjamini-Hochberg correction: female WT versus parthenogenic, P = 0.003; female WT versus parthenogenic single-mother, P = 0.036; parthenogenic versus parthenogenic single-mother, P = 0.29. Sample sizes: female WT (N = 33), parthenogenic (N = 31), parthenogenic single-mother (N = 28). Asterisks denote statistical significance: p < 0.05 (*), p < 0.01 (**), p < 0.001 (***). Only in (F), asterisks indicate the number of significant comparisons across days.

**Supplementary Fig. S16:**
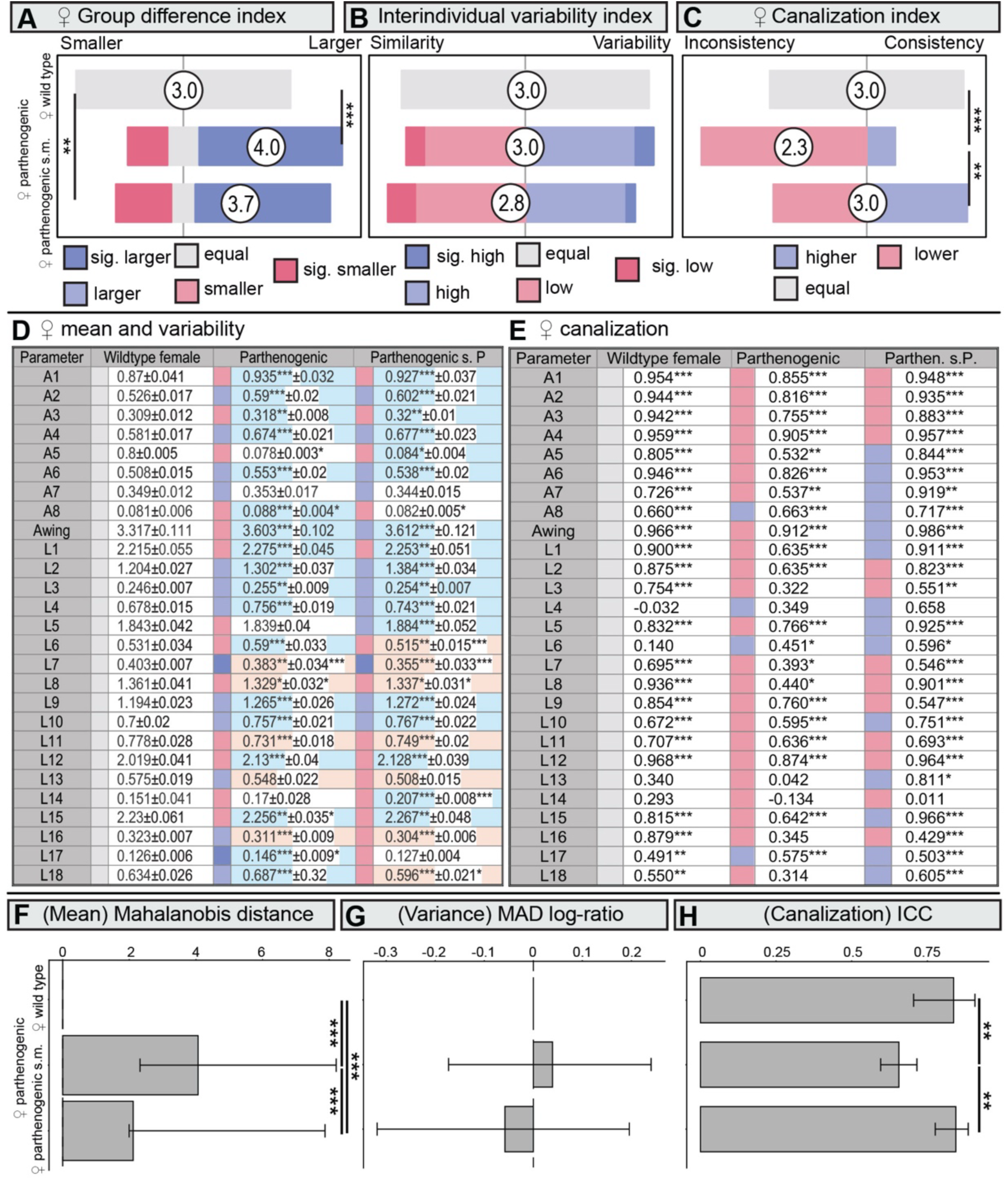
Parthenogenic offspring derived from a single mother largely recapitulate the wing phenotype of the parthenogenic stock. **(A) Mean-difference index** across 27 wing parameters shows broad changes in wing morphology in both parthenogenic groups relative to wild-type females. In contrast, the stock parthenogenic and single-mother parthenogenic groups do not differ from one another. Pairwise Wilcoxon tests with Benjamini-Hochberg correction: female WT versus parthenogenic, P = 0.0006; female WT versus parthenogenic single-mother line, P = 0.0091; parthenogenic versus parthenogenic single-mother line, P = 0.69. **(B) Variability index** based on the median absolute deviation (MAD) across the 27 wing parameters indicates no significant difference in overall interindividual wing variation between groups. Pairwise Wilcoxon tests with Benjamini-Hochberg correction: female WT versus parthenogenic, P = 0.81; female WT versus parthenogenic single-mother line, P = 0.45; parthenogenic versus parthenogenic single-mother line, P = 0.5. **(C) Canalization index** based on left-right correspondence across the 27 wing parameters shows reduced developmental stability in the parthenogenic stock, but not in the single-mother parthenogenic line. Pairwise Wilcoxon tests with Benjamini-Hochberg correction: female WT versus parthenogenic, P = 0.02; female WT versus parthenogenic single-mother line, P = 0.9; parthenogenic versus parthenogenic single-mother line, P = 0.02. **(D)** Parameter-level analysis of mean and dispersion across all 27 wing parameters. The table shows group medians, with significance assessed by pairwise Wilcoxon tests with Benjamini-Hochberg correction, and MAD values, with significance assessed by Levene’s test. Colors denote relative MAD values, from low (blue) to high (red), and darker colors indicate parameters that differ significantly between the two comparison groups. **(E)** Parameter-level analysis of left-right correspondence across the 27 wing parameters. The table shows Pearson correlation coefficients for right-left wing correspondence, with asterisks denoting statistical significance. Colors denote correlation strength, from low (red) to high (blue), with intermediate values in grey. **(F-H)** Reanalysis of the same wing anatomical data shown in (D-E). **(F)** Mahalanobis distance indicates significant multivariate changes in wing morphology between wild-type females and both parthenogenic groups, as well as between the two parthenogenic groups. Pairwise permutation tests with Benjamini-Hochberg correction: female WT versus parthenogenic, P < 0.001; female WT versus parthenogenic single-mother line, P < 0.001; parthenogenic versus parthenogenic single-mother line, P < 0.001. **(G)** MAD log-ratio indicates no significant difference in overall interindividual wing variation between groups. Pairwise permutation tests with Benjamini-Hochberg correction: female WT versus parthenogenic, P = 0.69; female WT versus parthenogenic single-mother line, P = 0.69; parthenogenic versus parthenogenic single-mother line, P = 0.69. **(H)** Intraclass correlation coefficient (ICC) analysis indicates reduced left-right correspondence in the parthenogenic stock relative to wild-type controls and the single-mother line. In contrast, wild-type and single-mother flies do not differ. Pairwise permutation tests with Benjamini-Hochberg correction: female WT versus parthenogenic, P = 0.008; female WT versus parthenogenic single-mother line, P = 0.89; parthenogenic versus parthenogenic single-mother line, P = 0.008. Sample sizes: female WT (N = 33), parthenogenic (N = 31), parthenogenic single-mother (N = 28). Asterisks denote statistical significance: p < 0.05 (*), p < 0.01 (**), p < 0.001 (***). Only in (E), asterisks indicate the number of significant comparisons across days.

**Supplementary Fig. S17:**
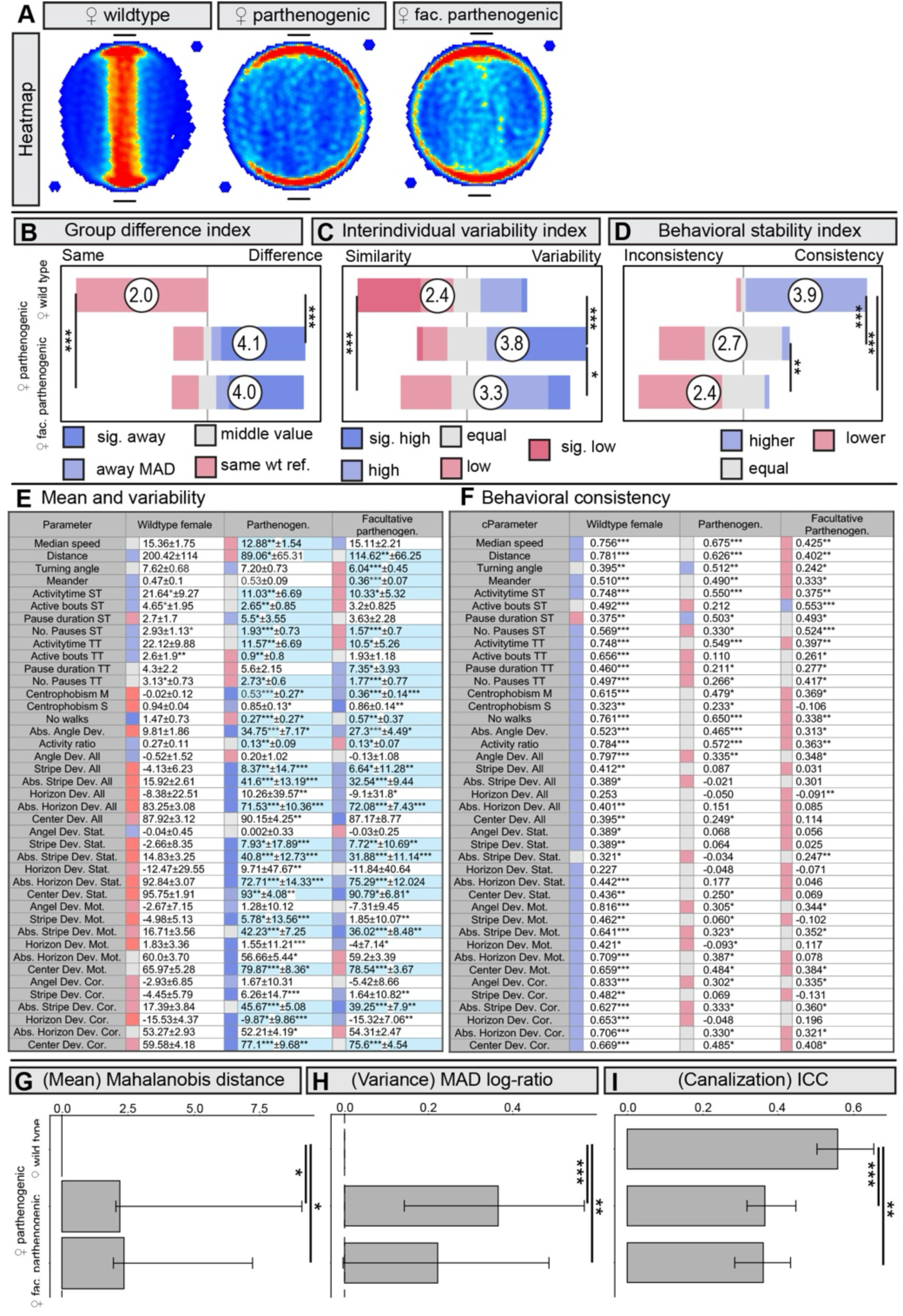
A facultative parthenogenic stock recapitulates the Buridan phenotype of the parthenogenic stocks. Buridan analysis shows that obligately parthenogenic flies and facultatively parthenogenic flies are highly similar to one another and differ markedly from wild-type controls. **(A)** Mean occupancy heatmaps show reduced stripe fixation in both parthenogenic groups relative to wild-type females. **(B) Mean-difference index** across 41 behavioral parameters shows broad changes in behavioral means in both parthenogenic groups relative to wild-type controls. Full parameter-level data are shown in e. Pairwise Wilcoxon tests with Benjamini-Hochberg correction: female WT versus parthenogenic, P = 1.6 × 10⁻¹²; female WT versus facultative parthenogenic, P = 7.4 × 10⁻¹³; parthenogenic versus facultative parthenogenic, P = 0.7. **(C) Variability index** based on the median absolute deviation (MAD) across the 41 behavioral parameters indicates increased interindividual behavioral variation in both parthenogenic groups relative to wild-type controls. Full parameter-level data are shown in e. Pairwise Wilcoxon tests with Benjamini-Hochberg correction: female WT versus parthenogenic, P = 4.5 × 10⁻⁶; female WT versus facultative parthenogenic, P = 0.0005; parthenogenic versus facultative parthenogenic, P = 0.04. **(D) Behavioral consistency (canalization) index** across the 41 Buridan parameters shows reduced day-to-day behavioral consistency in both parthenogenic groups. Full parameter-level data are shown in f. Pairwise Wilcoxon tests with Benjamini-Hochberg correction: female WT versus parthenogenic, P = 8.2 × 10⁻¹⁴; female WT versus facultative parthenogenic, P = 5.3 × 10⁻¹⁵; parthenogenic versus facultative parthenogenic, P = 0.009. **(E)** Parameter-level analysis of mean and dispersion across all 41 Buridan parameters. The table shows group medians, with significance assessed by pairwise Wilcoxon tests with Benjamini-Hochberg correction, and MAD values, with significance assessed by Levene’s test. Colors denote relative MAD values, from low (blue) to high (red), and darker colors indicate parameters that differ significantly between the two comparison groups. **(F)** Parameter-level analysis of day-to-day behavioral consistency across 3 testing days for all 41 Buridan parameters. The table shows Pearson correlation coefficients for all pairwise day comparisons after Fisher’s z-transformation. Asterisks denote significant correlations across the three-day pair comparisons. **(G-I)** Reanalysis of the same behavioral data as in (E,F). **(G)** Mahalanobis distance with pairwise permutation tests and Benjamini-Hochberg correction: female WT versus parthenogenic, P = 0.038; female WT versus facultative parthenogenic, P = 0.021; parthenogenic versus facultative parthenogenic, P = 0.73. **(H)** MAD log-ratio with pairwise permutation tests and Benjamini-Hochberg correction: female WT versus parthenogenic, P < 0.001; female WT versus facultative parthenogenic, P = 0.005; parthenogenic versus facultative parthenogenic, P = 0.07. **(I)** ICC with pairwise permutation tests and Benjamini-Hochberg correction: female WT versus parthenogenic, P < 0.001; female WT versus facultative parthenogenic, P = 0.005; parthenogenic versus facultative parthenogenic, P = 0.95. Sample sizes: female WT (N = 33), parthenogenic (N = 31), facultative parthenogenic (N = 28). Asterisks denote statistical significance: p < 0.05 (*), p < 0.01 (**), p < 0.001 (***). Only in (F), asterisks indicate the number of significant comparisons across days.

**Supplementary Fig. S18:**
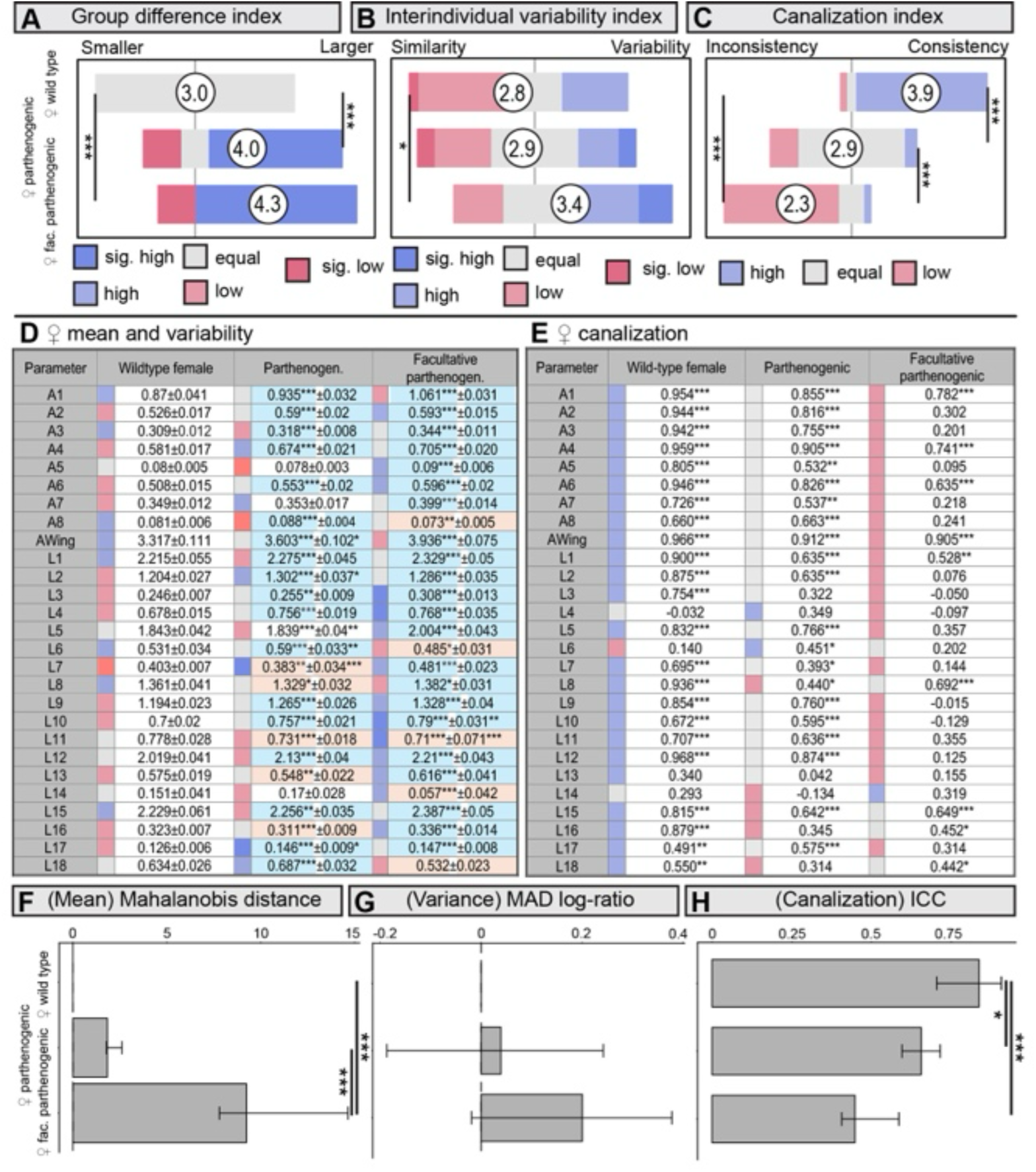
A facultative parthenogenic stock largely recapitulates the wing phenotype and reduced developmental stability of the parthenogenic stock. **(A) Mean-difference index** across 27 wing parameters shows broad changes in wing morphology in both parthenogenic groups relative to wild-type females. In contrast, the obligate and facultative parthenogenic groups do not differ significantly from one another. Full parameter-level data are shown in (D). Pairwise Wilcoxon tests with Benjamini-Hochberg correction: female WT versus parthenogenic, P = 0.0006; female WT versus facultative parthenogenic, P = 1.0 × 10⁻¹⁵; parthenogenic versus facultative parthenogenic, P = 0.3. **(B) Variability index** based on the median absolute deviation (MAD) across the 27 wing parameters indicates increased interindividual variation in the facultative parthenogenic group. In contrast, the obligate parthenogenic group does not differ significantly from wild-type females in this index. Full parameter-level data are shown in (D). Pairwise Wilcoxon tests with Benjamini-Hochberg correction: female WT versus parthenogenic, P = 0.7; female WT versus facultative parthenogenic, P = 0.04; parthenogenic versus facultative parthenogenic, P = 0.1. **(C) Canalization index** based on left-right correspondence across the 27 wing parameters confirms reduced developmental stability in both parthenogenic groups. Full parameter-level data are shown in (E). Pairwise Wilcoxon tests with Benjamini-Hochberg correction: female WT versus parthenogenic, P = 2.5 × 10⁻⁸; female WT versus facultative parthenogenic, P = 4.6 × 10⁻¹⁰; parthenogenic versus facultative parthenogenic, P = 4.1 × 10⁻⁵. **(D)** Parameter-level analysis of mean and dispersion across all 27 wing parameters in wild-type, parthenogenic, and facultative parthenogenic females. The table shows group medians, with significance assessed by pairwise Wilcoxon tests with Benjamini-Hochberg correction, and MAD values, with significance assessed by Levene’s test. Colors denote relative MAD values, from low (blue) to high (red), and darker colors indicate parameters that differ significantly from both comparison groups. **(E)** Parameter-level analysis of left-right correspondence across the 27 wing parameters. The table shows Pearson correlation coefficients for bilateral wing correspondence, with asterisks denoting statistical significance. Colors denote correlation strength, from low (red) to high (blue), with intermediate values in grey. **(F-H)** Reanalysis of the same wing anatomical data shown in (D-E). **(F)** Mahalanobis distance indicates no significant multivariate change between wild-type and obligate parthenogenic females, but a strong change in the facultative parthenogenic group relative to both other groups. Pairwise permutation tests with Benjamini-Hochberg correction: female WT versus parthenogenic, P = 0.21; female WT versus facultative parthenogenic, P < 0.001; parthenogenic versus facultative parthenogenic, P < 0.001. **(G)** MAD log-ratio indicates no significant difference in overall interindividual wing variation in this reanalysis. Pairwise permutation tests with Benjamini-Hochberg correction: female WT versus parthenogenic, P = 0.71; female WT versus facultative parthenogenic, P = 0.19; parthenogenic versus facultative parthenogenic, P = 0.19. **(H)** Intraclass correlation coefficient (ICC) analysis indicates reduced left-right correspondence in parthenogenic females relative to wild-type controls, with a stronger reduction in the facultative parthenogenic group. Pairwise permutation tests with Benjamini-Hochberg correction: female WT versus parthenogenic, P = 0.014; female WT versus facultative parthenogenic, P < 0.001; parthenogenic versus facultative parthenogenic, P = 0.084. Sample sizes: female WT (N = 29), parthenogenic (N = 31), facultative parthenogenic (N = 27). Asterisks denote statistical significance: p < 0.05 (*), p < 0.01 (**), p < 0.001 (***). Only in (E), asterisks indicate the number of significant comparisons across days.

**Supplementary Fig. S19:**
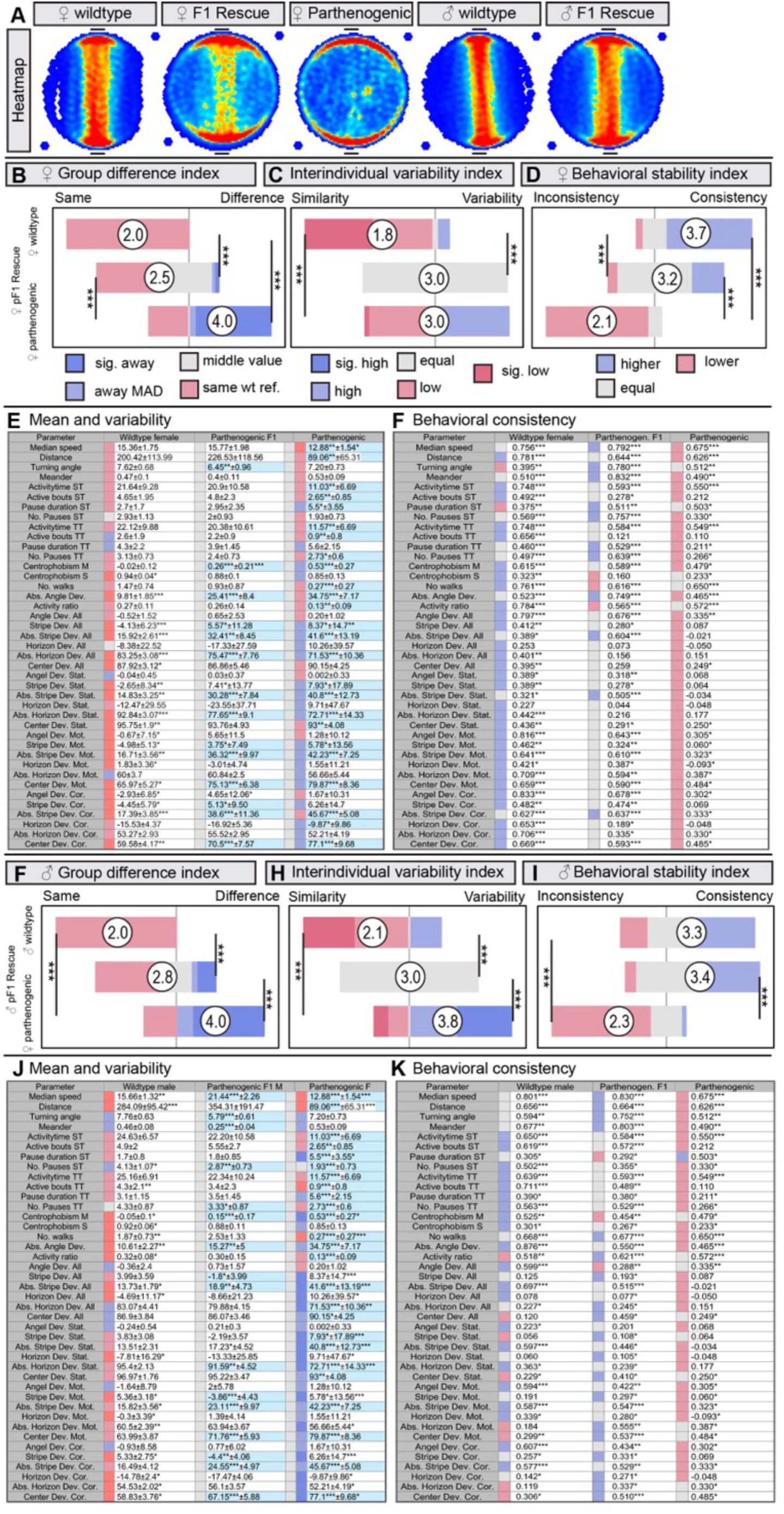
F1 hybrids from parthenogenic mothers partially rescue the Buridan phenotype and behavioral stability of parthenogenic flies. Buridan analysis shows that offspring from mating parthenogenic females with wild-type males exhibit behavioral phenotypes intermediate between wild-type and parthenogenic flies, consistent with partial restoration of heterozygosity and behavioral robustness. **(A)** Mean occupancy heatmaps show that female F_1_ rescue flies exhibit stripe fixation intermediate between wild-type and parthenogenic females. **(B) Mean-difference index** across 41 behavioral parameters shows that female F_1_ rescue flies are changeed away from both wild-type and parthenogenic groups. Full parameter-level data are shown in (E). Pairwise Wilcoxon tests with Benjamini-Hochberg correction: female WT versus female F_1_ rescue, P = 2.0 × 10⁻⁶; female WT versus parthenogenic, P = 1.4 × 10⁻¹⁰; parthenogenic versus female F_1_ rescue, P = 1.9 × 10⁻⁵. **(C) Variability index** based on the median absolute deviation (MAD) across the 41 behavioral parameters indicates increased interindividual behavioral variation in both parthenogenic and female F_1_ rescue flies relative to wild-type controls. In contrast, the two groups do not differ. Full parameter-level data are shown in (E). Pairwise Wilcoxon tests with Benjamini-Hochberg correction: female WT versus female F_1_ rescue, P = 1.2 × 10⁻¹⁰; female WT versus parthenogenic, P = 7.4 × 10⁻⁷; parthenogenic versus female F_1_ rescue, P = 0.8. **(D) Behavioral consistency (canalization) index** across the 41 Buridan parameters shows mildly reduced day-to-day behavioral consistency in female F_1_ rescue flies and strongly reduced consistency in parthenogenic flies. Full parameter-level data are shown in (F). Pairwise Wilcoxon tests with Benjamini-Hochberg correction: female WT versus female F_1_ rescue, P = 9.0 × 10⁻⁵; female WT versus parthenogenic, P = 7.5 × 10⁻¹⁵; parthenogenic versus female F_1_ rescue, P = 8.2 × 10⁻¹³. **(E)** Parameter-level analysis of mean and dispersion across all 41 Buridan parameters for female wild-type, female F_1_ rescue, and parthenogenic flies. The table shows group medians, with significance assessed by pairwise Wilcoxon tests with Benjamini-Hochberg correction, and MAD values, with significance assessed by Levene’s test. Colors denote relative MAD values, from low (blue) to high (red), and darker colors indicate parameters that differ significantly from both comparison groups. **(F)** Parameter-level analysis of day-to-day behavioral consistency across 3 testing days for all 41 Buridan parameters in female wild-type, female F_1_ rescue, and parthenogenic flies. The table shows Pearson correlation coefficients for all pairwise day comparisons after Fisher’s z-transformation. Asterisks denote significant correlations across the three-day pair comparisons. Colors denote correlation strength, from low (red) to high (blue), with intermediate values in grey. Sample sizes: female WT, n = 33; female F_1_ rescue, n = 29; parthenogenic, n = 31. **(G) Mean-difference index** shows that male F_1_ rescue flies are intermediate between wild-type males and parthenogenic flies across the 41 behavioral parameters. Full parameter-level data are shown in (J). Pairwise Wilcoxon tests with Benjamini-Hochberg correction: male WT versus male F_1_ rescue, P = 9.9 × 10⁻⁷; male WT versus parthenogenic, P = 1.8 × 10⁻¹¹; parthenogenic versus male F_1_ rescue, P = 9.3 × 10⁻⁵. **(H) Variability index** based on MAD indicates increased interindividual behavioral variation in both parthenogenic and male F_1_ rescue flies relative to wild-type males. Full parameter-level data are shown in (J). Pairwise Wilcoxon tests with Benjamini-Hochberg correction: male WT versus male F_1_ rescue, P = 5.9 × 10⁻⁶; male WT versus parthenogenic, P = 4.9 × 10⁻⁷; parthenogenic versus male F_1_ rescue, P = 3.9 × 10⁻⁵. **(I) Behavioral consistency (canalization) index** shows no reduction in day-to-day behavioral consistency in male F_1_ rescue flies relative to wild-type males, whereas parthenogenic flies remain strongly impaired. Full parameter-level data are shown in (K). Pairwise Wilcoxon tests with Benjamini-Hochberg correction: male WT versus male F_1_ rescue, P = 0.9; male WT versus parthenogenic, P = 4.0 × 10⁻⁸; parthenogenic versus male F_1_ rescue, P = 2.5 × 10⁻¹⁰. **(J)** Parameter-level analysis of mean and dispersion across all 41 Buridan parameters for male wild-type, male F_1_ rescue, and parthenogenic flies. The table shows group medians, with significance assessed by pairwise Wilcoxon tests with Benjamini-Hochberg correction, and MAD values, with significance assessed by Levene’s test. Colors denote relative MAD values, from low (blue) to high (red), and darker colors indicate parameters that differ significantly from both comparison groups. **(K)** Parameter-level analysis of day-to-day behavioral consistency across 3 testing days for all 41 Buridan parameters in male wild-type, male F_1_ rescue, and parthenogenic flies. The table shows Pearson correlation coefficients for all pairwise day comparisons after Fisher’s z-transformation. Asterisks denote significant correlations across the three-day pair comparisons. Colors denote correlation strength, from low (red) to high (blue), with intermediate values in grey. Sample sizes: male WT (N = 31), male F_1_ rescue (N = 31), parthenogenic (N = 31). Asterisks denote statistical significance: p < 0.05 (*), p < 0.01 (**), p < 0.001 (***). Only in (F) and (K) do asterisks refer to the number of significant comparisons between different days.

**Supplementary Fig. S20:**
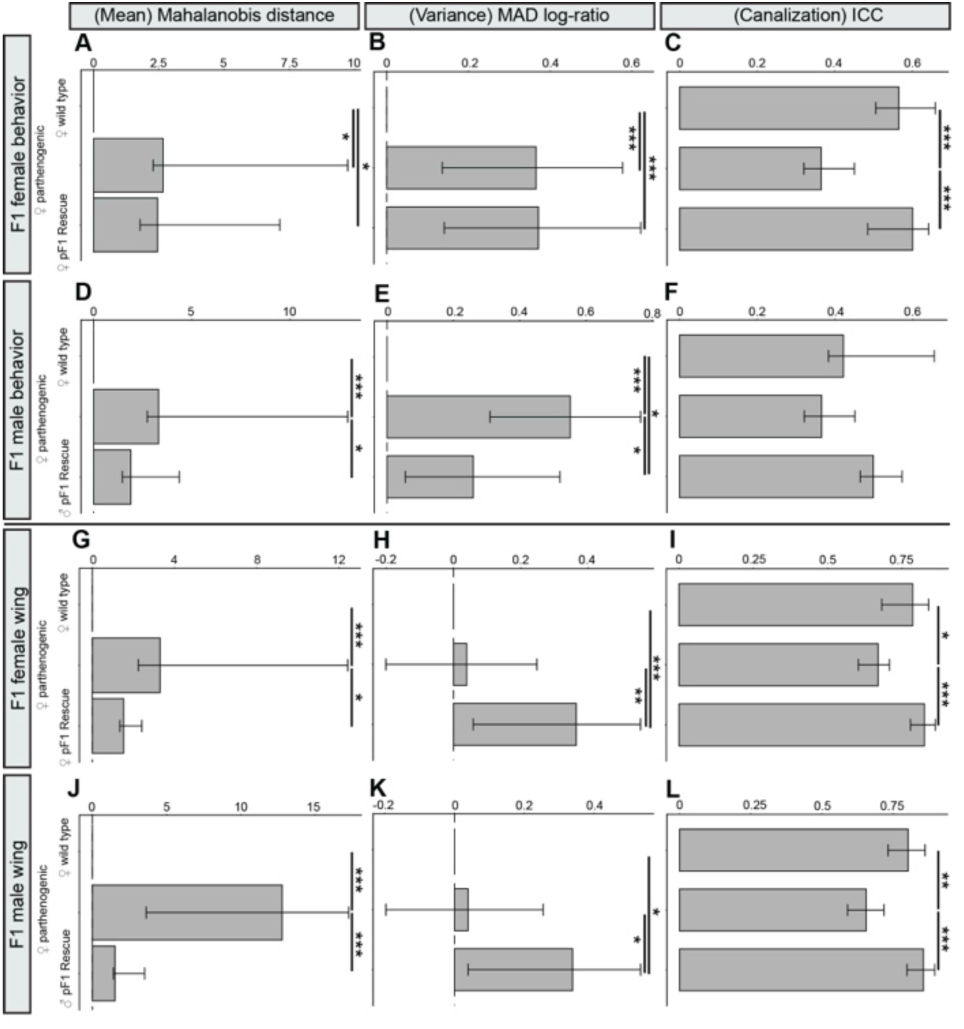
Reanalysis of F_1_ rescue data shows partial restoration of behavioral and anatomical robustness in offspring of parthenogenic mothers. Buridan and wing data from Supplementary Figs. S19 and S21 were reanalyzed using Mahalanobis distance, MAD log-ratio, and the intraclass correlation coefficient (ICC). **(A-C)** Buridan data from female F_1_ rescue flies. **(A)** Mahalanobis distance indicates significant multivariate changes in behavioral means between wild-type females and both parthenogenic and female F_1_ rescue flies, whereas parthenogenic and female F_1_ rescue flies do not differ. Pairwise permutation tests with Benjamini-Hochberg correction: female WT versus parthenogenic, P = 0.016; female WT versus female pF1, P = 0.027; parthenogenic versus female pF1, P = 0.67. **(B)** MAD log-ratio indicates increased overall interindividual behavioral variation in both parthenogenic and female F_1_ rescue flies relative to wild-type females, with no difference between the two. Pairwise permutation tests with Benjamini-Hochberg correction: female WT versus parthenogenic, P < 0.001; female WT versus female pF1, P < 0.001; parthenogenic versus female pF1, P = 0.97. **(C)** ICC analysis indicates reduced temporal behavioral consistency in parthenogenic flies relative to both wild-type and female F_1_ rescue flies, whereas wild-type and female F_1_ rescue flies do not differ. Pairwise permutation tests with Benjamini-Hochberg correction: female WT versus parthenogenic, P < 0.001; female WT versus female pF1, P = 0.51; parthenogenic versus female pF1, P < 0.001. **(D-F)** Buridan data from male F_1_ rescue flies. **(D)** Mahalanobis distance indicates a strong multivariate change between male wild-type and parthenogenic flies, while male F_1_ rescue flies are intermediate and do not differ significantly from male wild-type controls. Pairwise permutation tests with Benjamini-Hochberg correction: male WT versus parthenogenic, P < 0.001; male WT versus male pF1, P = 0.097; parthenogenic versus male pF1, P = 0.02. **(E)** MAD log-ratio indicates increased behavioral variation in both parthenogenic and male F_1_ rescue flies relative to male wild-type controls. Pairwise permutation tests with Benjamini-Hochberg correction: male WT versus parthenogenic, P < 0.001; male WT versus male pF1, P = 0.029; parthenogenic versus male pF1, P = 0.024. **(F)** ICC analysis shows no significant difference in temporal behavioral consistency among male wild-type, parthenogenic, and male F_1_ rescue flies. Pairwise permutation tests with Benjamini-Hochberg correction: male WT versus parthenogenic, P = 0.38; male WT versus male pF1, P = 0.34; parthenogenic versus male pF1, P = 0.1. Sample sizes: female WT, n = 33; female F_1_ rescue, n = 29; parthenogenic, n = 31; male WT, n = 31; male F_1_ rescue, n = 31. **(G-I)** Wing anatomical data from female F_1_ rescue flies. **(G)** Mahalanobis distance indicates a strong multivariate change in wing anatomy between wild-type and parthenogenic females. In contrast, female F_1_ rescue flies do not differ significantly from wild type and differ modestly from parthenogenic flies. Pairwise permutation tests with Benjamini-Hochberg correction: female WT versus parthenogenic, P < 0.001; female WT versus female pF1, P = 0.08; parthenogenic versus female pF1, P = 0.011. **(H)** MAD log-ratio indicates no difference in overall wing variation between female wild-type and parthenogenic flies, but increased variation in female F_1_ rescue flies relative to both groups. Pairwise permutation tests with Benjamini-Hochberg correction: female WT versus parthenogenic, P = 0.7; female WT versus female pF1, P < 0.001; parthenogenic versus female pF1, P = 0.002. **(I)** ICC analysis indicates reduced left-right correspondence in parthenogenic females relative to wild-type controls, whereas female F_1_ rescue flies are similar to wild type and differ from parthenogenic flies. Pairwise permutation tests with Benjamini-Hochberg correction: female WT versus parthenogenic, P = 0.018; female WT versus female pF1, P = 0.43; parthenogenic versus female pF1, P < 0.001. **(J-L)** Wing anatomical data from male F_1_ rescue flies. **(J)** Mahalanobis distance indicates a strong multivariate change between male wild-type and parthenogenic flies. In contrast, male F_1_ rescue flies do not differ from wild-type controls and differ strongly from parthenogenic flies. Pairwise permutation tests with Benjamini-Hochberg correction: male WT versus parthenogenic, P < 0.001; male WT versus male pF1, P = 0.61; parthenogenic versus male pF1, P < 0.001. **(K)** MAD log-ratio indicates increased wing variation in male F_1_ rescue flies relative to both male wild-type and parthenogenic flies. Pairwise permutation tests with Benjamini-Hochberg correction: male WT versus parthenogenic, P = 0.75; male WT versus male pF1, P = 0.039; parthenogenic versus male pF1, P = 0.045. **(L)** ICC analysis indicates reduced left-right correspondence in parthenogenic males relative to wild- type controls, whereas male F_1_ rescue flies are similar to wild-type and differ from parthenogenic flies. Pairwise permutation tests with Benjamini-Hochberg correction: male WT versus parthenogenic, P = 0.009; male WT versus male pF1, P = 0.13; parthenogenic versus male pF1, P < 0.001. Sample sizes: female WT (N = 29), female F_1_ Rescue (N = 26), parthenogenic (N = 31), male WT (N = 33), male F_1_ Rescue (N = 28). Asterisks denote statistical significance: p < 0.05 (*), p < 0.01 (**), p < 0.001 (***).

**Supplementary Fig. S21:**
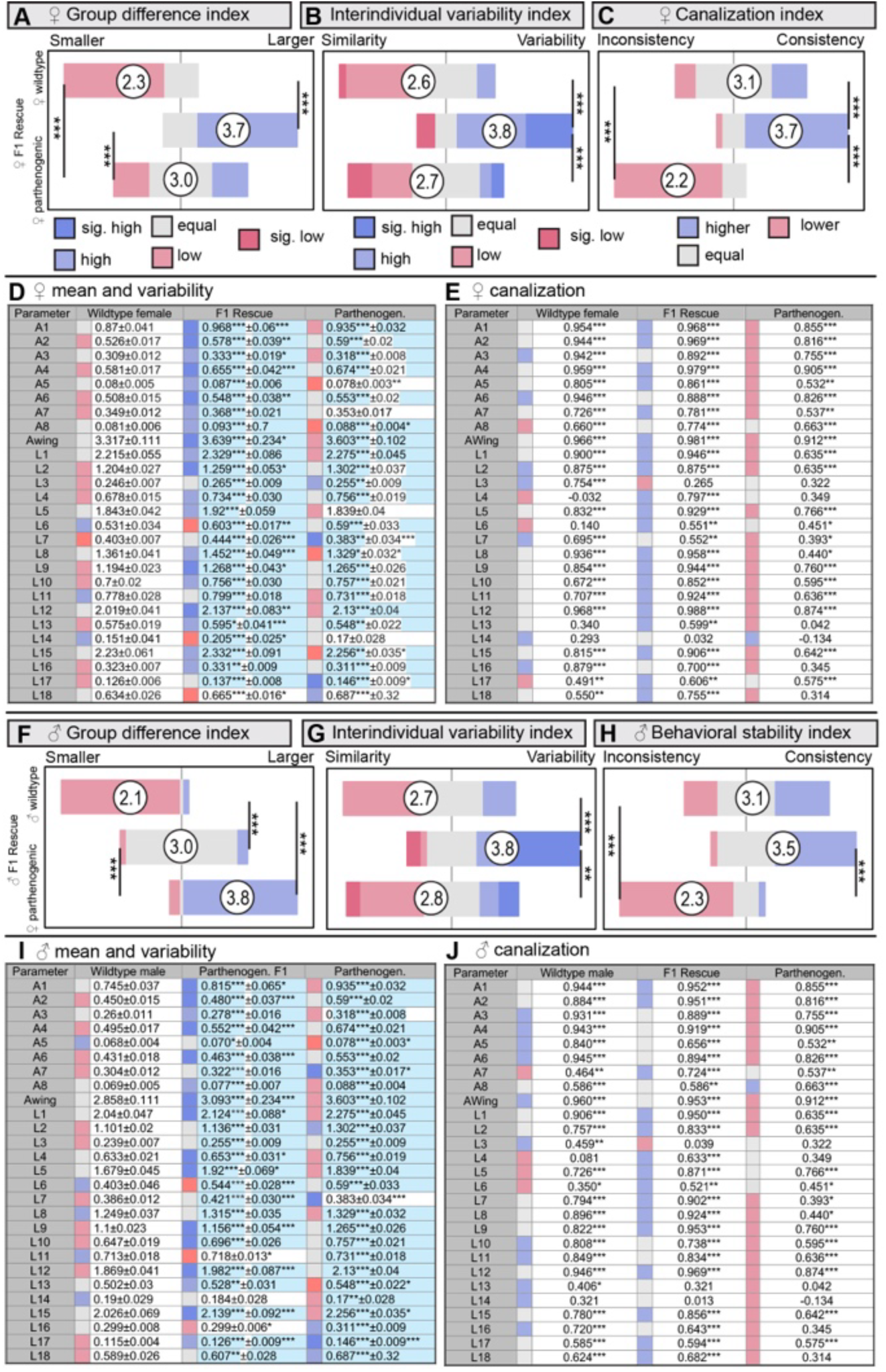
F1 rescue alters wing anatomy, increases interindividual variation, and restores developmental stability. (A) **Mean-difference index** across 27 wing parameters shows broad changes in wing morphology in female F_1_ rescue flies relative to both wild-type and parthenogenic females. F_1_ rescue flies have the largest wings among the three groups. Full parameter-level data are shown in (D). Pairwise Wilcoxon tests with Benjamini-Hochberg correction: female WT versus F_1_ rescue, P = 3.9 × 10⁻¹⁰; female WT versus parthenogenic, P = 0.0001; F_1_ rescue versus parthenogenic, P = 0.0001. (B) **Variability index** based on the median absolute deviation (MAD) across the 27 wing parameters indicates increased interindividual variation in F_1_ rescue flies, whereas wild-type and parthenogenic females do not differ. Full parameter-level data are shown in (D). Pairwise Wilcoxon tests with Benjamini-Hochberg correction: female WT versus F_1_ rescue, P = 0.0004; female WT versus parthenogenic, P = 0.9; F_1_ rescue versus parthenogenic, P = 0.0003. (C) **Canalization index** based on left-right correspondence across the 27 wing parameters indicates increased developmental stability in F_1_ rescue flies relative to both wild-type and parthenogenic females. Full parameter-level data are shown in (E). Pairwise Wilcoxon tests with Benjamini-Hochberg correction: female WT versus F_1_ rescue, P = 0.0002; female WT versus parthenogenic, P = 7.6 × 10⁻⁷; F_1_ rescue versus parthenogenic, P = 3.7 × 10⁻¹⁰. (D) Parameter-level analysis of mean and dispersion across all 27 wing parameters of female wild-type, female F_1_ rescue, and parthenogenic flies. The table shows group medians, with significance assessed by pairwise Wilcoxon tests with Benjamini-Hochberg correction, and MAD values, with significance assessed by Levene’s test. Colors denote relative MAD values, from low (blue) to high (red), and darker colors indicate parameters that differ significantly from both comparison groups. (E) Parameter-level analysis of right-left correspondence across the 27 wing parameters of female wild-type, female F_1_ rescue, and parthenogenic flies. The table shows Pearson correlation coefficients for bilateral wing correspondence, with asterisks denoting statistical significance. Colors denote correlation strength, from low (red) to high (blue), with intermediate values in grey. Sample sizes: female WT (N = 29), female F_1_ rescue (N = 26), parthenogenic (N = 31). (F) **Mean-difference index** shows that male F_1_ rescue flies differ significantly from both male wild-type controls and parthenogenic flies across the 27 wing parameters. Male F_1_ rescue flies have larger wings than wild-type males but smaller wings than parthenogenic females. Full parameter-level data are shown in (I). Pairwise Wilcoxon tests with Benjamini-Hochberg correction: male WT versus male F_1_ rescue, P = 1.2 × 10⁻⁹; male WT versus parthenogenic, P = 2.7 × 10⁻¹⁰; male F_1_ rescue versus parthenogenic, P = 1.1 × 10⁻⁷. (G) **Variability index** based on MAD indicates increased interindividual variation in male F_1_ rescue flies across the 27 wing parameters. Full parameter-level data are shown in (I). Pairwise Wilcoxon tests with Benjamini-Hochberg correction: male WT versus male F_1_ rescue, P = 3.4 × 10⁻⁵; male WT versus parthenogenic, P = 0.8; male F_1_ rescue versus parthenogenic, P = 0.003. (H) **Canalization index** indicates increased developmental stability in male F_1_ rescue flies relative to parthenogenic flies. Full parameter-level data are shown in (J). Pairwise Wilcoxon tests with Benjamini-Hochberg correction: male WT versus male F_1_ rescue, P = 0.07; male WT versus parthenogenic, P = 2.9 × 10⁻⁵; male F_1_ rescue versus parthenogenic, P = 2.7 × 10⁻⁸. (I) Parameter-level analysis of mean and dispersion across all 27 wing parameters for male wild-type, male F_1_ rescue, and parthenogenic flies. The table shows group medians, with significance assessed by pairwise Wilcoxon tests with Benjamini-Hochberg correction, and MAD values, with significance assessed by Levene’s test. Colors denote relative MAD values, from low (blue) to high (red), and darker colors indicate parameters that differ significantly from both comparison groups. (J) Parameter-level analysis of right-left correspondence across the 27 wing parameters for male wild-type, male F_1_ rescue, and parthenogenic flies. The table shows Pearson correlation coefficients for bilateral wing correspondence, with asterisks denoting statistical significance. Colors denote correlation strength, from low (red) to high (blue), with intermediate values in grey. Sample sizes: male WT (N = 33), male F_1_ rescue (N = 28), parthenogenic (N = 31). Asterisks denote statistical significance: p < 0.05 (*), p < 0.01 (**), p < 0.001 (***). Only in (E) and (J) do asterisks refer to the number of significant comparisons between different days.

**Supplementary Fig. S22:**
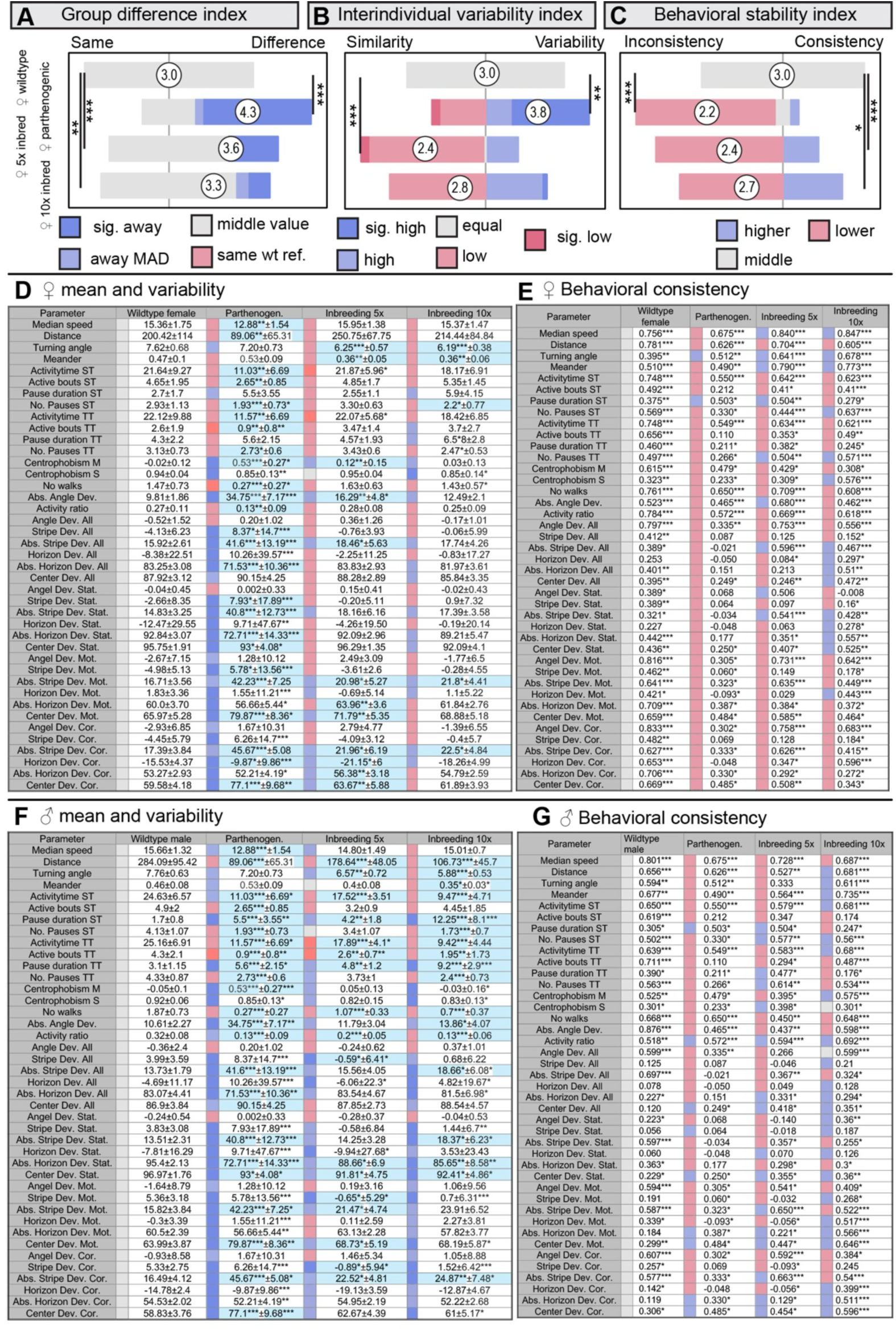
Inbreeding in Brazilian *Drosophila mercatorum* alters mean behavior and behavioral canalization, with weaker effects on overall interindividual variation. Buridan analysis shows that 5 and 10 generations of inbreeding affect behavioral means and day-to-day behavioral consistency, whereas effects on overall interindividual variation are comparatively modest. **(A) Mean-difference index** across 41 behavioral parameters shows that the 5× and 10× inbred lines are intermediate between wild-type and parthenogenic females. Full parameter-level data are shown in (D). Pairwise Wilcoxon tests with Benjamini-Hochberg correction: female WT versus parthenogenic, P = 1.2 × 10⁻¹⁰; female WT versus 5× inbred, P = 2.0 × 10⁻⁴; female WT versus 10× inbred, P = 0.003. **(B) Variability index** based on the median absolute deviation (MAD) across the 41 behavioral parameters indicates increased interindividual variation in parthenogenic females and reduced variation in the 5× inbred line relative to wild-type females. Full parameter-level data are shown in (D). Pairwise Wilcoxon tests with Benjamini-Hochberg correction: female WT versus parthenogenic, P = 0.008; female WT versus 5× inbred, P = 4.5 × 10⁻⁷; female WT versus 10× inbred, P = 0.06. **(C) Behavioral consistency (canalization) index** across the 41 tested behavioral parameters shows reduced day-to-day behavioral consistency in parthenogenic and inbred females. Full parameter-level data are shown in (E). Pairwise Wilcoxon tests with Benjamini-Hochberg correction: female WT versus parthenogenic, P = 5.8 × 10⁻¹³; female WT versus 5× inbred, P = 1.3 × 10⁻⁶; female WT versus 10× inbred, P = 0.02. **(D)** Parameter-level analysis of mean and dispersion across all 41 Buridan parameters for female wild-type, parthenogenic, 5× inbred, and 10× inbred flies. The table shows group medians, with significance assessed by pairwise Wilcoxon tests with Benjamini-Hochberg correction, and MAD values, with significance assessed by Levene’s test. Colors denote relative MAD values, from low (blue) to high (red), and darker colors indicate parameters that differ significantly from both comparison groups. **(E)** Parameter-level analysis of day-to-day behavioral consistency across 3 testing days for all 41 Buridan parameters in female wild-type, parthenogenic, 5× inbred, and 10× inbred flies. The table shows Pearson correlation coefficients for all pairwise day comparisons after Fisher’s z-transformation. Asterisks denote significant correlations across the three-day pair comparisons. Colors denote correlation strength, from low (red) to high (blue), with intermediate values in grey. Sample sizes: female WT, n = 33; parthenogenic, n = 31; female 5× inbred, n = 24; female 10× inbred, n = 30. **(F)** Parameter-level analysis of mean and dispersion across all 41 Buridan parameters for male wild-type, parthenogenic, 5× inbred, and 10× inbred flies. The table shows group medians, with significance assessed by pairwise Wilcoxon tests with Benjamini-Hochberg correction, and MAD values, with significance assessed by Levene’s test. Colors denote relative MAD values, from low (blue) to high (red), and darker colors indicate parameters that differ significantly from both comparison groups. **(G)** Parameter-level analysis of day-to-day behavioral consistency across 3 testing days for all 41 Buridan parameters in male wild-type, parthenogenic, 5× inbred, and 10× inbred flies. The table shows Pearson correlation coefficients for all pairwise day comparisons after Fisher’s z-transformation. Asterisks denote significant correlations across the three-day pair comparisons. Colors denote correlation strength, from low (red) to high (blue), with intermediate values in grey. Sample sizes: male WT (N = 31), parthenogenic (N = 31), male 5x inbred (N = 23), male 10x inbred (N = 30). Asterisks denote statistical significance: p < 0.05 (*), p < 0.01 (**), p < 0.001 (***). Only in (E) and (G) do asterisks indicate the number of significant comparisons between different days.

**Supplementary Fig. S23:**
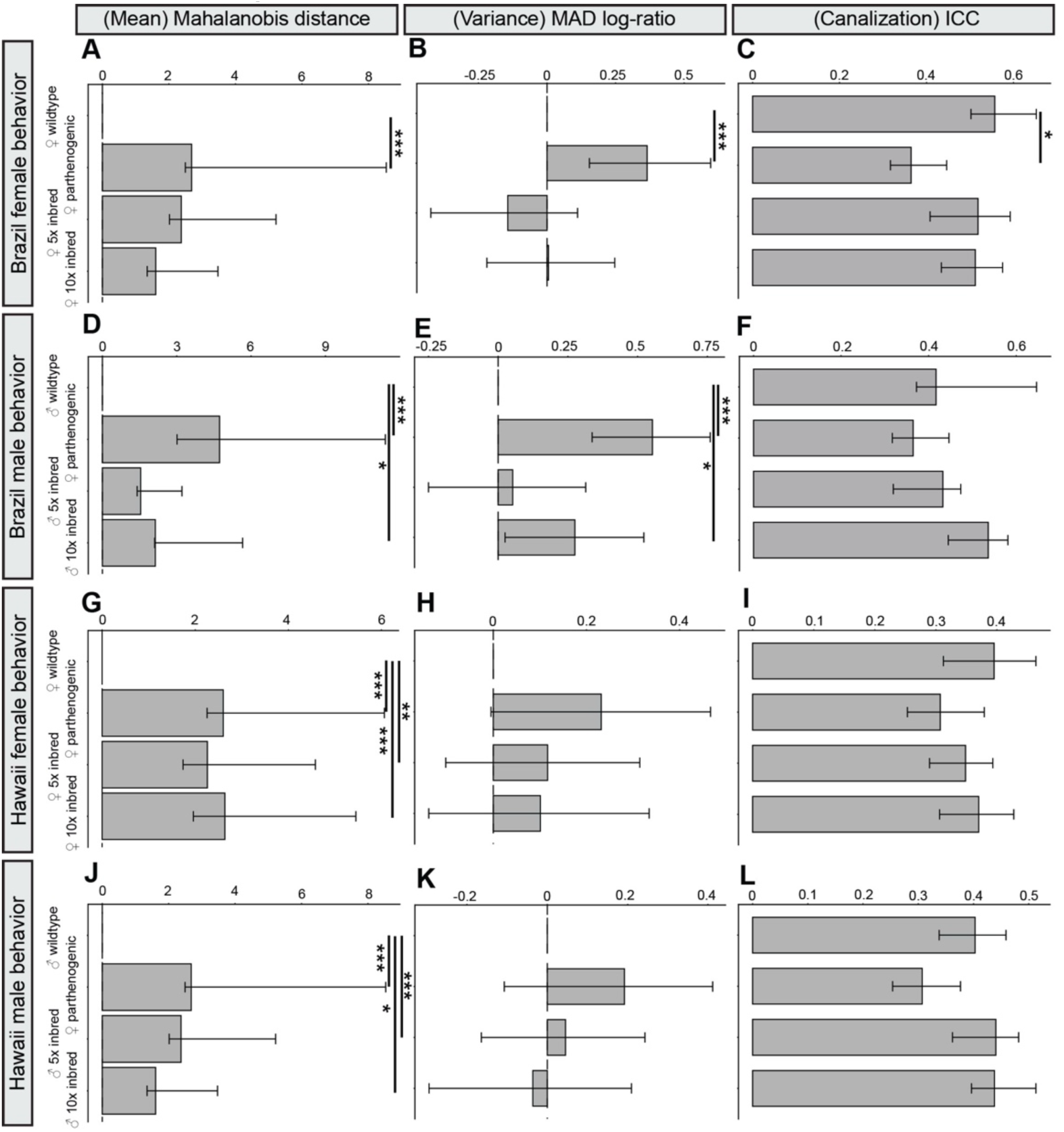
Reanalysis of Buridan inbreeding data confirms altered behavioral means in inbred and parthenogenic flies, with mixed effects on overall interindividual variation and behavioral consistency. Buridan data from Fig. 4 and Supplementary Figs. S22 and S24 were reanalyzed using Mahalanobis distance, MAD log-ratio, and the intraclass correlation coefficient (ICC). **(A-C)** Brazil females. **(A)** Mahalanobis distance indicates a strong multivariate change in behavioral means between female wild-type and parthenogenic flies, whereas neither the 5× nor the 10× inbred line differs significantly from wild-type controls. Pairwise permutation tests with Benjamini-Hochberg correction: female WT versus parthenogenic, P < 0.001; female WT versus female 5× inbred, P = 0.31; female WT versus female 10× inbred, P = 0.3. **(B)** MAD log-ratio indicates increased overall interindividual behavioral variation in parthenogenic females, but not in the 5× or 10× inbred lines. Pairwise permutation tests with Benjamini-Hochberg correction: female WT versus parthenogenic, P < 0.001; female WT versus female 5× inbred, P = 0.35; female WT versus female 10× inbred, P = 0.96. **(C)** ICC analysis of behavioral consistency in Brazil-derived females. The current source text contains placeholder values and should be completed before submission: female WT versus parthenogenic, P = 0.012; female WT versus female 5× inbred, P = 0.6; female WT versus female 10× inbred, P = 0.3. **(D-F)** Brazil males. **(D)** Mahalanobis distance indicates a strong multivariate change in behavioral means between male wild-type and parthenogenic flies, no detectable change in the 5× inbred line, and a weaker but significant change in the 10× inbred line. Pairwise permutation tests with Benjamini-Hochberg correction: male WT versus parthenogenic, P < 0.001; male WT versus male 5× inbred, P = 0.22; male WT versus male 10× inbred, P = 0.012. **(E)** MAD log-ratio indicates increased overall interindividual behavioral variation in parthenogenic males and in the 10× inbred line, but not in the 5× inbred line. Pairwise permutation tests with Benjamini-Hochberg correction: male WT versus parthenogenic, P < 0.001; male WT versus male 5× inbred, P = 0.67; male WT versus male 10× inbred, P = 0.032. **(F)** ICC analysis of behavioral consistency in Brazil-derived males. The current source text contains placeholder values and should be completed before submission: male WT versus parthenogenic, P = 0.49; male WT versus male 5× inbred, P = 0.82; male WT versus male 10× inbred, P = 0.13. **(G-I)** Hawaii females. **(G)** Mahalanobis distance indicates strong multivariate changes in behavioral means between female wild-type and all three comparison groups. Pairwise permutation tests with Benjamini-Hochberg correction: female WT versus parthenogenic, P < 0.001; female WT versus female 5× inbred, P = 0.002; female WT versus female 10× inbred, P < 0.001. **(H)** MAD log-ratio indicates no significant difference in overall interindividual behavioral variation between female wild-type and Hawaiian parthenogenic or inbred lines. Pairwise permutation tests with Benjamini-Hochberg correction: female WT versus parthenogenic, P = 0.14; female WT versus female 5× inbred, P = 0.34; female WT versus female 10× inbred, P = 0.36. **(I)** ICC analysis shows no significant difference in behavioral consistency between female wild-type and Hawaiian parthenogenic or inbred lines. Pairwise permutation tests with Benjamini-Hochberg correction: female WT versus parthenogenic, P = 0.43; female WT versus female 5× inbred, P = 0.57; female WT versus female 10× inbred, P = 0.65. **(J-L)** Hawaii males. **(J)** Mahalanobis distance indicates strong multivariate changes in behavioral means between male wild-type and Hawaiian parthenogenic and 5× inbred flies, and a weaker but significant change in the 10× inbred line. Pairwise permutation tests with Benjamini-Hochberg correction: male WT versus parthenogenic, P < 0.001; male WT versus male 5× inbred, P < 0.001; male WT versus male 10× inbred, P = 0.038. **(K)** MAD log-ratio indicates no significant difference in overall interindividual behavioral variation between male wild-type and Hawaiian parthenogenic or inbred lines. Pairwise permutation tests with Benjamini-Hochberg correction: male WT versus parthenogenic, P = 0.12; male WT versus male 5× inbred, P = 0.7; male WT versus male 10× inbred, P = 0.7. **(L)** ICC analysis shows no significant difference in behavioral consistency between male wild-type and Hawaiian parthenogenic or inbred lines. Pairwise permutation tests with Benjamini-Hochberg correction: male WT versus parthenogenic, P = 0.098; male WT versus male 5× inbred, P = 0.49; male WT versus male 10× inbred, P = 0.49. Asterisks denote statistical significance: p < 0.05 (*), p < 0.01 (**), p < 0.001 (***).

**Supplementary Fig. S24:**
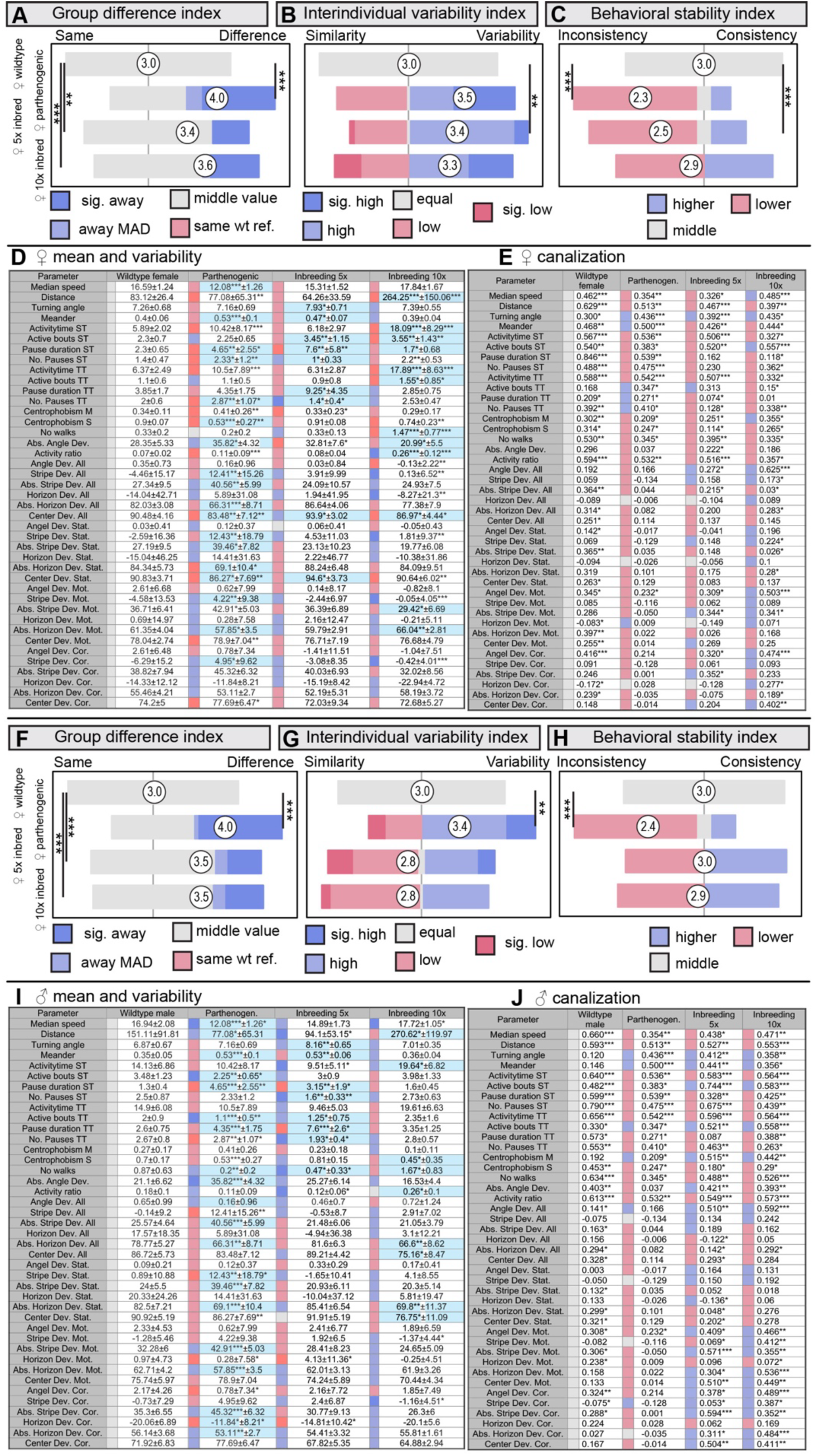
Inbreeding in Hawaiian *Drosophila mercatorum* alters mean behavior and behavioral canalization, with limited effects on overall interindividual variation. Buridan’s analysis shows that 5 and 10 generations of inbreeding affect behavioral means and day-to-day behavioral consistency, whereas effects on overall interindividual variation are more selective. **(A) Mean-difference index** across 41 behavioral parameters shows that the 5× and 10× inbred female lines are intermediate between wild-type and parthenogenic females. Full parameter-level data are shown in (D). Pairwise Wilcoxon tests with Benjamini-Hochberg correction: female WT versus parthenogenic, P = 6.5 × 10⁻⁸; female WT versus 5× inbred, P = 0.002; female WT versus 10× inbred, P = 4.7 × 10⁻⁵. **(B) Variability index** based on the median absolute deviation (MAD) across the 41 behavioral parameters indicates increased interindividual variation in the 5× inbred line. In contrast, parthenogenic and 10× inbred females do not differ significantly from wild-type females. Full parameter-level data are shown in (D). Pairwise Wilcoxon tests with Benjamini-Hochberg correction: female WT versus parthenogenic, P = 0.1; female WT versus 5× inbred, P = 0.004; female WT versus 10× inbred, P = 0.2. **(C) Behavioral consistency (canalization) index** across the 41 tested behavioral parameters shows reduced day-to-day behavioral consistency in parthenogenic females and in the 5× inbred line. In contrast, the 10× inbred line does not differ significantly from wild-type females. Full parameter-level data are shown in (E). Pairwise Wilcoxon tests with Benjamini-Hochberg correction: female WT versus parthenogenic, P = 5.7 × 10⁻⁹; female WT versus 5× inbred, P = 5.1 × 10⁻⁵; female WT versus 10× inbred, P = 0.3. **(D)** Parameter-level analysis of mean and dispersion across all 41 Buridan parameters for female wild-type, parthenogenic, 5× inbred, and 10× inbred flies. The table shows group medians, with significance assessed by pairwise Wilcoxon tests with Benjamini-Hochberg correction, and MAD values, with significance assessed by Levene’s test. Colors denote relative MAD values, from low (blue) to high (red), and darker colors indicate parameters that differ significantly from both comparison groups. **(E)** Parameter-level analysis of day-to-day behavioral consistency across 3 testing days for all 41 Buridan parameters in female wild-type, parthenogenic, 5× inbred, and 10× inbred flies. The table shows Pearson correlation coefficients for all pairwise day comparisons after Fisher’s z-transformation. Asterisks denote significant correlations across the three-day pair comparisons. Colors denote correlation strength, from low (red) to high (blue), with intermediate values in grey. Sample sizes: female WT (N = 33), parthenogenic (N = 31), female 5x inbred (N = 24), female 10x inbred (N = 30). **(F) Mean-difference index** across 41 behavioral parameters shows that the 5× and 10× inbred male lines are intermediate between wild-type and parthenogenic flies. Full parameter-level data are shown in (I). Pairwise Wilcoxon tests with Benjamini-Hochberg correction: male WT versus parthenogenic, P = 1.4 × 10⁻⁷; male WT versus 5× inbred, P = 0.0004; male WT versus 10× inbred, P = 0.0002. **(G) Variability index** based on MAD indicates increased interindividual behavioral variation in parthenogenic males, whereas the 5× and 10× inbred male lines do not differ significantly from wild-type males. Full parameter-level data are shown in (I). Pairwise Wilcoxon tests with Benjamini-Hochberg correction: male WT versus parthenogenic, P = 0.002; male WT versus 5× inbred, P = 0.3; male WT versus 10× inbred, P = 0.09. **(H)** The behavio**ral consistency (canalization) index** shows reduced day-to-day behavioral consistency in parthenogenic males, whereas the 5× and 10× inbred male lines do not differ significantly from wild-type males. Full parameter-level data are shown in (J). Pairwise Wilcoxon tests with Benjamini-Hochberg correction: male WT versus parthenogenic, P = 7.6 × 10⁻⁸; male WT versus 5× inbred, P = 0.8; male WT versus 10× inbred, P = 0.5. **(I)** Parameter-level analysis of mean and dispersion across all 41 Buridan parameters for male wild-type, parthenogenic, 5× inbred, and 10× inbred flies. The table shows group medians, with significance assessed by pairwise Wilcoxon tests with Benjamini-Hochberg correction, and MAD values, with significance assessed by Levene’s test. Colors denote relative MAD values, from low (blue) to high (red), and darker colors indicate parameters that differ significantly from both comparison groups. **(J)** Parameter-level analysis of day-to-day behavioral consistency across 3 testing days for all 41 Buridan parameters in male wild-type, parthenogenic, 5× inbred, and 10× inbred flies. The table shows Pearson correlation coefficients for all pairwise day comparisons after Fisher’s z-transformation. Asterisks denote significant correlations across the three-day pair comparisons. Colors denote correlation strength, from low (red) to high (blue), with intermediate values in grey. Sample sizes: male WT (N = 31), parthenogenic (N = 31), male 5x inbred (N = 23), male 10x inbred (N = 30). Asterisks denote statistical significance: p < 0.05 (*), p < 0.01 (**), p < 0.001 (***). Only in (E) and (J) do the asterisks indicate the number of significant comparisons between days.

**Supplementary Fig. S25:**
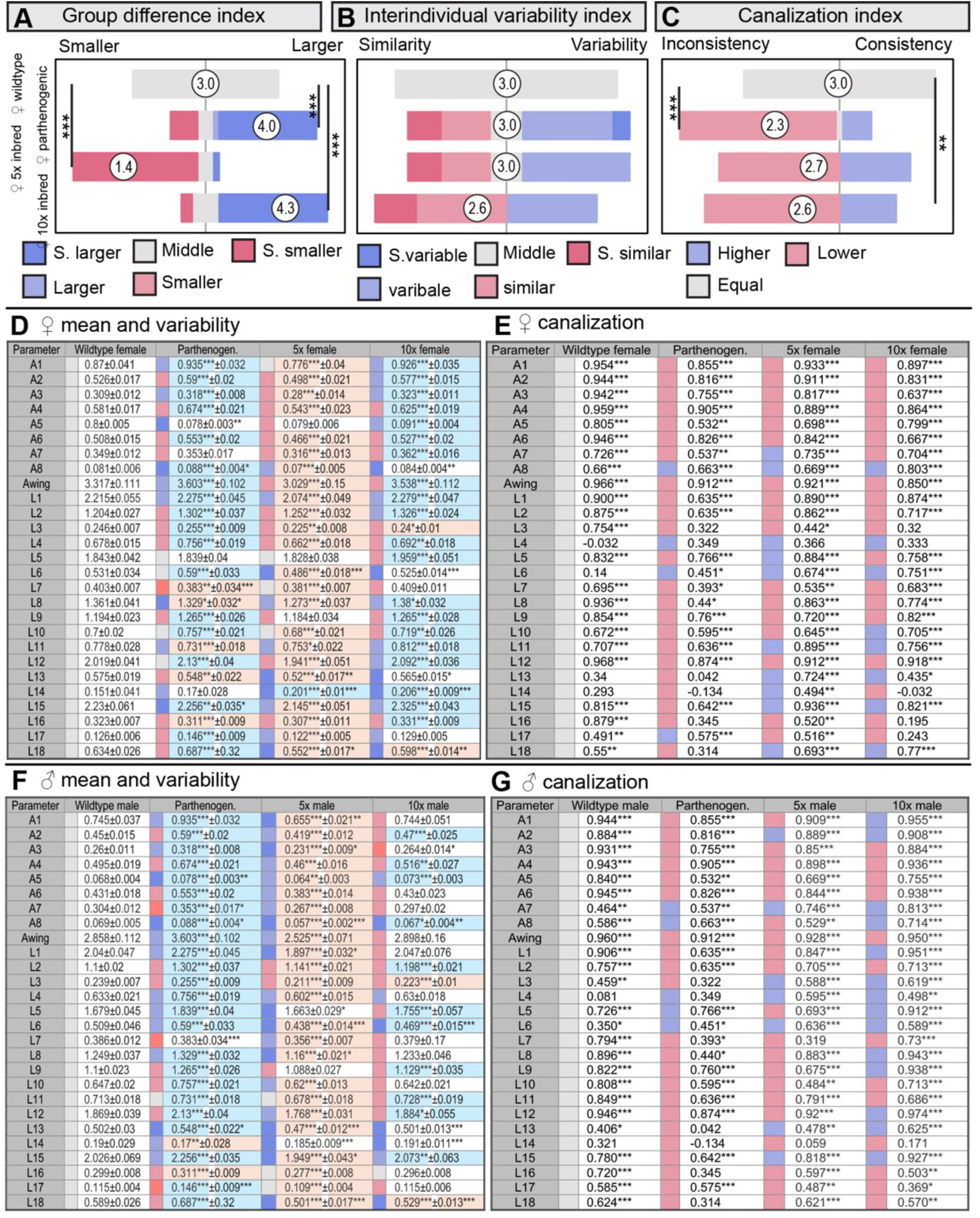
Inbreeding in Brazilian *Drosophila mercatorum* alters wing morphology and developmental stability, with limited effects on overall interindividual variation. Wing morphometry shows that inbreeding changes mean wing anatomy and affects developmental canalization, whereas effects on overall interindividual variation are comparatively modest. **(A) Mean-difference index** across 27 wing parameters shows that the 5×, 10×, and parthenogenic female lines all differ from wild-type controls. Mild inbreeding for 5 generations changes wing morphology towards smaller values, whereas 10 generations changes wing morphology towards the enlarged phenotype observed in parthenogenic flies. Full parameter-level data are shown in (D). Pairwise Wilcoxon tests with Benjamini-Hochberg correction: female WT versus parthenogenic, P = 2.0 × 10⁻⁴; female WT versus 5× inbred, P = 3.3 × 10⁻⁹; female WT versus 10× inbred, P = 1.1 × 10⁻⁶. **(B) Variability index** based on the median absolute deviation (MAD) across the 27 wing parameters indicates no significant difference in overall interindividual wing variation among female groups. Full parameter-level data are shown in (D). Pairwise Wilcoxon tests with Benjamini-Hochberg correction: female WT versus parthenogenic, P = 0.44; female WT versus 5× inbred, P = 0.44; female WT versus 10× inbred, P = 0.21. **(C) Canalization index** based on left-right correspondence across the 27 wing parameters shows reduced developmental stability in parthenogenic females and in the 10× inbred line. In contrast, the 5× inbred line does not differ significantly from wild-type females. Full parameter-level data are shown in (E). Pairwise Wilcoxon tests with Benjamini-Hochberg correction: female WT versus parthenogenic, P = 2.5 × 10⁻⁶; female WT versus 5× inbred, P = 0.08; female WT versus 10× inbred, P = 0.005. **(D)** Parameter-level analysis of mean and dispersion across all 27 wing parameters for female wild-type, parthenogenic, 5× inbred, and 10× inbred flies. The table shows group medians, with significance assessed by pairwise Wilcoxon tests with Benjamini-Hochberg correction, and MAD values, with significance assessed by Levene’s test. Colors denote relative MAD values, from low (blue) to high (red), and darker colors indicate parameters that differ significantly from both comparison groups. **(E)** Parameter-level analysis of right-left correspondence across the 27 wing parameters for female wild-type, parthenogenic, 5× inbred, and 10× inbred flies. The table shows Pearson correlation coefficients for bilateral wing correspondence, with asterisks denoting statistical significance. Colors denote correlation strength, from low (red) to high (blue), with intermediate values in grey. Sample sizes: female WT, n = 29; parthenogenic, n = 31; female 5× inbred, n = 27; female 10× inbred, n = 31. **(F)** Parameter-level analysis of mean and dispersion across all 27 wing parameters for male wild-type, parthenogenic, 5× inbred, and 10× inbred flies. The table shows group medians, with significance assessed by pairwise Wilcoxon tests with Benjamini-Hochberg correction, and MAD values, with significance assessed by Levene’s test. Colors denote relative MAD values, from low (blue) to high (red), and darker colors indicate parameters that differ significantly from both comparison groups. **(G)** Parameter-level analysis of right-left correspondence across the 27 wing parameters for male wild-type, parthenogenic, 5× inbred, and 10× inbred flies. The table shows Pearson correlation coefficients for bilateral wing correspondence, with asterisks denoting statistical significance. Colors denote correlation strength, from low (red) to high (blue), with intermediate values in grey. Sample sizes: male WT (N = 33), parthenogenic (N = 31), male 5x inbred (N = 29), male 10x inbred (N = 29). Asterisks denote statistical significance: p < 0.05 (*), p < 0.01 (**), p < 0.001 (***). Only in (E) and (G) do asterisks indicate the number of significant comparisons between different days.

**Supplementary Fig. S26:**
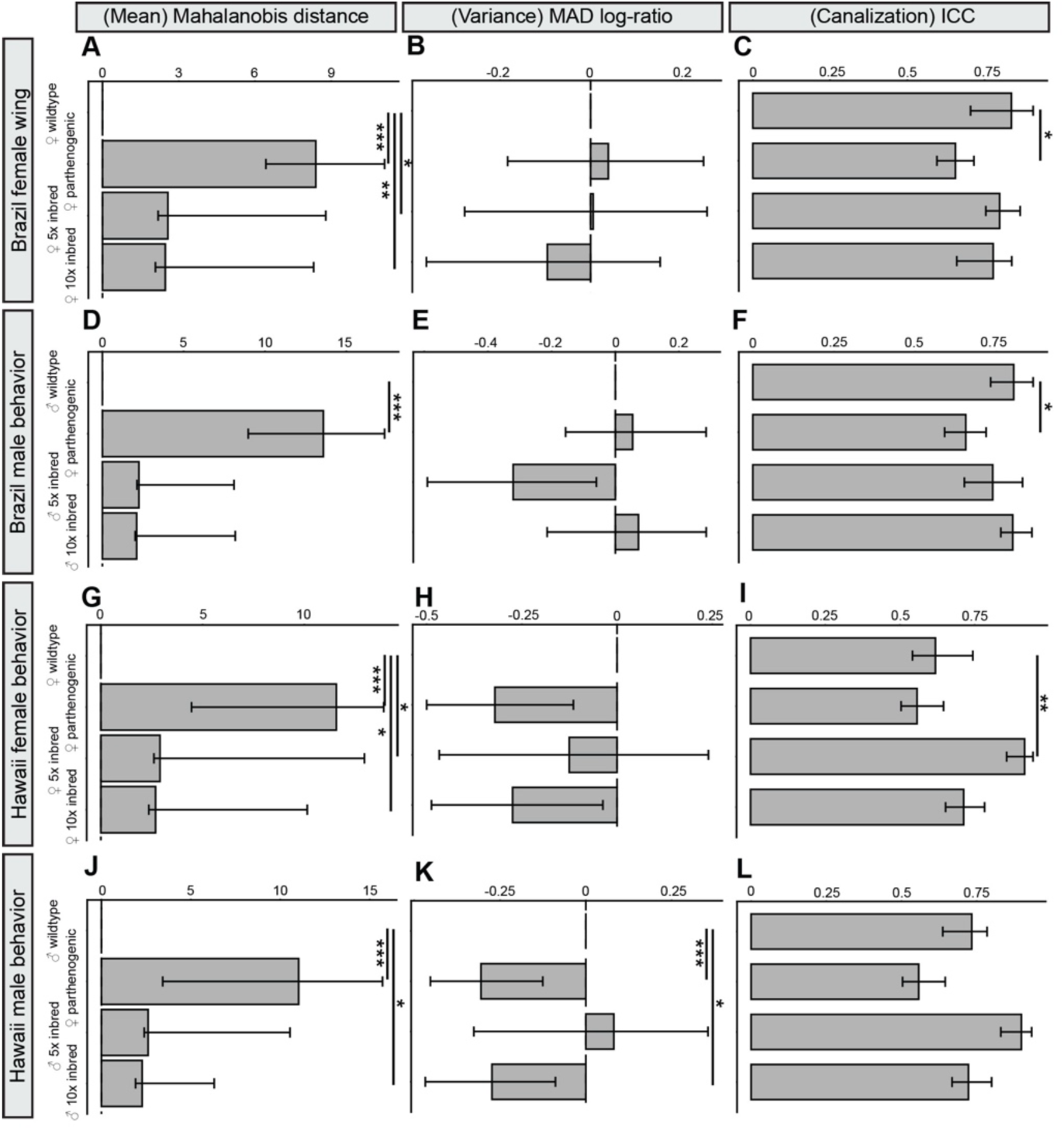
Reanalysis of wing inbreeding data confirms altered mean morphology and selective effects on developmental stability, with limited effects on overall interindividual variation. Wing morphometry data from Fig. 4 and Supplementary Figs. S25 and S27 were reanalyzed using Mahalanobis distance, MAD log-ratio, and the intraclass correlation coefficient (ICC). **(A-C)** Brazil females. **(A)** Mahalanobis distance indicates significant multivariate changes in wing morphology between female wild-type and parthenogenic flies, and between 5× and 10× inbred flies. Pairwise permutation tests with Benjamini-Hochberg correction: female WT versus parthenogenic, P < 0.001; female WT versus female 5× inbred, P = 0.01; female WT versus female 10× inbred, P = 0.009. **(B)** MAD log-ratio indicates no significant difference in overall interindividual wing variation between female wild-type and parthenogenic or inbred lines. Pairwise permutation tests with Benjamini-Hochberg correction: female WT versus parthenogenic, P = 0.88; female WT versus female 5× inbred, P = 0.96; female WT versus female 10× inbred, P = 0.71. **(C)** ICC analysis indicates reduced left-right correspondence in parthenogenic females, whereas the 5× and 10× inbred lines do not differ significantly from female wild-type controls. Pairwise permutation tests with Benjamini-Hochberg correction: female WT versus parthenogenic, P = 0.012; female WT versus female 5× inbred, P = 0.57; female WT versus female 10× inbred, P = 0.4. **(D-F)** Brazil males. **(D)** Mahalanobis distance indicates a strong multivariate change in wing morphology between male wild-type and parthenogenic flies. In contrast, neither the 5× nor the 10× inbred line differs significantly from male wild-type controls. Pairwise permutation tests with Benjamini-Hochberg correction: male WT versus parthenogenic, P < 0.001; male WT versus male 5× inbred, P = 0.07; male WT versus male 10× inbred, P = 0.09. **(E)** MAD log-ratio indicates no significant difference in overall interindividual wing variation between male wild-type and parthenogenic or inbred lines. Pairwise permutation tests with Benjamini-Hochberg correction: male WT versus parthenogenic, P = 0.91; male WT versus male 5× inbred, P = 0.12; male WT versus male 10× inbred, P = 0.91. **(F)** ICC analysis indicates reduced left-right correspondence in parthenogenic males, whereas the 5× and 10× inbred male lines do not differ significantly from wild-type controls. Pairwise permutation tests with Benjamini-Hochberg correction: male WT versus parthenogenic, P = 0.024; male WT versus male 5× inbred, P = 0.27; male WT versus male 10× inbred, P = 0.95. **(G-I)** Hawaii females. **(G)** Mahalanobis distance indicates significant multivariate changes in wing morphology between female wild-type and parthenogenic flies, and between 5× and 10× inbred flies. Pairwise permutation tests with Benjamini-Hochberg correction: female WT versus parthenogenic, P < 0.001; female WT versus female 5× inbred, P = 0.023; female WT versus female 10× inbred, P = 0.015. **(H)** MAD log-ratio indicates no significant difference in overall interindividual wing variation between female wild-type and Hawaiian parthenogenic or inbred lines. Pairwise permutation tests with Benjamini-Hochberg correction: female WT versus parthenogenic, P = 0.3; female WT versus female 5× inbred, P = 0.59; female WT versus female 10× inbred, P = 0.3. **(I)** ICC analysis indicates reduced left-right correspondence in the Hawaiian 5× inbred female line, whereas parthenogenic and 10× inbred females do not differ significantly from wild-type controls. Pairwise permutation tests with Benjamini-Hochberg correction: female WT versus parthenogenic, P = 0.66; female WT versus female 5× inbred, P = 0.006; female WT versus female 10× inbred, P = 0.49. **(J-L)** Hawaii males. **(J)** Mahalanobis distance indicates a strong multivariate change in wing morphology between male wild-type and parthenogenic flies, with a weaker but significant change in the 10× inbred line and no significant change in the 5× inbred line after correction. Pairwise permutation tests with Benjamini-Hochberg correction: male WT versus parthenogenic, P < 0.001; male WT versus male 5× inbred, P = 0.053; male WT versus male 10× inbred, P = 0.04. **(K)** MAD log-ratio indicates no significant difference in overall interindividual wing variation between male wild-type and Hawaiian parthenogenic or inbred lines. Pairwise permutation tests with Benjamini-Hochberg correction: male WT versus parthenogenic, P = 0.12; male WT versus male 5× inbred, P = 0.81; male WT versus male 10× inbred, P = 0.16. **(L)** ICC analysis indicates no significant difference in left-right correspondence between male wild-type and Hawaiian parthenogenic or inbred lines. Pairwise permutation tests with Benjamini-Hochberg correction: male WT versus parthenogenic, P = 0.22; male WT versus male 5× inbred, P = 0.15; male WT versus male 10× inbred, P = 0.91. Asterisks denote statistical significance: p < 0.05 (*), p < 0.01 (**), p < 0.001 (***).

**Supplementary Fig. S27:**
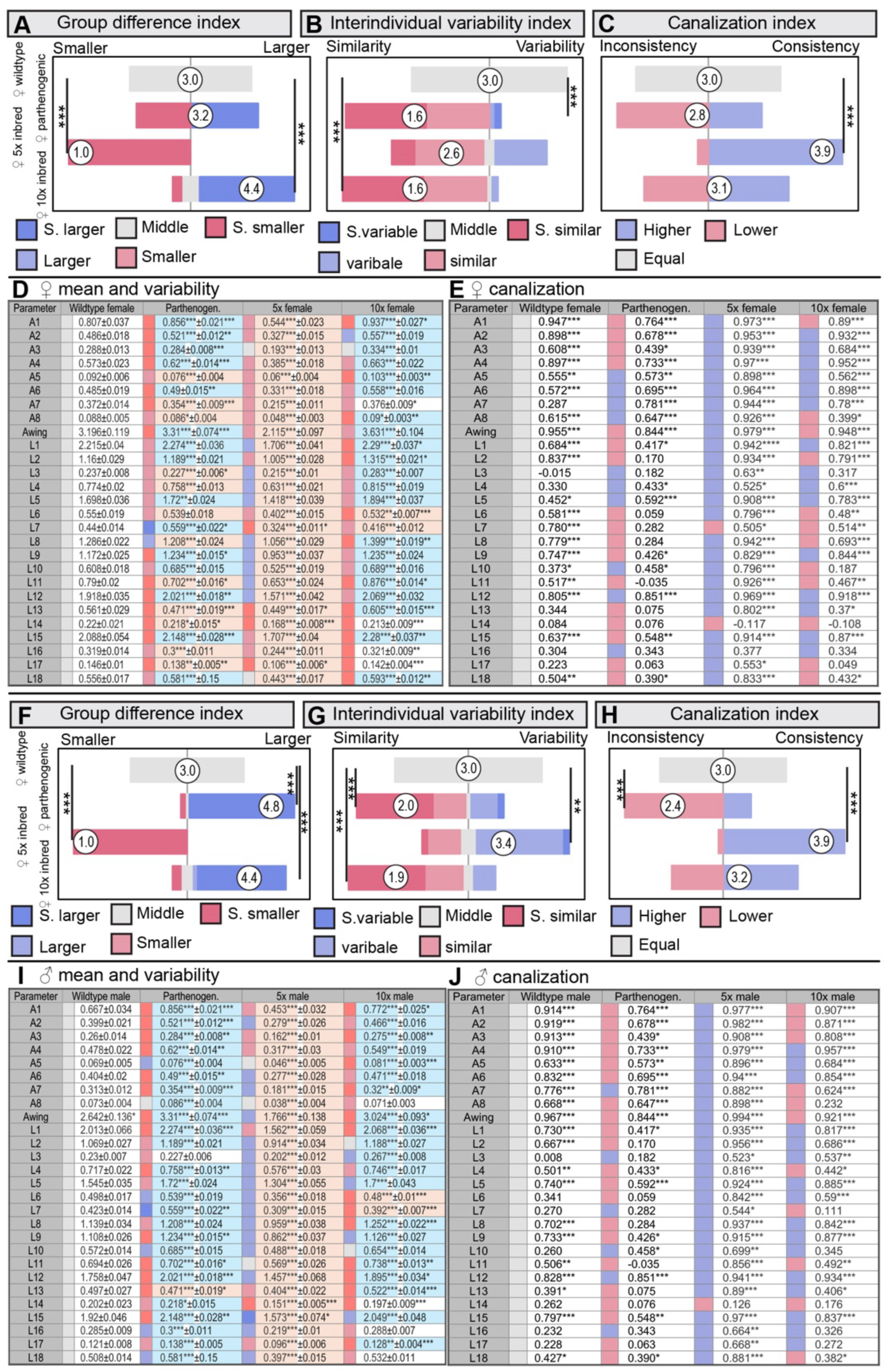
Inbreeding in Hawaiian *Drosophila mercatorum* alters wing morphology, interindividual variation, and developmental stability. Wing morphometry shows that inbreeding in the Hawaiian strain affects mean wing anatomy, interindividual variation, and developmental canalization, with sex- and genotype-specific effects. **(A) Mean-difference index** across 27 wing parameters shows that the 5× and 10× inbred female lines differ significantly from wild-type controls, whereas parthenogenic females do not. Mild inbreeding for 5 generations changes wing morphology towards smaller values, whereas 10 generations changes wing morphology towards larger values. Full parameter-level data are shown in (D). Pairwise Wilcoxon tests with Benjamini-Hochberg correction: female WT versus parthenogenic, P = 0.45; female WT versus 5× inbred, P = 3.6 × 10⁻¹³; female WT versus 10× inbred, P = 3.3 × 10⁻⁷. **(B) Variability index** based on the median absolute deviation (MAD) across the 27 wing parameters indicates reduced interindividual variation in parthenogenic and 10× inbred females, whereas the 5× inbred female line does not differ significantly from wild type. Full parameter-level data are shown in (D). Pairwise Wilcoxon tests with Benjamini-Hochberg correction: female WT versus parthenogenic, P = 5.9 × 10⁻⁹; female WT versus 5× inbred, P = 0.07; female WT versus 10× inbred, P = 9.1 × 10⁻¹⁰. **(C) Canalization index** based on left-right correspondence across the 27 wing parameters indicates increased developmental stability in the 5× inbred female line. In contrast, parthenogenic and 10× inbred females do not differ significantly from wild-type controls. Full parameter-level data are shown in (E). Pairwise Wilcoxon tests with Benjamini-Hochberg correction: female WT versus parthenogenic, P = 0.08; female WT versus 5× inbred, P = 1.1 × 10⁻⁹; female WT versus 10× inbred, P = 0.45. **(D)** Parameter-level analysis of mean and dispersion across all 27 wing parameters for female wild-type, parthenogenic, 5× inbred, and 10× inbred flies. The table shows group medians, with significance assessed by pairwise Wilcoxon tests with Benjamini-Hochberg correction, and MAD values, with significance assessed by Levene’s test. Colors denote relative MAD values, from low (blue) to high (red), and darker colors indicate parameters that differ significantly from both comparison groups. **(E)** Parameter-level analysis of right-left correspondence across the 27 wing parameters for female wild-type, parthenogenic, 5× inbred, and 10× inbred flies. The table shows Pearson correlation coefficients for bilateral wing correspondence, with asterisks denoting statistical significance. Colors denote correlation strength, from low (red) to high (blue), with intermediate values in grey. Sample sizes: female WT (N = 30), parthenogenic (N = 30), female 5x inbred (N = 20), female 10x inbred (N = 32). **(F) Mean-difference index** across 27 wing parameters shows that parthenogenic, 5× inbred, and 10× inbred male lines all differ significantly from wild-type controls. Mild inbreeding for 5 generations changes wing morphology towards smaller values, whereas 10 generations changes wing morphology towards larger values, similar to those of parthenogenic flies. Full parameter-level data are shown in (I). Pairwise Wilcoxon tests with Benjamini-Hochberg correction: male WT versus parthenogenic, P = 1.4 × 10⁻¹⁰; male WT versus 5× inbred, P = 3.6 × 10⁻¹³; male WT versus 10× inbred, P = 1.1 × 10⁻⁷. **(G) Variability index** based on MAD indicates reduced interindividual variation in parthenogenic and 10× inbred males, but increased variation in the 5× inbred male line. Full parameter-level data are shown in (I). Pairwise Wilcoxon tests with Benjamini-Hochberg correction: male WT versus parthenogenic, P = 0.0004; male WT versus 5× inbred, P = 0.009; male WT versus 10× inbred, P = 1.3 × 10⁻⁵. **(H) Canalization index** indicates increased developmental stability in the 5× inbred male line and reduced developmental stability in parthenogenic males. In contrast, the 10× inbred male line does not differ significantly from wild-type males. Full parameter-level data are shown in (J). Pairwise Wilcoxon tests with Benjamini-Hochberg correction: male WT versus parthenogenic, P = 0.0001; male WT versus 5× inbred, P = 2.5 × 10⁻¹¹; male WT versus 10× inbred, P = 0.21. **(I)** Parameter-level analysis of mean and dispersion across all 27 wing parameters for male wild-type, parthenogenic, 5× inbred, and 10× inbred flies. The table shows group medians, with significance assessed by pairwise Wilcoxon tests with Benjamini-Hochberg correction, and MAD values, with significance assessed by Levene’s test. Colors denote relative MAD values, from low (blue) to high (red), and darker colors indicate parameters that differ significantly from both comparison groups. **(J)** Parameter-level analysis of right-left correspondence across the 27 wing parameters for male wild-type, parthenogenic, 5× inbred, and 10× inbred flies. The table shows Pearson correlation coefficients for bilateral wing correspondence, with asterisks denoting statistical significance. Colors denote correlation strength, from low (red) to high (blue), with intermediate values in grey. Sample sizes: male WT (N = 30), parthenogenic (N = 30), male 5x inbred (N = 16), male 10x inbred (N = 32). Asterisks denote statistical significance: p < 0.05 (*), p < 0.01 (**), p < 0.001 (***). Only in (E) and (J) do asterisks refer to the number of significant comparisons between different days.

**Supplementary Table 1:** Genotypes used in each figure. Strain identity, NDSSC stock number, geographic origin, and experimental role of each *Drosophila mercatorum* genotype used in the main and supplementary figures, including wild-type, parthenogenic, facultative parthenogenic, F_1_ rescue, and inbred lines.

| Figure | Genotypes |
| --- | --- |
| Fig. 1 | Brazil wildtype (NDSSC15082-1521.25) and Brazil parthenogenic (NDSSC15082-1521.04). |
| Fig. 2 | Brazil wildtype (NDSSC15082-1521.25) and Brazil parthenogenic (NDSSC15082-1521.04). |
| Fig. 3 | Brazil wildtype (NDSSC15082-1521.25) and Brazil parthenogenic (NDSSC15082-1521.04). |
| Fig. 4 | Brazil wildtype (NDSSC15082-1521.25), Brazil parthenogenic (NDSSC15082-1521.04), 5x inbred Brazil wildtype (NDSSC15082-1521.25) and 10x inbred Brazil wildtype (NDSSC15082-1521.25). |
| Fig. S1 | (A and E) Brazil wildtype (NDSSC15082-1521.25) and Brazil parthenogenic (NDSSC15082-1521.04). (B-D and F) Hawaii wildtype (NDSSC 15082-1521.22), and Hawaii parthenogenic (NDSSC 15082-1525.05). |
| Fig. S2 | (A-C) Brazil wildtype (NDSSC15082-1521.25) and Brazil parthenogenic (NDSSC15082-1521.04). (D-F) Hawaii wildtype (NDSSC 15082-1521.22), and Hawaii parthenogenic (NDSSC 15082-1525.05). |
| Fig. S3 | Brazil wildtype (NDSSC15082-1521.25) and Brazil parthenogenic (NDSSC15082-1521.04). |
| Fig. S4 | Brazil wildtype (NDSSC15082-1521.25) and Brazil parthenogenic (NDSSC15082-1521.04). |
| Fig. S5 | (A and C) Brazil wildtype (NDSSC15082-1521.25) and Brazil parthenogenic (NDSSC15082-1521.04). (B and D) Hawaii wildtype (NDSSC 15082-1521.22), and Hawaii parthenogenic (NDSSC 15082-1525.05). |
| Fig. S6 | Brazil wildtype (NDSSC15082-1521.25) and Brazil parthenogenic (NDSSC15082-1521.04). |
| Fig. S7 | Hawaii wildtype (NDSSC 15082-1521.22) and Hawaii parthenogenic (NDSSC 15082-1525.05). |
| Fig. S8 | (A-C) Brazil wildtype (NDSSC15082-1521.25) and Brazil parthenogenic (NDSSC15082-1521.04). (D-H) Hawaii wildtype (NDSSC 15082-1521.22) and Hawaii parthenogenic (NDSSC 15082-1525.05). |
| Fig. S9 | (A-C) Brazil wildtype (NDSSC15082-1521.25) and Brazil parthenogenic (NDSSC15082-1521.04). (D-F) Hawaii wildtype (NDSSC 15082-1521.22), and Hawaii parthenogenic (NDSSC 15082-1525.05). |
| Fig. S10 | (A) Brazil wildtype (NDSSC15082-1521.25). (B-F) Brazil wildtype (NDSSC15082-1521.25) and Brazil parthenogenic (NDSSC15082-1521.04). |
| Fig. S11 | Brazil wildtype (NDSSC15082-1521.25) and Brazil parthenogenic (NDSSC15082-1521.04). |
| Fig. S12 | Brazil wildtype (NDSSC15082-1521.25) and Brazil parthenogenic (NDSSC15082-1521.04). |
| Fig. S13 | Brazil wildtype (NDSSC15082-1521.25) and Brazil parthenogenic (NDSSC15082-1521.04). |
| Fig. S14 | Brazil wildtype (NDSSC15082-1521.25) and Brazil parthenogenic (NDSSC15082-1521.04). |
| Fig. S15 | (A-I) Brazil wildtype (NDSSC15082-1521.25), Brazil parthenogenic (NDSSC15082-1521.04) and Brazil parthenogenic (NDSSC15082-1521.04) from a single mother. |
| Fig. S16 | (A-H) Brazil wildtype (NDSSC15082-1521.25), Brazil parthenogenic (NDSSC15082-1521.04) and Brazil parthenogenic (NDSSC15082-1521.04) from a single mother. |
| Fig. S17 | Brazil wildtype (NDSSC15082-1521.25), Brazil parthenogenic (NDSSC15082-1521.04), and a facultative parthenogenic strain (NDSSC 15082-1527.03). |
| Fig. S18 | Brazil wildtype (NDSSC15082-1521.25), Brazil parthenogenic (NDSSC15082-1521.04), and a facultative parthenogenic strain (NDSSC 15082-1527.03). |
| Fig. S19 | Brazil wildtype (NDSSC15082-1521.25), Brazil parthenogenic (NDSSC15082-1521.04) and an F1 of (NDSSC15082-1521.25) with (NDSSC15082-1521.04). |
| Fig. S20 | Brazil wildtype (NDSSC15082-1521.25), Brazil parthenogenic (NDSSC15082-1521.04) and an F1 of (NDSSC15082-1521.25) with (NDSSC15082-1521.04). |
| Fig. S21 | Brazil wildtype (NDSSC15082-1521.25), Brazil parthenogenic (NDSSC15082-1521.04) and an F1 of (NDSSC15082-1521.25) with (NDSSC15082-1521.04). |
| Fig. S22 | Brazil wildtype (NDSSC15082-1521.25), Brazil parthenogenic (NDSSC15082-1521.04), 5x and 10x inbred Brazil wildtype (NDSSC15082-1521.25). |
| Fig. S23 | (A-F) Brazil wildtype (NDSSC15082-1521.25), Brazil parthenogenic (NDSSC15082-1521.04), 5x and 10x inbred Brazil wildtype (NDSSC15082-1521.25). (G-L) Hawaii wildtype (NDSSC 15082-1521.22), Hawaii parthenogenic (NDSSC 15082-1525.05), and 5x and 10x inbred Hawaii wildtype (NDSSC 15082-1521.22). |
| Fig. S24 | Hawaii wildtype (NDSSC 15082-1521.22), Hawaii parthenogenic (NDSSC 15082-1525.05), and 5x and 10x inbred Hawaii wildtype (NDSSC 15082-1521.22). |
| Fig. S25 | Brazil wildtype (NDSSC15082-1521.25), Brazil parthenogenic (NDSSC15082-1521.04), 5x and 10x inbred Brazil wildtype (NDSSC15082-1521.25) |
| Fig. S26 | (A-F) Brazil wildtype (NDSSC15082-1521.25), Brazil parthenogenic (NDSSC15082-1521.04), 5x and 10x inbred Brazil wildtype (NDSSC15082-1521.25). (G-L) Hawaii wildtype (NDSSC 15082-1521.22), Hawaii parthenogenic (NDSSC 15082-1525.05), and 5x and 10x inbred Hawaii wildtype (NDSSC 15082-1521.22). |
| Fig. S27 | Hawaii wildtype (NDSSC 15082-1521.22), Hawaii parthenogenic (NDSSC 15082-1525.05), and 5x and 10x inbred Hawaii wildtype (NDSSC 15082-1521.22) |

**Supplementary Table 2:**
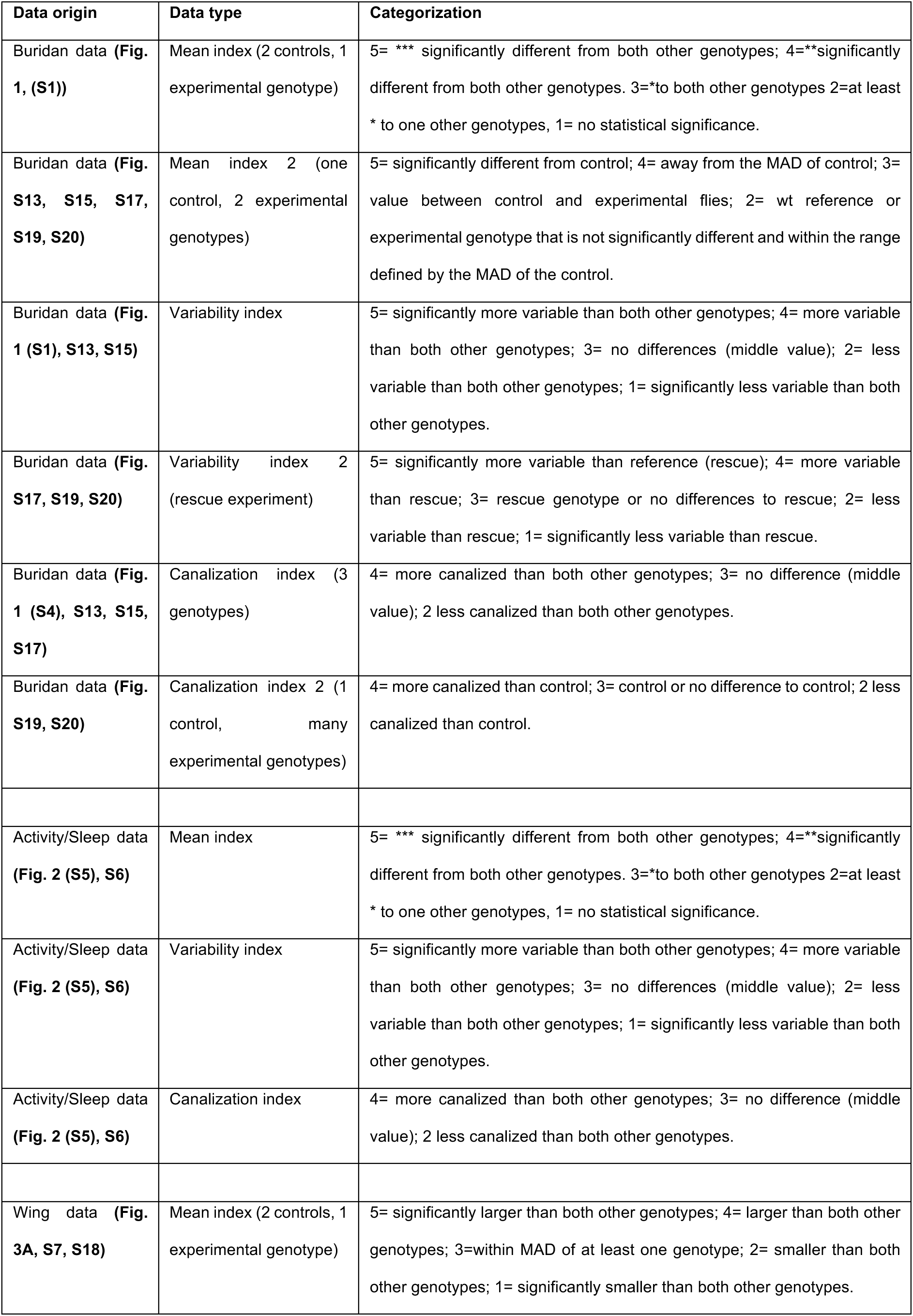
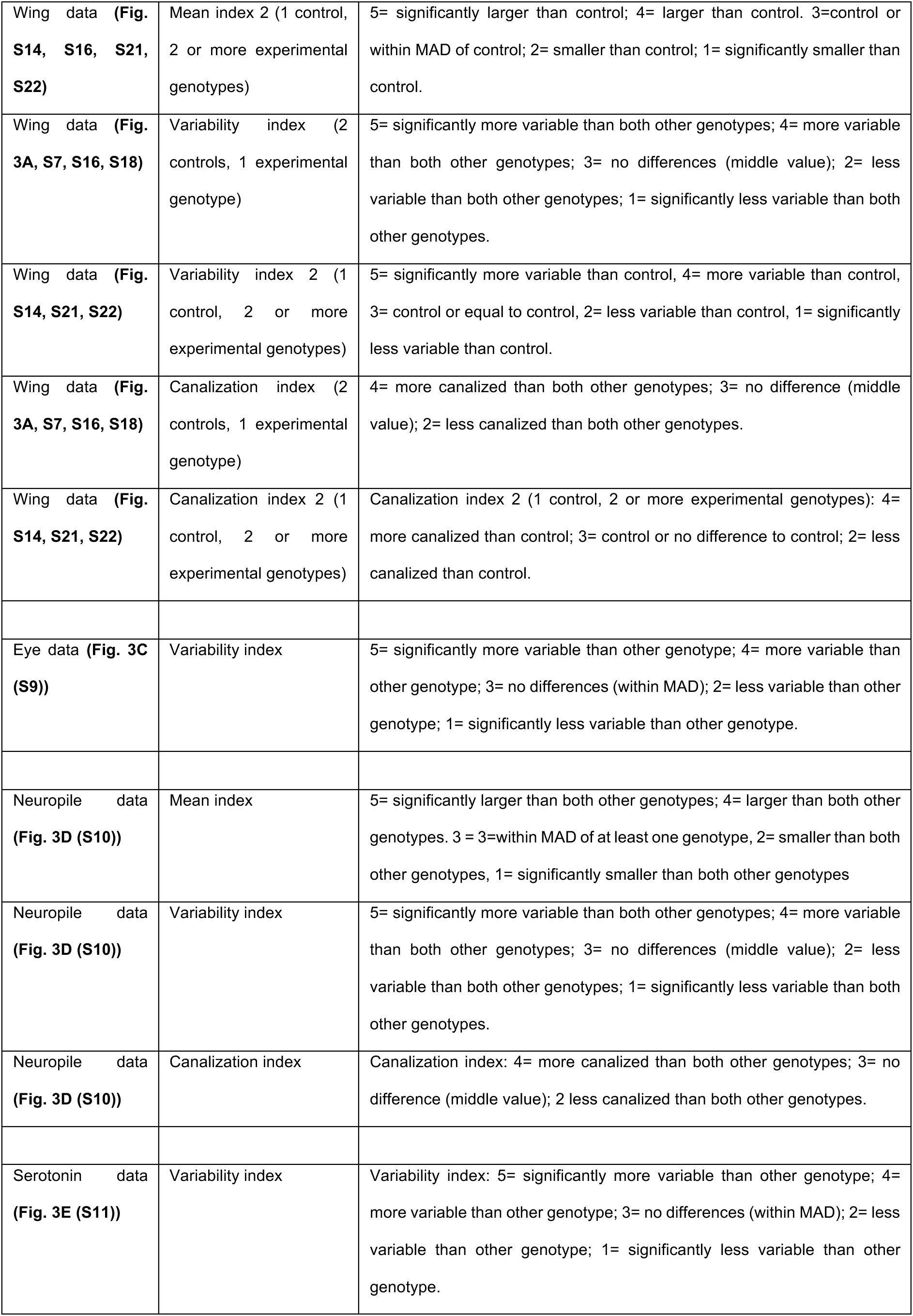
Calculation of index values.

| Data origin | Data type | Categorization |
| --- | --- | --- |
| Buridan data (**Fig. 1, (S1)**) | Mean index (2 controls, 1 experimental genotype) | 5= *** significantly different from both other genotypes; 4=**significantly different from both other genotypes. 3=*to both other genotypes 2=at least * to one other genotypes, 1= no statistical significance. |
| Buridan data (**Fig. S13, S15, S17, S19, S20**) | Mean index 2 (one control, 2 experimental genotypes) | 5= significantly different from control; 4= away from the MAD of control; 3= value between control and experimental flies; 2= wt reference or experimental genotype that is not significantly different and within the range defined by the MAD of the control. |
| Buridan data (**Fig. 1 (S1), S13, S15**) | Variability index | 5= significantly more variable than both other genotypes; 4= more variable than both other genotypes; 3= no differences (middle value); 2= less variable than both other genotypes; 1= significantly less variable than both other genotypes. |
| Buridan data (**Fig. S17, S19, S20**) | Variability index 2 (rescue experiment) | 5= significantly more variable than reference (rescue); 4= more variable than rescue; 3= rescue genotype or no differences to rescue; 2= less variable than rescue; 1= significantly less variable than rescue. |
| Buridan data (**Fig. 1 (S4), S13, S15, S17**) | Canalization index (3 genotypes) | 4= more canalized than both other genotypes; 3= no difference (middle value); 2 less canalized than both other genotypes. |
| Buridan data (**Fig. S19, S20**) | Canalization index 2 (1 control, many experimental genotypes) | 4= more canalized than control; 3= control or no difference to control; 2 less canalized than control. |
| Activity/Sleep data (**Fig. 2 (S5), S6**) | Mean index | 5= *** significantly different from both other genotypes; 4=**significantly different from both other genotypes. 3=*to both other genotypes 2=at least * to one other genotypes, 1= no statistical significance. |
| Activity/Sleep data (**Fig. 2 (S5), S6**) | Variability index | 5= significantly more variable than both other genotypes; 4= more variable than both other genotypes; 3= no differences (middle value); 2= less variable than both other genotypes; 1= significantly less variable than both other genotypes. |
| Activity/Sleep data (**Fig. 2 (S5), S6**) | Canalization index | 4= more canalized than both other genotypes; 3= no difference (middle value); 2 less canalized than both other genotypes. |
| Wing data (**Fig. 3A, S7, S18**) | Mean index (2 controls, 1 experimental genotype) | 5= significantly larger than both other genotypes; 4= larger than both other genotypes; 3=within MAD of at least one genotype; 2= smaller than both other genotypes; 1= significantly smaller than both other genotypes. |
| Wing data (**Fig. S14, S16, S21, S22**) | Mean index 2 (1 control, 2 or more experimental genotypes) | 5= significantly larger than control; 4= larger than control. 3=control or within MAD of control; 2= smaller than control; 1= significantly smaller than control. |
| Wing data (**Fig. 3A, S7, S16, S18**) | Variability index (2 controls, 1 experimental genotype) | 5= significantly more variable than both other genotypes; 4= more variable than both other genotypes; 3= no differences (middle value); 2= less variable than both other genotypes; 1= significantly less variable than both other genotypes. |
| Wing data (**Fig. S14, S21, S22**) | Variability index 2 (1 control, 2 or more experimental genotypes) | 5= significantly more variable than control, 4= more variable than control, 3= control or equal to control, 2= less variable than control, 1= significantly less variable than control. |
| Wing data (**Fig. 3A, S7, S16, S18**) | Canalization index (2 controls, 1 experimental genotype) | 4= more canalized than both other genotypes; 3= no difference (middle value); 2= less canalized than both other genotypes. |
| Wing data (**Fig. S14, S21, S22**) | Canalization index 2 (1 control, 2 or more experimental genotypes) | Canalization index 2 (1 control, 2 or more experimental genotypes): 4= more canalized than control; 3= control or no difference to control; 2= less canalized than control. |
| Eye data (**Fig. 3C (S9)**) | Variability index | 5= significantly more variable than other genotype; 4= more variable than other genotype; 3= no differences (within MAD); 2= less variable than other genotype; 1= significantly less variable than other genotype. |
| Neuropile data (**Fig. 3D (S10)**) | Mean index | 5= significantly larger than both other genotypes; 4= larger than both other genotypes. 3 = 3=within MAD of at least one genotype, 2= smaller than both other genotypes, 1= significantly smaller than both other genotypes |
| Neuropile data (**Fig. 3D (S10)**) | Variability index | 5= significantly more variable than both other genotypes; 4= more variable than both other genotypes; 3= no differences (middle value); 2= less variable than both other genotypes; 1= significantly less variable than both other genotypes. |
| Neuropile data (**Fig. 3D (S10)**) | Canalization index | Canalization index: 4= more canalized than both other genotypes; 3= no difference (middle value); 2 less canalized than both other genotypes. |
| Serotonin data (**Fig. 3E (S11)**) | Variability index | Variability index: 5= significantly more variable than other genotype; 4= more variable than other genotype; 3= no differences (within MAD); 2= less variable than other genotype; 1= significantly less variable than other genotype. |

